# RNA virus polymerase-helicase coupling enables rapid elongation through duplex RNA

**DOI:** 10.1101/2025.03.05.641625

**Authors:** Pim P. B. America, Subhas C. Bera, Arnab Das, Thomas K. Anderson, John C. Marecki, Flávia S. Papini, Jamie J. Arnold, Robert N. Kirchdoerfer, Craig E. Cameron, Kevin D. Raney, Martin Depken, David Dulin

## Abstract

Positive-sense (+) RNA viruses often encode helicases presumed to support replication. Their precise role remains unresolved though, especially in coronaviruses (CoV) where the helicase translocates in the opposite direction to the polymerase. Using high-throughput single-molecule magnetic tweezers, we show that the coronavirus helicase enhances RNA synthesis through duplex RNA by tenfold, forming a directional complex with the viral polymerase. Despite opposing polarity, the helicase coordinates elongation by engaging the non-template strand. A detailed kinetic model derived from large datasets reveals distinct dynamic states, including fast bursting and slow, backtracking-prone modes, which are governed by helicase engagement. These results uncover an active coupling mechanism that modulates replication dynamics and provide a mechanistic basis for continuous versus discontinuous RNA synthesis in coronaviruses. Our findings establish the viral helicase as a central regulator of RNA replication rather than a passive accessory enzyme.

## Introduction

(+)RNA viruses have a single-stranded RNA genome that directly encodes for the viral proteins, such as a helicase (*1, 2*). These helicases are monomeric or hexameric, and hydrolyze NTP and some have been shown to translocate on single-stranded (ss) RNA and displace double-stranded (ds) RNA on their own (*3*). They have been proposed to support viral RNA synthesis by associating with the viral RNA-dependent RNA polymerase (RdRp), though this hypothesis has never been directly demonstrated (*3*). In some (+)RNA virus families, such as alphavirus (e.g. chikungunya) and coronavirus (e.g. SARS-CoV-1, SARS-CoV-2, MERS-CoV), the helicase even translocates in the opposite direction of the RdRp. This raises questions regarding how these helicases assist their RdRp during replication. While such polymerase-helicase complexes have been structurally resolved (*4-6*), there is a lack of functional studies reporting their precise role, and specifically whether they assist replication.

We investigated the replication-transcription complex (RTC) from SARS-CoV-2, the causative agent of the COVID19 pandemic. Extensive structural studies have established that the RTC is composed of a core made of the non-structural protein (nsp) 12-polymerase, associated with the co-factors nsp7 and nsp8 in a 1:1:2 stoichiometry (**Fig. 1A**) (*7-9*). This core RTC may bind either one (nsp13.1) or two nsp13-helicases (nsp13.1 and nsp13.2) (**Fig. 1A**) (*4, 10, 11*). Nsp13-helicase has the typical structure of the super family 1B helicases, where the single-stranded nucleic acid is sandwiched between the 1B and stalk domains on one side and the two RecA domains (RecA1 and RecA2) on the other side (**Fig. 1A**) (*12*). Single-molecule studies have reported rapid and processive dsRNA unwinding by nsp13-helicase when mechanically assisted (*13, 14*). Cryo-electron microscopy (EM) studies have reported that only nsp13.1 interacts with the template strand (**Fig. 1A**) (*4, 10*). Due to the opposite polarity between nsp13-helicase and nsp12-polymerase, nsp13.1 is expected to push the RdRp backward, inducing RdRp backtracking. The backtracked state has been proposed as an intermediate for sub-genomic RNA synthesis, viral RNA recombination, and proofreading (*4*). However, no specific function has been attributed to nsp13.2, apart from allosterically controlling the productive engagement of nsp13.1 with the template RNA (*15*). Downstream duplex RNA represents a mechanical barrier to the elongating core RTC, resulting in an increased probability of long-lived pauses and very slow overall RNA synthesis (*16*). This suggests that one of the two nsp13-helicases may be actively supporting the CoV RTC during replication of the heavily structured coronavirus genomic RNA (*17*), although this has never been demonstrated.

**Fig. 1.**
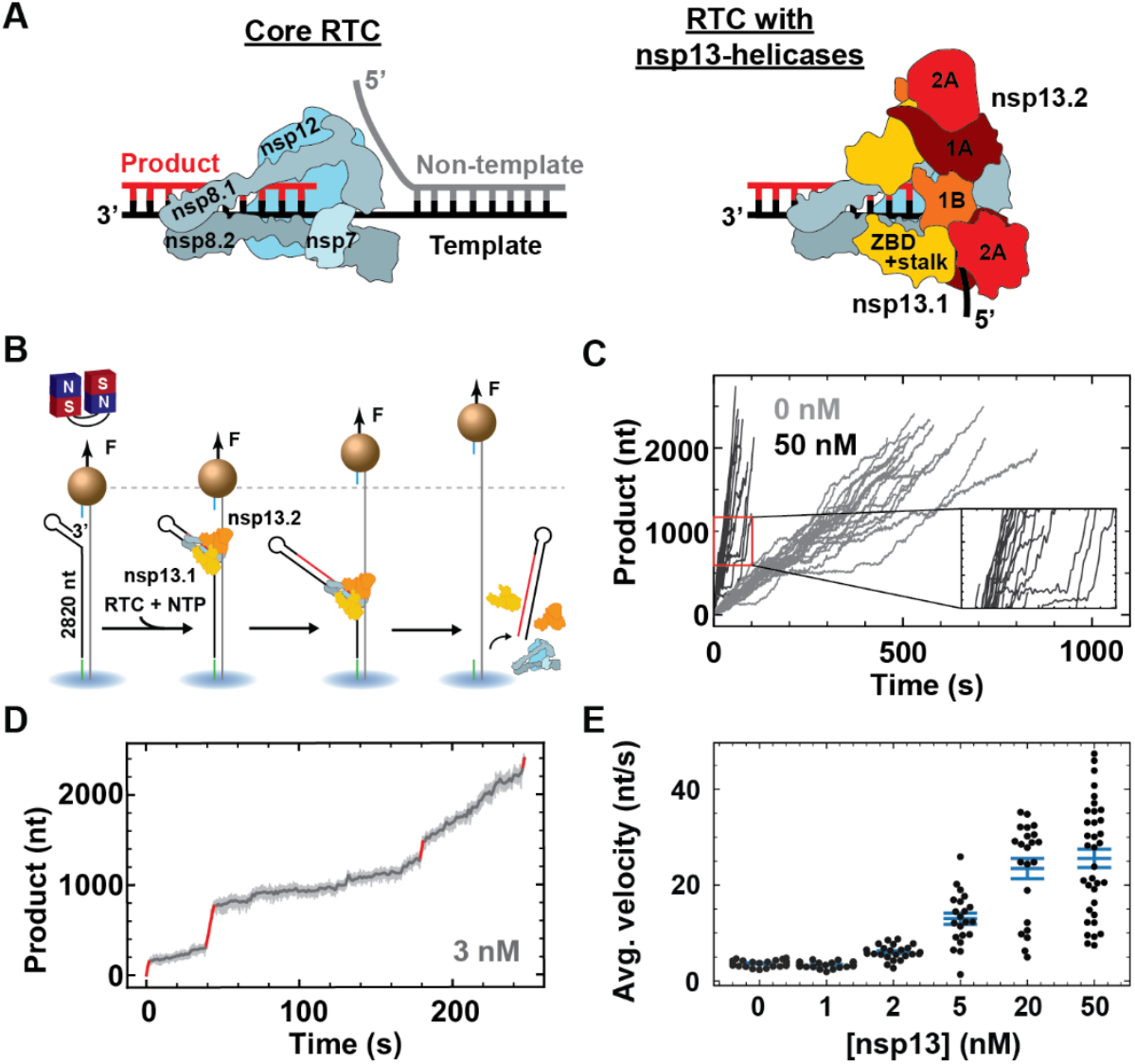
The SARS-CoV-2 nsp13-helicase enables fast RTC elongation on a dsRNA template. **(A)** Core RTC (left) and RTC with two nsp13-helicases (right). Core RTC subunits and nsp13-helicase domains are indicated. **(B)** Schematic of the magnetic tweezers assay to monitor the elongation on dsRNA of the RTC in complex with nsp13-helicase. **(C)** Elongation traces obtained without (grey) or with (black) 50 nM nsp13-helicase. Inset: zoom-in of the traces obtained at saturating nsp13-helicase concentration. Slow elongation dynamics intervals interrupt long bursts of very fast nucleotide addition. **(D)** RTC elongation trace obtained at 3 nM nsp13-helicase (raw data: grey; filtered data: dark grey). The bursts of very fast nucleotide addition are highlighted in red in the filtered trace. **(E)** Average elongation velocity versus nsp13-helicase concentration from traces longer than 1000 nt (dots). Mean values and error bars (SEM) are indicated in blue.

We employed high-throughput magnetic tweezers to investigate how nsp13-helicase associates with the CoV RTC and impacts the dynamics of RNA synthesis. We show that nsp13-helicase specifically associates with the elongating core RTC and not with the polymerase in solution. Furthermore, of the two associated helicases, nsp13.2 enables rapid RNA synthesis through long dsRNA by translocating on the non-template strand. We show that three different complexes, populated as a function of nsp13-helicase concentration, are capable of RNA synthesis and co-exist at equilibrium: the core RTC alone, the core RTC associated with nsp13.1 and the core RTC associated with nsp13.1 and nsp13.2. We modelled that in complex with the RTC, nsp13.2 can either be non-engaged or engaged with the non-template RNA, and only the engaged mode enables very fast RTC elongation. Mechanochemical analysis shows that nsp13.2 also increases the elongation rate by allostery when binding the RTC. Our work establishes a quantitative link between CoV RTC assembly and function. Specifically, we provide a functional demonstration of a novel role for (+)RNA virus helicases in supporting polymerase elongation through structured RNA.

## Results

### Nsp13-helicase promotes rapid RNA synthesis on a dsRNA template by the CoV RTC

We employed a high-throughput magnetic tweezers assay to monitor CoV RTC elongation on dsRNA, as previously described for poliovirus (PV) RdRp (*18, 19*). Briefly, a pair of permanent magnets located above a flow chamber is used to apply an attractive force to the magnetic beads. The force stretches the dsRNA tether that attaches each bead to the flow chamber glass surface (**Fig. 1A, Materials and Methods**) (*20*). After flushing the reaction buffer containing the viral proteins and NTPs (**Materials and Methods**), the RTC assembles on a small hairpin terminating the template strand 3’-end. Once assembled, the RTC elongates through the dsRNA until reaching the end of the 2820 nt long template strand (**Fig. 1B**). The RTC elongation activity converts the dsRNA tether into ssRNA by displacing the template from the non-template strand, resulting in an increase of the tether extension (**Fig. 1B**) (*21*). The force was kept constant throughout the experiment by maintaining the magnets at a constant height above the flow chamber (*22, 23*). In absence of nsp13-helicase, we found that the core RTC elongates slowly on dsRNA, with an average velocity of (3.6 ± 0.1) nt/s (**Fig. 1CE**). Adding 50 nM of nsp13-helicase to the reaction buffer dramatically increases the average elongation velocity to (25.6 ± 1.9) nt/s (**Fig. 1CE**). This shows that nsp13-helicase supports RNA synthesis through duplex RNA. Decreasing nsp13-helicase concentration to 3 nM, we can distinguish very fast nucleotide addition bursts (∼30 nt/s) of tens to hundreds of nucleotides long that are interrupted by slow nucleotide addition cycles (**Fig. 1E**). The average elongation velocity increases when the concentration of nsp13-helicase is varied from 0 to 50 nM, saturating above 10 nM (**Fig. 1E**).

### Nsp13-helicases specifically associate with the CoV core RTC and nsp13.2 translocates on the non-template RNA strand

To investigate whether nsp13-helicase associates specifically with the core RTC to increase the RNA synthesis rate, we replaced the core RTC by the poliovirus RdRp and tracked RNA synthesis on the same dsRNA template. As poliovirus RdRp has a different structure from the CoV core RTC (*8*), we expected only nonspecific interactions between poliovirus RdRp and nsp13-helicase. Looking at poliovirus RdRp elongation dynamics with and without nsp13-helicase, we observe no significant differences in the average elongation velocities of (18.7 ± 0.5) nt/s and (19.2 ± 0.7) nt/s (**Fig. 2A, Fig. S3ACD**). To further interrogate the specificity of nsp13-helicase association with the RTC, we monitored the core RTC elongation dynamics in the presence of the ATPase-dead mutant K288A nsp13-helicase (**Fig. 2B**). A significant reduction of the RTC average elongation velocity relative to the core RTC alone was noticeable, from (3.6 ± 0.1) nt/s to (1.1 ± 0.1) nt/s.

**Fig. 2.**
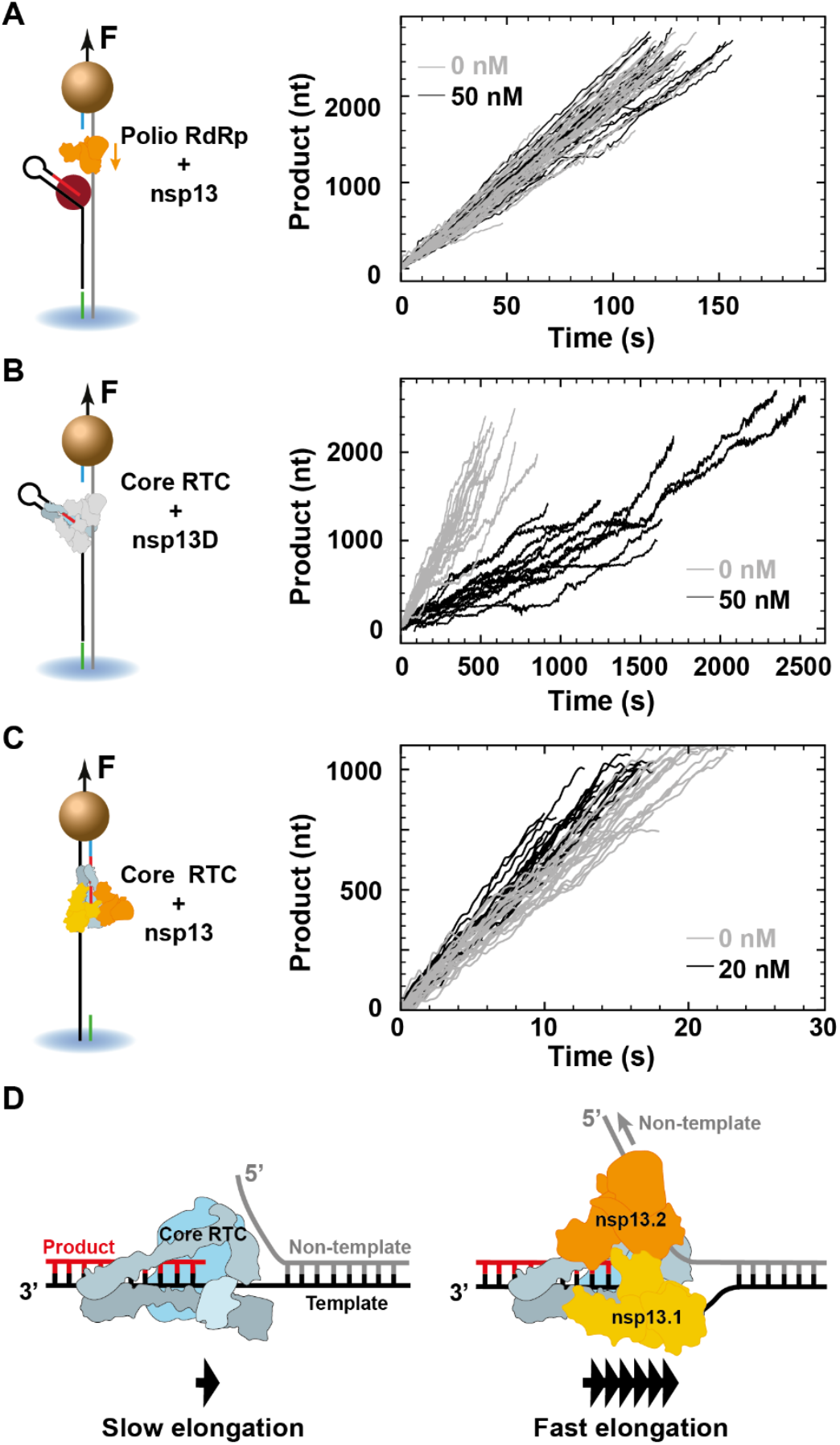
Nsp13-helicase forms a specific complex with the SARS-CoV-2 core RTC and nsp13.2 supports the elongating polymerase by translocating on the non-template strand. **(A, B, C)** On the left, schematics describing the measurements performed with nsp13-helicase using either (A) poliovirus RdRp or (B, C) SARS-CoV-2 core RTC, elongating on either a dsRNA (A, B) or ssRNA (C) template, with either a wild-type nsp13-helicase (A, C) or an ATPase dead K288A nsp13-helicase mutant (B). The corresponding elongation activity traces are shown on the right, i.e. without (grey) or with (black) nsp13-helicase at the indicated concentration. **(D)** Schematic representation resuming the observations. The core RTC elongates slowly against duplex RNA. Once associated with the two helicases, nsp13.2 binds to the non-template RNA and utilizes ATP hydrolysis to translocate 5’ to 3’ and enable fast RTC elongation.

We then employed a ssRNA template to investigate whether nsp13-helicase still affected the elongation dynamics of the core RTC. Nsp13-helicase translocates very fast from 5’ to 3’ (∼1 kb/s) on ssRNA (*24*), while the elongating core RTC is much slower (∼50 nt/s) (*16*). The difference in polarity between helicase and RTC suggests that head-to-head collisions may destabilize the RTC and/or slow down elongation. To test this hypothesis, we employed a ∼1 kb long ssRNA template to monitor the RTC primer-extension dynamics (*16, 25*) at increasing concentrations of nsp13-helicase (**Fig. 2C**). Surprisingly, when comparing the cases with or without 20 nM nsp13-helicase we observed a stable product length ((951 ± 20) nt vs. (972 ± 29) nt), and an increase in the RTC average elongation velocity ((62.4 ± 1.2) nt/s vs. (52.1 ± 1.6) nt/s) (**Fig. 2C**). This indicates that nsp13-helicase allosterically increases the elongation rate after binding the core RTC. If collisions between translocating nsp13-helicase and RTC occurred, they did not significantly impact the elongation of the latter.

Cryo-EM studies found that either nsp13.1 or nsp13.1 and nsp13.2 are associated with the core RTC, but never with nsp13.2 alone (*4, 10*). This observation suggests a stepwise assembly process, where nsp13.1 binds the core RTC first, followed by nsp13.2. Furthermore, it was shown that nsp13.1 associates with the template RNA (*4, 10*). Our data shows rapid RNA synthesis on dsRNA when the RTC is associated with an ATPase active nsp13-helicase. Using an ATPase dead mutant instead resulted in slower elongation dynamics. Altogether, we conclude that nsp13.2 accelerates RNA synthesis through duplex RNA by specifically associating to the RTC and translocating on the non-template strand (**Fig. 2D**).

In the following sections we employ statistical analysis and kinetic modeling to establish the nucleotide addition pathways and the energy landscape of the RTC-nsp13 assembly (**Fig. 3**). In addition, our kinetic modeling captures how nsp13.2 binding increases RNA synthesis velocity (**Fig. 4**).

**Fig. 3.**
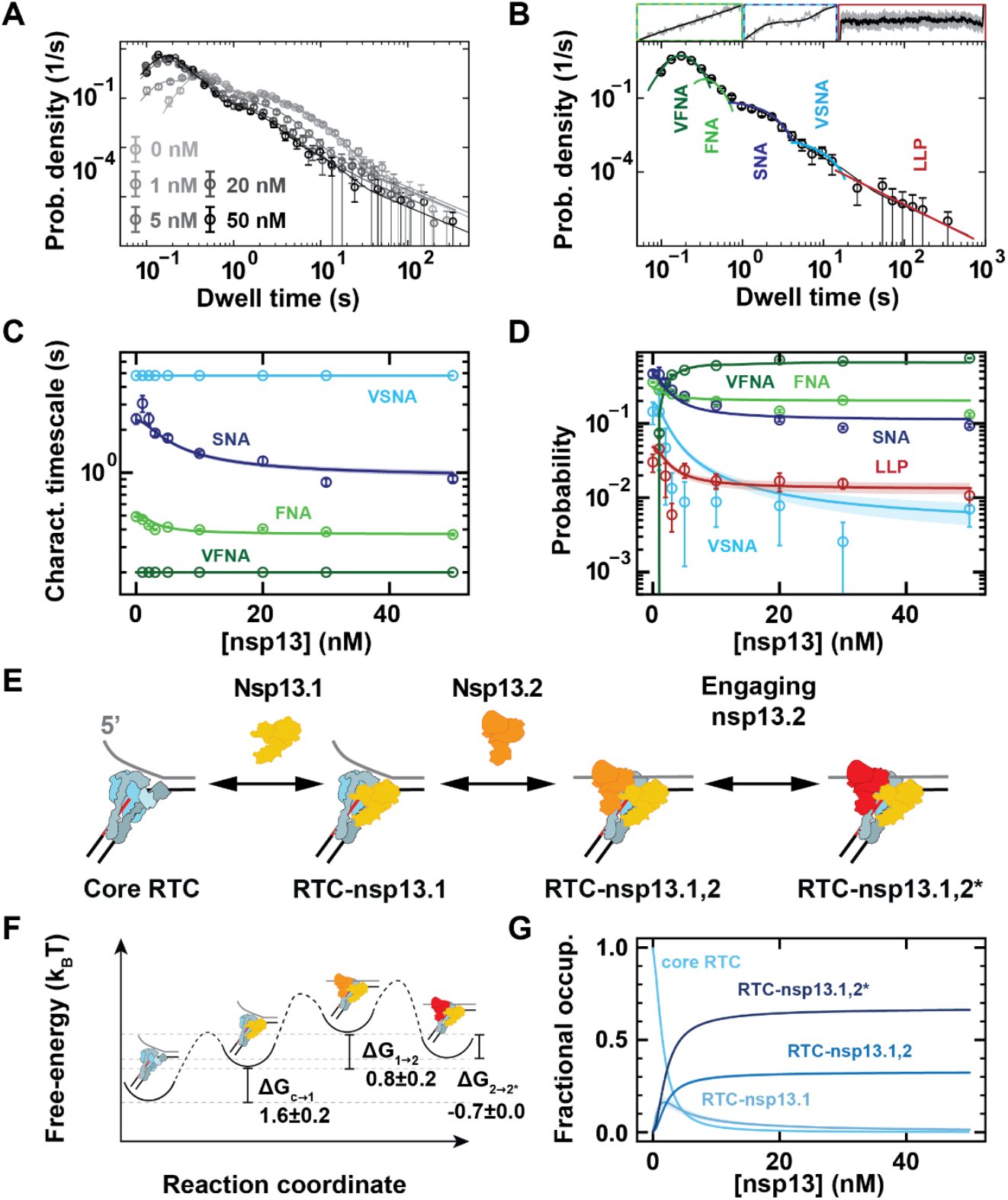
Assembly and stability of the SARS-CoV-2 polymerase-helicase complex. **(A)** Dwell-time distributions extracted from RTC elongation traces obtained at the indicated nsp13-helicase concentrations (circles) and their respective dwell-time fits (solid lines). **(B)** The dwell-time fit-function consists of five probability density functions (pdf’s): very fast, fast, slow and very slow nucleotide addition (VFNA, FNA, SNA, VSNA) pathways and long-lived pauses (LLP). These pdf’s are combined and fitted (black line) on the dwell-time distribution (circles). The insets above are snapshots of representative parts of the traces. The error bars in (A, B) represent one standard deviation from 100 dwell-time resamples. **(C, D)** Characteristic timescales and probabilities of the different NA pathways and LLP state versus nsp13-helicase concentration. The circles with error bars and the solid lines with shaded areas represent the mean values and one standard deviation from 100 bootstraps of the individual fits and the RTC assembly - Elongation dynamics model fits, respectively. VFNA and VSNA characteristic timescales were fixed to 0.2 s and 4.8 s. **(E)** Schematic of the RTC assembly model. **(F)** The extracted free-energy landscape and **(G)** corresponding fractional occupancies versus nsp13-helicase concentration fitted on the dwell-time distributions with shaded areas as error bars as in (C, D).

**Fig. 4.**
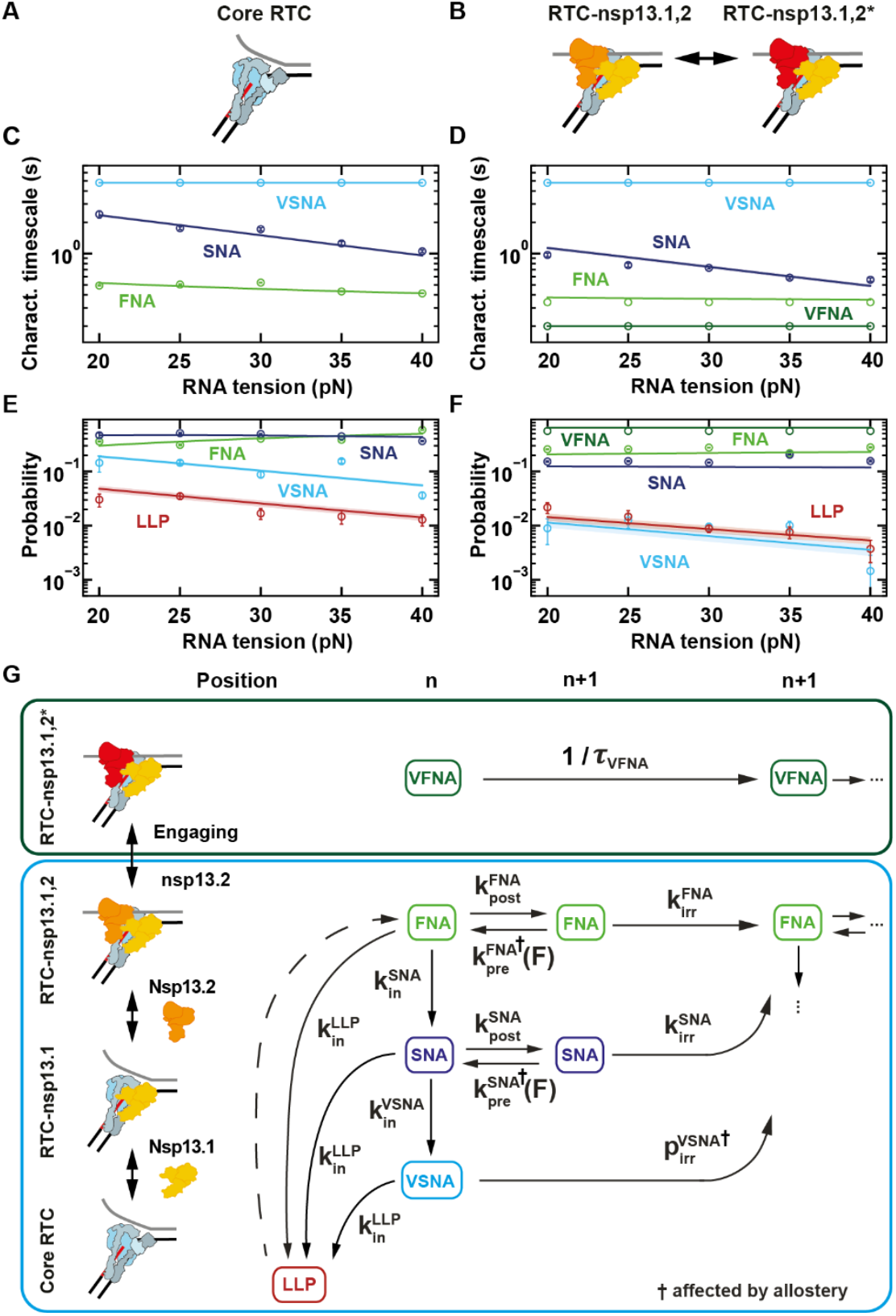
Mechanochemical reaction scheme for the nucleotide addition cycle of the SARS-CoV-2 core RTC alone or in complex with nsp13-helicase. Schematic representation of the different elongating RTCs **(A)** without and **(B)** with nsp13-helicase at saturating concentration. **(C, D)** Characteristic timescales and **(E, F)** Probabilities for the different nucleotide addition pathways (VFNA, FNA, SNA, VSNA) and long-lived pause (LLP) state versus tension applied to the non-template RNA strand and either (C, E) without or (D, F) with saturating nsp13-helicase concentration. The circles with error bars represent the mean value and one standard deviation from 100 bootstraps of the individual dwell-time fits. The solid lines and shaded areas represent the mean values and the region within one standard deviation of the mean obtained from 100 bootstraps of the global fit with the mechanochemical reaction scheme shown in (G) (**Materials and Methods**). **(G)** Mechanochemical reaction scheme underlying the timescales and probabilities (green box) for the VFNA pathway, which is only entered by the RTC-nsp13.1,2* complex, and (blue box) for the nucleotide addition pathways entered by the core RTC, RTC-nsp13.1 and RTC-nsp13.1,2. The solid and dashed arrows represent kinetic steps for which rates could either be extracted or not, respectively, from the global fit.

### Nsp13-helicase modifies the kinetic fingerprint of the RTC elongation traces

We analyzed the RTC elongation dynamics on a dsRNA template as a function of nsp13-helicase concentration (**Fig. 1C-E**). We scanned the elongation traces with a non-overlapping window of ten nucleotides and measured the duration of ten consecutive nucleotide addition cycles, which we coined dwell times (*21, 26*). We previously showed that there are four kinds of events in the core RTC elongation dynamics that can dominate the dwell-time window (*16*). The shortest dwell times (less than ∼1 s) are described by a gamma distribution representing the ten uninterrupted fast nucleotide addition (FNA) events. Intermediate duration dwell times (from 1 to 20 s) are described by two exponential distributions, corresponding to the slow (SNA) and the very slow (VSNA) nucleotide addition pathways (*16*). A power-law distribution consistent with a *t*^−3/2^ decay describes the dwell-times longer than ∼20 s and are referred to as long-lived pauses (LLP) (**Fig. 3AB, Supplementary Information Section S1**). In the presence of nsp13-helicase we observe very fast nucleotide addition (VFNA) bursts in the traces (**Fig. 1D**), resulting in a new short-time gamma peak in the dwell-time distribution (**Fig. 3AB**).

The different distributions are combined into a single fit-function and their characteristic timescales and probabilities are extracted using maximum-likelihood estimation (MLE) (**Fig. 3AB, Fig. S8, Materials and Methods, Supplementary Information Section S1**) (*26*).

### Connection of the RTC-nsp13 complex assembly model with RTC elongation dynamics

We now set out to discern the assembly of the RTC-nsp13 complex from the features captured in the traces and the dwell time distributions. We noticed that the characteristic timescale of the VFNA cycles was independent of nsp13-helicase concentration (**Fig. 3D, Fig. S2AI**) and that the VFNA bursts always lasted tens to hundreds of nucleotide addition cycles (**Fig. 1D**). These observations suggest that single stable complexes are responsible for the VFNA bursts over multiple dwell-time windows.

The dwell-time distributions representing the RTC elongation dynamics on dsRNA do not change when increasing nsp13-helicase concentrations above 10 nM (**Fig. 3B**). This indicates that the two nsp13-helicase binding sites on the RTC are occupied at these concentrations (*4, 10*). However, above 10 nM nsp13-helicase, we still observe slow kinetic events in the traces (**Fig. 1C**), consistent with entering the FNA, SNA and VSNA pathways, and the LLP state (**Fig. 3B**). We conclude that a single RTC-nsp13 complex with two bound nsp13-helicases can exist in two configurations: one only entering the VFNA pathway and the other entering the slow kinetic events. Structural work has shown that the nsp13.2 conformation allosterically controls nsp13.1 engagement with the RNA template strand (*15*). We propose that the nsp13.1 conformation can similarly control nsp13.2 engagement with the non-template RNA and entry into the VFNA pathway. Based on these findings, we consider an RTC dynamic equilibrium model for several complexes and states: nsp13.1 in association with the core RTC (RTC-nsp13.1); nsp13.2 in association with the RTC-nsp13.1 and not engaged with the non-template strand (RTC-nsp13.1,2); nsp13.2 in association with the RTC-nsp13.1 and engaged with the non-template strand (RTC-nsp13.1,2*). The latter configuration results in the VFNA bursts (**Fig. 3E, Fig. S4, Fig S6, Supplementary Information Section S2**).

Assuming that RTC-nsp13.1 shows the same elongation dynamics as the core RTC, we use the RTC dynamic equilibrium model to globally fit the dwell-time distributions at all measured nsp13-helicase concentrations (solid lines, **Fig. 3CD, Fig. S10**). From this fit, we extract the free energy differences between each RTC state (**Fig. 3F**): (1.6 ± 0.2) k_B_T for core RTC to RTC-nsp13.1, (0.8 ± 0.2) k_B_T for RTC-nsp13.1 to RTC-nsp13.1,2 and (−0.7 ± 0.0) k_B_T for RTC-nsp13.1,2 to RTC-nsp13.1,2*. From these free energy differences, we obtained the equilibrium association constants *K*_a_ for the binding of nsp13.1 and nsp13.2 to the RTC, i.e. (0.22±0.05) nM^-1^ and (0.47±0.08) nM^-1^. We also derived the fractional occupancy of each complex at 20 nM nsp13-helicase (saturating): (0.64±0.01) for RTC-nsp13.1,2*, (0.31±0.01) for RTC-nsp13.1,2, (0.034±0.005) for RTC-nsp13.1 and (0.0081±0.0007) for the core RTC (**Fig. 3G, Materials and Methods**).

Interestingly, the saturating nsp13-helicase concentration for two bound helicases to the core RTC (∼10 nM, **Fig. 3G**) is much lower than the saturating nsp12-polymerase concentration for binding to the RNA (∼200 nM) (*16, 27*), indicating that nsp13-helicase binds the elongating core RTC with a much higher affinity than it binds freely diffusing nsp12-polymerase or core RTC. This likely originates from nsp13.1 and nsp13.2 forming interactions with the N-terminal poles of nsp8.2 and nsp8.1 (**Fig. 1A**) (*4, 10*). Indeed, these poles are very flexible in absence of duplex RNA (*8, 28*), which may hinder nsp13-helicase association.

### The CoV RTC nucleotide addition cycle

We now build a mechanochemical reaction scheme that captures the non-bursting elongation dynamics of the different RTC complexes and use the dependence on RNA tension to extract how the NA pathways and LLP state are connected. To this end, we consider that the polymerase nucleotide addition cycle starts from the pre-translocated state and consists of several successive steps: translocation, NTP binding, catalysis, and pyrophosphate release (**Fig. S7A**) (*16*).

We monitored the RTC elongation dynamics on a dsRNA template as a function of the applied tension, either for the core RTC alone (**Fig. 4ACE**) or at saturating nsp13-helicase concentration (**Fig. 4BDF**). The characteristic timescale of the VFNA bursts shows no significant trend with tension (**Fig. 4DF**) so we can only conclude that it is not rate-limited by translocation. The non-bursting NA pathways show a dependence on tension (**Fig. 4C-F**), allowing us to establish how they are connected.

For a dsRNA template, the downstream dsRNA fork opposes RTC translocation as an energy barrier that is modulated by force (*19, 21*). In our previous work, we showed that the back-and-forth translocation process is not equilibrated, while NTPs are at saturating concentration and therefore the NTP binding step is equilibrated (*16*). Furthermore, we found that only the backward translocation in the FNA pathway is affected by the force applied on the template (**Fig. S7A**) (*16*). Finally, we assume that all non-bursting NA pathways can be similarly modelled.

From the fits to the dwell-time distributions (circles, **Fig. 4C-F**), we see that the FNA characteristic timescale of the core RTC decreases with increasing RNA tension, confirming that the tension biases the RTC towards the FNA post-translocated state. Furthermore, the FNA probability increases with tension, while both the SNA and VSNA probabilities decrease (circles, **Fig. 4E**). This tension dependence suggests that the slower NA pathways originate from the FNA pre-translocated state (**Fig. S7B**), as we previously established for the core RTC (*16*). With increasing tension, the VSNA probability decreases with respect to the SNA probability, indicating that the VSNA is entered from the SNA pre-translocated state (**Fig. 4EF**). The ratio of the LLP and VSNA probability is constant with tension (**Fig. 4EF**), suggesting that the LLP state is also predominantly entered from the SNA pre-translocated state. However, the ratio of the LLP and VSNA probability is different between the core RTC and RTC-nsp13.1,2 (**Fig. 4EF**). To allow the model to capture this shift, we assume the LLP can be entered from the pre-translocated state of each NA pathway (**Fig. 4G, Supplementary Information Section S3**).

In summary, we model the non-bursting nucleotide addition pathways as starting from the pre-translocated state of the FNA pathway. From there, the RTC can either directly incorporate a nucleotide, enter the SNA pre-translocated state or enter the LLP. The RTC can then either incorporate a nucleotide via the SNA pathway, enter the VSNA pre-translocated state and incorporate a nucleotide, or enter the LLP state. Once the LLP state is entered, we cannot establish from which NA pathway the RTC eventually exits, since the dwell-time is dominated by the pause duration.

### Allosteric effect from nsp13.2 association on the RTC-nsp13.1,2 nucleotide addition cycle

Interestingly, when the RTC elongates on a dsRNA template, we find that the FNA and SNA characteristic timescales significantly decrease with nsp13-helicase concentration, saturating above 10 nM (**Fig. 3C**). The similar saturation for the VFNA pathway indicates that RTC-nsp13.1,2 elongation dynamics is also affected. On an ssRNA template, we find that the FNA characteristic timescale decreases significantly when either active or ATPase-dead nsp13-helicase is added at saturating concentration (**Fig. S5**), indicating that the decrease in timescales is the result from an allosteric effect of nsp13 association. The decrease in FNA timescale is still apparent at saturating nsp13-helicase concentration, indicating that at least the association of nsp13.2 impacts the elongation velocity.

From these findings we infer that the elongation dynamics of the core RTC and RTC-nsp13.1 can be assumed indistinguishable, while RTC-nsp13.1,2 employs the same pathways, but with different timescales and probabilities due to allostery of nsp13.2 association. The VFNA pathway is only entered from the nsp13.2-engaged state (RTC-nsp13.1,2*) (**Fig. S6**). We confirm this model by connecting the RTC-nsp13 assembly with RTC elongation dynamics and global fitting to the dwell-time distributions with nsp13-concentration for elongation dynamics on the dsRNA template (**Fig. 3A, Fig. S10-S13, Table S2, Materials and Methods, Supplementary Information Section S2**).

We now investigate how the allostery of nsp13.2 association to the RTC can be described on the mechanochemical level. Increasing RNA tension has no significant effect on the FNA characteristic timescale and probability for RTC-nsp13.1,2 (**Fig. 4DF**). There is only an RNA tension dependency on the backward translocation rates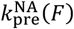in our model for the FNA and SNA pathways (**Fig. 4G**). Therefore, the force independence of the FNA characteristic timescale and probability can be captured by the FNA backward translocation rate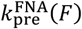becoming negligible compared to the effective nucleotide incorporation rate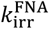upon nsp13.2 association (**Table S3**). In contrast, the SNA characteristic timescale still decreases with tension for RTC-nsp13.1,2 (**Fig. 4CD**), indicating that SNA backward translocation with rate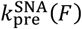is not completely abolished in the SNA pathway by nsp13.2 association. Furthermore, at the same tension, the ratio of VSNA to SNA probabilities is smaller for the core RTC than for the RTC-nsp13.1,2 (**Fig. 3D, Fig. 4EF**). This effect is captured by a decreased probability for nucleotide incorporation in the VSNA pathway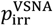for the RTC-nsp13.1,2 complex (**Fig. 4G, Table S3, Supplementary Information Section S3**).

From this model we conclude that nsp13.2 association to the RTC makes backward translocation negligible in the FNA pathway by allostery. Our model suggests that the binding of nsp13.2 also reduces the backward translocation rate in the SNA pathway and decreases the probability for nucleotide incorporation in the VSNA pathway.

### A unified model captures all the data

Combining RTC assembly (**Supplementary Information Section S2.1**), elongation dynamics (**Section S2.2**), and the mechanochemical reaction scheme (**Section S3, Fig. S1**), we capture the complete dependence of the dwell-time distributions on the nsp13-helicase concentration and tension through a global fit (**Fig. 3CD, Fig. 4C-F, Fig. S14, Materials and Methods**). Our ability to quantitively capture all dependencies and distributions with a unified model lends confidence to our reaction scheme (**Fig. 4G**) and allows us to extract its rates and the free-energy differences controlling the RTC Assembly (**Table S3, Supplementary Information Section S3**).

## Discussion

While nsp13-helicase has been proposed to support RNA synthesis (*11*), this function remained to be shown. Here, we employed high-throughput, single-molecule, magnetic tweezers to show that nsp13-helicase specifically associates with the elongating core RTC. Furthermore, nsp13.2 supports RTC elongation on a dsRNA template by translocating on the non-template strand (**Fig. 1, Fig. 2**). These results solve the conundrum of nsp13-helicase supporting nsp12-polymerase translocation while having opposite polarities. Our large statistics in many conditions enabled us to reveal the mechanism of assembly of the CoV RTC with nsp13-helicase through extensive statistical analysis and kinetic modeling. Namely, two nsp13-helicases associate sequentially to the core RTC, with nsp13.1 binding first, followed by nsp13.2 (**Fig. 2, Fig. 3**). Moreover, our results support a model where nsp13.2 alternates between a non-engaged and an engaged state with the non-template RNA. We showed that at saturating concentration of nsp13-helicase, ∼31% and ∼65% of the incorporated nucleotides respectively originate from an RTC associated with either a non-engaged or an engaged nsp13.2 **(Fig. 3**). Finally, we derived a unified model describing both RTC assembly and the nucleotide addition mechanochemistry of the RTC complexes **(Fig. 4**).

Our data provide unequivocal evidence that nsp13-helicase directly enhances the RTC by enabling efficient elongation through structured RNA. Cryo-EM tomography has revealed the presence of dsRNA within the replication organelles of infected cells (*29, 30*), and duplex RNA has been suggested to serve as intermediates in both viral replication and transcription (*31*). The existence of such intermediates would substantiate a functional role for nsp13-helicase in supporting polymerase progression. However, even if dsRNA intermediates are not required during CoV replication, the highly structured nature of the viral genome (*17*) alone would likely necessitate helicase assistance to sustain rapid and processive RTC elongation.

Nsp13.2 has been shown to allosterically control the productive engagement of nsp13.1 with the RNA template strand (*15*). Similarly, our data supports a model where nsp13.1 allosterically controls nsp13.2 productive engagement with the non-template RNA. We propose that this mechanism ensures that only one helicase at a time is actively engaged with its associated RNA strand, preventing antagonistic activities of the two helicases. Our results suggest the VFNA pathway is only populated by RTC-nsp13.1,2*, and nsp13.2 must be not engaged with the non-template strand for the RTC to enter a long-lived pause, including polymerase backtracking (*32*).

The RTC bound with two nsp13-helicases stochastically alternates between the two elongation states RTC-nsp13.1,2 and RTC-nsp13.1,2*. This state-switching provides a striking example of dynamic heterogeneity (*33*), and may regulate nsp13.1 productive engagement with the template RNA. Nsp13.1 has been proposed to stimulate RTC backtracking, to promote either polymerase template switching (*34*) or proofreading by the 3’-to-5’ exonuclease nsp14 (*4*), though these functions remain to be demonstrated (*11*). This ability to stochastically switch between nsp13.1 and nsp13.2 productive engagement may enable a high level of regulation through sensing sequence-dependent context during replication. This could help detecting transcription regulatory sequences to regulate the fraction of complexes performing continuous versus discontinuous replication, or to sense nucleotide mismatch incorporation to enable proofreading nsp14.

We anticipate that our results will impact our understanding of the mechanism of action of antiviral nucleotide analogs targeting the RTC such as the FDA-approved remdesivir (*35*). Indeed, remdesivir incorporation by nsp12-polymerase has been shown to induce long-lived, polymerase backtrack-related pauses during elongation (*25, 36-39*). As nsp13.2 assists the RTC to elongate through dsRNA, it is tempting to think that nsp13-helicase may help the RTC to overcome the pause induced by remdesivir incorporation. Whether the mechanism of action of antiviral nucleoside analogs targeting the CoV RTC should be revised when evaluated in the presence of additional viral co-factors associating with the core RTC (nsp13-helicase, nsp14-Exonuclease) will require future investigations.

Our approach combining high-throughput single-molecule biophysics and kinetic modeling is uniquely capable of reconstituting an RNA synthesis-competent extended CoV RTC, complementing our structural knowledge with function. This study paves the way towards the investigation of other positive-sense RNA virus RTCs and reveal whether the function discovered here for CoV nsp13-helicase is conserved for other viral helicases.

## Authors contribution

DD, KDR, CEC and JJA designed the research. DD and SCB designed the experiments. SCB and AD performed the experiments. SCB and PPBA analyzed the data. FSP made the RNA construct used for the study. TKA and RNK provided SARS-CoV-2 core RTC proteins. JJA and CEC provided poliovirus proteins. KDR and JCM provided the coronavirus nsp13-helicase. PPBA and MD developed the kinetic model. PPBA, MD and DD interpreted the results. PPBA, MD and DD wrote the article. DD supervised the research. All authors have edited the manuscript.

## Data and materials availability

The data and analysis scripts used for every figure and table in this manuscript will be made available as soon as the manuscript will be accepted.

## Supplementary Materials

Materials and Methods Supplementary Text Figs. S1 to S14

Tables S1 and S3

