## Supplementary Materials for "RNA virus polymerase-helicase coupling enables rapid elongation through duplex RNA"

**The PDF file includes:**

Materials and Methods

Supplementary Text

Figs. S1 to S14

Tables S1 to S3

**Fig. S1** Schematic of the build up of the complete model describing elongation dynamics by the core RTC and RTC-nsp13 complexes on dsRNA.

**Fig. S2** Parameters from the dwell-time fits with all parameters free or the VFNA and VSNA timescales fixed for RTC elongation dynamics with nsp13-helicase on a dsRNA template.

**Fig. S3** Nsp13-helicase specifically associates with the SARS-CoV-2 core RTC and nsp13.2 assists elongation dynamics by engaging with the non-template RNA strand.

**Fig. S4** Comparison of RTC-nsp13 assembly models shows that nsp13.2 engages with the non-template RNA strand.

**Fig. S5** Comparison of RTC elongation dynamics on a ssRNA template.

**Fig. S6** Model connecting the RTC assembly with elongation dynamics on dsRNA.

**Fig. S7** Simplification and connections made in the process of building the mechanochemical model for the SARS-CoV-2 RTC elongation dynamics on a dsRNA template.

**Fig. S8** Dwell-time fits with all parameters free for RTC elongation dynamics on a dsRNA template with nsp13-helicase.

**Fig. S9** Dwell-time fits with the VFNA and VSNA characteristic timescale fixed for RTC elongation dynamics on a dsRNA template with nsp13-helicase.

**Fig. S10** Global fit of RTC Assembly-Elongation dynamics model with the single nucleotide parameters free for the RTC-nsp13.1,2 complex.

**Fig. S11** Global fit of RTC Assembly-Elongation dynamics model with the single nucleotide probabilities free for the RTC-nsp13.1,2 complex.

**Fig. S12** Global fit of RTC Assembly-Elongation dynamics model with the single nucleotide probabilities and timescales fixed to the core RTC values.

**Fig. S13** Check on timescale averaging with fitted parameters from the RTC Assembly-Elongation dynamics model versus nsp13-helicase concentration.

**Fig. S14** Global fit of mechanochemical reaction scheme for RTC elongation dynamics on a dsRNA template in presence and absence of nsp13-helicase.

**Table S1** Parameters for RTC assembly model fits to the VFNA probabilities versus nsp13-helicase concentration.

**Table S2** Parameter values for the RTC Assembly - Elongation dynamics model.

**Table S3** Global fit parameters for the RTC assembly, elongation dynamics and mechanochemical model.

### Materials and Methods

#### Recombinant Protein Expression of RdRp (nsp12) and cofactors (nsp7 and nsp8) from SARS-CoV-2.

This protocol was described in Ref. (Seifert, Bera et al. 2021). The SARS-CoV-2 nsp12 gene was codon optimized and cloned into pFastBac with C-terminal additions of a TEV site and strep tag (Genscript). The pFastBac plasmid and DH10Bac E. coli (Life Technologies) were used to create recombinant bacmids. The bacmid was transfected into Sf9 cells (Expression Systems) with Cellfectin II (Life Technologies) to generate recombinant baculovirus. The baculovirus was amplified through two passages in Sf9 cells, and then used to infect 1 L of Sf21 cells (Expression Systems) and incubated for 48 hrs at 27°C. Cells were harvested by centrifugation, resuspended in wash buffer (25 mM HEPES pH 7.4, 300 mM NaCl, 1 mM $MgCl_2$, 5 mM DTT) with 143 µl of BioLock per liter of culture. Cells were lysed via microfluidization (Microfluidics). Lysates were cleared by centrifugation and filtration. The protein was purified using StrepTactin Superflow agarose (IBA). StrepTactin eluted protein was further purified by size exclusion chromatography using a Superdex 200 Increase 10/300 column (GE Life Sciences) in 25 mM HEPES, 300 mM NaCl, 100 µM $MgCl_2$, 2 mM TCEP, at pH 7.4. Pure protein was concentrated by ultrafiltration prior to flash freezing in liquid nitrogen. The SARS-CoV-2 nsp7 and nsp8 genes were codon optimized and cloned into pET46 (Novagen) with an N-terminal 6x histidine tag, an enterokinase site, and a TEV protease site. Rosetta2 pLys E. coli cells (Novagen) were used for bacterial expression. Cultures were grown to an OD600 of 0.8 and induced with a final concentration of 0.5 mM isopropyl β-D-1-thiogalactopyranoside (IPTG) and growth temperature was reduced to 16°C for 16 hrs. Cells were harvested by centrifugation and pellets were resuspended in wash buffer (10 mM Tris pH 8.0, 300 mM NaCl, 30 mM imidazole, 2 mM DTT). Cells were lysed via microfluidization and lysates were cleared by centrifugation and filtration. Proteins were purified using Ni-NTA agarose beads (Qiagen) and eluted with wash buffer containing 300 mM imidazole. Eluted proteins were digested with 1% w/w TEV protease during overnight room temperature dialysis (10 mM Tris pH 8.0, 300 mM NaCl, 2 mM DTT). Digested proteins were passed back over Ni-NTA to remove undigested protein before concentrating the proteins by ultrafiltration. Nsp7 and nsp8 proteins were further purified by size exclusion chromatography using a Superdex 200 Increase 10/300 column (GE Life Sciences). Purified proteins were concentrated by ultrafiltration prior to flash freezing with liquid nitrogen.

#### Recombinant Protein Expression of the wild-type and the ATPase dead mutant K288A nsp13-helicases from SARS-CoV-2.

The coding sequence for nsp13-helicase from the SARS CoV-2 Washington isolate (Genbank MN985325) was synthesized as an *E. coli* codon-optimized fragment (GenScript, Piscataway NJ) and cloned into the *Bsa*I site of the pSUMO plasmid (LifeSensors, Malvern, PA) to produce an *N*-terminal six histidine-tagged SUMO-NSP13-HELICASE fusion cassette (6XHis-SUMO-nsp13). Using the manufacturer's recommendations (QuikChange II Site-Directed Mutagenesis Kit, Agilent Technologies), oligonucleotide-directed site-directed mutagenesis was used to mutate the codon in the wild type nsp13-helicase for the critical catalytic lysine in the Walker A motif (K288) to an alanine residue. This change produced the K288A mutant lacking ATPase activity (designated nsp13D). Final wild-type and mutant plasmids were sequence-verified through the UAMS Sequencing Core Facility using a 3130XL Genetic Analyzer (Applied Biosystems, Foster City, CA). The SUMO-nsp13 and the SUMO-nsp13D constructs were transformed into Rosetta2 cells, and colonies were grown overnight at 37°C in NZCYM (Research Products International, Mount Prospect, IL) supplemented with kanamycin (50 μg/ml) and chloramphenicol (25 μg/ml). The cultures were diluted 1:100 into fresh antibiotic-containing NZCYM media and grown to an $OD_{600\ nm}$ of 0.8-1. The bacterial media was supplemented with 0.1 mM $ZnSO_4$ and 0.2% dextrose and cooled on ice for 10 min. Wild type and mutant protein expression was induced with 0.2 mM isopropyl β-D-1-thiogalactopyranoside (IPTG) at 18°C for 12-16 hours. The cells were harvested by centrifugation at 4,000 x *g* for 15 min at 4°C, and pellets stored at -80°C.All purifications steps were carried out on ice or at 4°C. Pellets were resuspended in lysis buffer (50 mM sodium phosphate, pH 8.0, 300 mM NaCl, 1 mM β-mercaptoethanol, 10% glycerol and 20 mM imidazole) supplemented with 2 mM phenylmethylsulfonyl fluoride (PMSF) and 1X EDTA-free protease inhibitor cocktail (Pierce). Bacteria were lysed by microfluidization and the lysate clarified by centrifugation at 17,000 x *g* for 1 hour at 4°C. The His-tagged SUMO-nsp13 was passed through a HisTrap FF column (Cytiva) equilibrated in lysis buffer at 1 ml/minute using a Cytiva Akta FPLC. The affinity resin was washed with 20 column volumes of lysis buffer, and the protein eluted with 10 column volumes of lysis buffer containing 200 mM imidazole. The pooled SUMO-nsp13-containing fractions was dialyzed overnight into two changes of 20 mM imidazole-containing lysis buffer, and the SUMO tag cleaved with Ulp-1 for 4 hours at 4°C. Digestion was confirmed by SDS-PAGE analysis. The His6-Ulp-1 and His6-SUMO proteins were separated from the native nsp13 and nsp13D with a second round of $Ni^{2+}$-affinity chromatography as before. Helicase-containing flow-thru fractions were pooled, dialyzed overnight against two changes of low salt buffer (50 mM sodium phosphate, pH 6.8, 150 mM NaCl, 4 mM β-mercaptoethanol, 0.5 mM EDTA and 10% glycerol) and passed through a HighTrap SP (Cytiva) cation exchange column. Under these conditions, neither nsp13-helicase prteins adhered to the SP column. The proteins were concentrated with an Amicon Ultra-15 centrifugation filter units to a volume of ~1.5 mls and loaded on to a Sephacyl S200-HR HiPrep 26/60 column (Cytiva) equilibrated with nsp13-helicase Storage Buffer (25 mM HEPES, pH 7.5, 150 mM NaCl, 0.5 mM TCEP and 20% glycerol). The final nsp13-helicase proteins were quantified by UV spectrophotometry at 280 nm using the expected extinction coefficient of 68,785 $M^{-1}$ $cm^{-1}$ and confirmed using the BCA Protein Assay (Pierce). Protein samples were aliquoted, flash frozen, and stored at -80°C.

**Recombinant Protein Expression of poliovirus RNA-dependent RNA polymerase.**

Poliovirus RdRp was expressed and purified as previously reported (Seifert, van Nies et al. 2020). Briefly, expression was performed at 25°C by auto-induction, cells harvested, lysed by French Press, subjected to PEI precipitation followed by AMS04 precipitation, Ni-NTA chromatography, cleavage by Ulp1, phosphocellulose chromatography, gel filtration and the protein concentrated using Vivaspin concentrators.

**Construct fabrication ssRNA template.**

The fabrication of the ssRNA template (RNA hairpin) has been described in detail in Ref. (Papini, Seifert et al. 2019). In brief, the RNA hairpin is made of a 499 bp double-stranded RNA stem terminated by a 20 nt loop that is assembled from three ssRNA annealed together, and two handles, one of 856 bp at the 5′ end and one 822 bp at the 3′ end. The handles include either a 343 nt digoxygenin-labeled ssRNA or a 443 nt biotin-labeled ssRNA to attach to the anti-digoxigenin coated glass surface and the streptavidin-coated magnetic bead (M270), respectively.

**Construct fabrication dsRNA template.**

The RNA construct used here has been previously described in detail in Ref. (Seifert, van Nies et al. 2020). In brief, a 4 kb long single-stranded splint to which four ssRNAs are annealed: a biotin-labeled strand to attach to the streptavidin-coated magnetic bead (M270), a spacer, ~ 2.9 kb template, and a digoxygenin-labeled strand to attach to the glass surface. The template strand ends in 3′-end with a small hairpin with the sequence ACGCUUUCGCGT followed by 15 U residues to initiate poliovirus RdRp catalyzed RNA synthesis via primer extension.

**High-throughput and ultra-stable magnetic tweezers apparatus.**

The high-throughput magnetic tweezers used in this study have already been described in detail elsewhere (Quack and Dulin 2024). Briefly, two vertically aligned permanent magnets (5 mm cubes, SuperMagnete, Switzerland) separated by a 1 mm gap are positioned above a flow cell (see paragraph below) which is mounted on a custom-built inverted microscope. The vertical position and rotation of the magnets are controlled by two linear motors, M-126-PD1 and C-150 (Physik Instrumente PI, GmbH & Co. KG, Karlsruhe, Germany), respectively. The field of view is illuminated through the magnets gap by a collimated LED-light source and is imaged onto a large chip CMOS camera (Dalsa Falcon2 FA-80-12M1H, Stemmer Imaging, Germany) using a 50× oil immersion objective (CFI Plan Achro 50 XH, NA 0.9, Nikon, Germany) and an achromatic doublet tube lens of 200 mm focal length and 50 mm diameter (Qioptic, Germany). To control the temperature, we used a system described in details in Ref. .(Seifert, van Nies et al. 2020). A flexible resistive foil heater with an integrated 10 MΩ thermistor

(HT10K, Thorlabs) is wrapped around the microscope objective and further insulated by several layers Kapton tape (KAP22-075, Thorlabs). The heating foil is connected to a PID temperature controller (TC200 PID controller, Thorlabs) to adjust the temperature within ~0.1 °C.

##### Flow-cell assembly.

The fabrication procedure for flow cells has been described in detail in Ref. (Quack and Dulin 2024). To summarize, we sandwiched a double layer of Parafilm by two #1 coverslips, the top one having one hole at each end serving as inlet and outlet, the bottom one being coated with a 0.1% m/V nitrocellulose dissolved in amyl acetate solution. The flow cell was mounted into a custom-built holder and rinsed with ~1 ml of 1x phosphate buffered saline (PBS). 3 µm diameter polystyrene reference beads were attached to the bottom coverslip surface by incubating 100 µl of a 1:1000 dilution in PBS (LB30, Sigma Aldrich, stock conc.: 1.828*1011 particles per milliliter) for ~3 minutes. After rinsing the flow cell with 0.5 ml of PBS, 50 µl of anti-digoxigenin (50 µg/ml in PBS) was incubated for 30 minutes in the flow cell. The flow cell was then flushed with 1 ml of 10 mM Tris, 1 mM EDTA pH 8.0, 750 mM NaCl, 2 mM sodium azide to remove excess of anti-digoxigenin followed by rinsing with another 0.5 ml of TE buffer (10 mM Tris, 1 mM EDTA pH 8.0, 150 mM NaCl, 2 mM sodium azide). The surface was then passivated with a solution of bovine serum albumin (BSA, New England Biolabs, 10 mg/ml in PBS and 5% glycerol) for 30 minutes, and rinsed with 0.5 ml of TE buffer.

##### Single-molecule SARS-CoV-2 RTC activity experiments on dsRNA template.

20 µl of streptavidin coated Dynal Dynabeads M-270 streptavidin coated magnetic beads (ThermoFisher) was mixed with ~0.1 ng of RNA (total volume 40 µl) and incubated for ~5 minutes before rinsing with ~2 ml of TE buffer to remove any unbound RNA and the magnetic beads in excess. RNA tethers were sorted by looking for the characteristic extension of the correct length (~1 µm at 40 pN) due to the stretching of the dsRNA during a force ramp experiment (Papini, Seifert et al. 2019). The flow cell was subsequently rinsed with 0.5 ml reaction buffer (50 mM HEPES pH 7.9, 10 mM DTT, 2 µM EDTA, 5 mM $MgCl_2$). After starting the data acquisition at the indicated force, 100 µl of reaction buffer containing 0.2 µM nsp12, 1.8 µM nsp7, 1.8 µM nsp8 (1:9:9 stoichiometry), the indicated concentration of nsp13-helicase (WT or mutant if used) and 1 mM NTP were flushed in the flow cell to start the reaction. For the pre-assembled polymerase experiments, all or some of the nsp's were incubated for five minutes in the flow cell, while applying 25 pN force on the tether. The excess RTC proteins were subsequently flushed away with 0.3 ml of reaction buffer (flow cell volume ~ 40 µl), followed by the injection of 100 µl of reaction buffer with desired nsp(s) and 1 mM NTP. The experiments were conducted at a constant force for a duration of 30 to 60 minutes. The camera frame rate and the temperature were respectively set to 58 Hz and 25°C. A custom written Labview routine controlled the data acquisition and the (x-, y-, z-) positions analysis/tracking of both the magnetic and

reference beads in real-time (Cnossen, Dulin et al. 2014). Mechanical drift correction was performed by subtracting the reference bead position from the magnetic bead positions and further corrected using an autofocus (i.e. along the z-axis) protocol previously described in Ref. (Bera, Seifert et al. 2021).

#### Single-molecule poliovirus RdRp activity experiments on dsRNA template.

20 µl of streptavidin coated Dynal Dynabeads M-270 streptavidin coated magnetic beads (ThermoFisher) was mixed with ~0.1 ng of RNA (total volume 40 µl) and incubated for ~5 minutes before rinsing with ~2 ml of TE buffer to remove any unbound RNA and the magnetic beads in excess. RNA tethers were sorted for functional dsRNA by looking for its characteristic contour length, i.e. ~1 µm. The flow cell was subsequently rinsed with 0.5 ml of reaction buffer (50 mM HEPES pH 7.9, 10 mM DTT, 2 µM EDTA, 5 mM $MgCl_2$). After starting the data acquisition at a suitable force, 100 µl of reaction buffer containing 0.5 µM poliovirus RdRp, indicated concentration of nsp13-helicase and 1 mM NTP were flushed in the flow cell to start the reaction. The experiments were conducted at a constant force for a duration of 30 to 60 minutes.

#### Single-molecule SARS-CoV-2 RTC activity experiments on ssRNA template.

20 µl of streptavidin-coated Dynal Dynabeads M-270 streptavidin-coated magnetic beads (ThermoFisher Scientific) was mixed with ~0.1 ng of RNA hairpin (total volume 40 µl) and incubated for ~5 min before rinsing with ~2 ml of TE buffer to remove any unbound RNA and the magnetic beads in excess. RNA tethers were sorted for functional hairpins by looking for the characteristic jump in extension of the correct length (~0.6 µm at 30 pN) due to the sudden opening of the hairpin during a force ramp experiment (Papini, Seifert et al. 2019). The flow cell was subsequently rinsed with 0.5 ml reaction buffer (50 mM HEPES pH 7.9, 10 mM DTT, 2 µM EDTA, and 5 mM MgCl2). After starting the data acquisition at a force (~25 pN) that would keep the hairpin open, 100 µl of reaction buffer containing 0.6 µM of nsp12, 1.8 µM of nsp7 and nsp8 and 20 nM of nsp13-helicase, 500 µM NTP were flushed in the flow cell to start the reaction. The experiments were conducted at a constant force for a duration of 30 to 40 minutes. The camera frame rate was fixed at 58 Hz the temperature set to 25°C. A custom written LabVIEW routine controlled the data acquisition and the (x-, y-, z-) positions analysis/tracking of both the magnetic and reference beads in real time. Mechanical drift correction was performed by subtracting the reference bead position to the magnetic bead position and by applying an autofocus as described in Ref. (Bera, Seifert et al. 2021).

#### Data processing.

The activity traces of SARS-CoV-2 RTC or PV RdRp on dsRNA template were first corrected from the mechanical drift by subtracting the reference bead position to the tethers position, then converted from micron to nucleotides using the difference in extension under the same tension of dsRNA and ssRNA, as previously described (Dulin, Vilfan et al. 2015). The traces were subsequently low-pass filtered at 0.5 Hz and the dwell-times were extracted using a dwell-time window of 10 nt, as previously described (Seifert, van Nies et al. 2020). Similarly, the data acquired on ssRNA template were filtered at 2 Hz and subsequently converted in nucleotide (Bera, Seifert et al. 2021). Dwell-times were extracted as for the data on the ssRNA. The dwell-times of all the traces for a given experimental condition were assembled into a single distribution and further analyzed using a maximum likelihood estimation (MLE) fitting routine to extract the parameters from the dwell-time fit-function (**Supplementary Materials Section S1**).

#### Maximum likelihood estimation fitting routine.

The dwell-time distributions were fitted to the experimentally collected dwell-times $\{t_i\}$ by maximizing the log-likelihood function

$$LL = \sum_i \ln P_{N\mathrm{nt}}(t_i) \qquad \textbf{(1),}$$

Where $P_{N\mathrm{nt}}$ represents the probability of every dwell-time $t_i$ in the empirical distribution based on the dwell-time fit-function, We calculated the statistical error on the parameters by applying the MLE fitting procedure on 100 bootstraps of the original data set, and reported the standard deviation for each fitting parameter.

To be able to compare fits of models with different number of parameters, we calculated the Bayesian Information Criterion (BIC)

$$\mathrm{BIC} = k\ln(N) - 2\ln(LL). \qquad \textbf{(2).}$$

This criterion compares the log-likelihood $LL$ to the number of parameters in the model $k$ and the number of datapoints fitted $N$ to account for the fact that using more parameters can lead to overfitting. The model fit with minimized BIC is considered the best fit with optimal number of parameters for the set of model fits tested (Konishi and Kitagawa 2008).

#### Dwell-time fit-function for nucleotide addition by SARS-CoV-2 core RTC.

The dwell-time distributions of the SARS-CoV-2 RTC in absence of active nsp13-helicase were fitted with a dwell-time fit-function consisting of one gamma distribution with characteristic timescale $T_{\mathrm{FNA}}$ fitting the peak at short timescale, two exponential distributions with characteristic timescales $T_{\mathrm{SNA}}$ and $T_{\mathrm{VSNA}}$ fitting two bumps in the dwell-time distributions and a power law distribution of $\sim t^{-3/2}$ fitting the fat-tail in the dwell-time distributions for longer timescales (Depken, Galburt et al. 2009)

$$P_{N\mathrm{nt}}(t) \approx \frac{f_{\mathrm{FNA}}N}{T_{\mathrm{FNA}}(N-1)!}\left(\frac{tN}{T_{\mathrm{FNA}}}\right)^{N-1} e^{-\frac{tN}{T_{\mathrm{FNA}}}} + Q(t)\left(\frac{f_{\mathrm{SNA}}}{T_{\mathrm{SNA}}}e^{-\frac{t-T_{\mathrm{FNA}}}{T_{\mathrm{SNA}}}} + \frac{f_{\mathrm{VSNA}}}{T_{\mathrm{VSNA}}}e^{-\frac{t-T_{\mathrm{FNA}}}{T_{\mathrm{VSNA}}}} + \frac{f_{\mathrm{LLP}}\sqrt{1+T_{\mathrm{FNA}}}}{2\left(1+\frac{t}{1s}\right)^{\frac{3}{2}}}\right) \quad \textbf{(3),}$$

The approximation of the SNA, VSNA and LLP dominated terms break down for the short timescales since $N$ sequential steps always need to be taken to get through the dwell-time window. This is accounted for by regularization function $Q(t) = \frac{(t/T_{\mathrm{FNA}})^{N-1}}{1+(t/T_{\mathrm{FNA}})^{N-1}}$ and normalization of the terms starting from the peak position of the gamma distribution $T_{\mathrm{FNA}}$. $\sum_j f_j = 1$ for $j \in \{\mathrm{FNA, SNA, VSNA, LLP}\}$ ensured the distribution $P_{N\mathrm{nt}}(t)$ is normalized.

Considering that we observed a clearly separated peak and two bumps in the dwell-time distributions, we distinguished three characteristic timescales in the dwell-time distribution dominated by fast, slow and very slow nucleotide addition and thus assumed clear separation of timescales for single nucleotide additions $\tau_{\mathrm{FNA}} \ll \tau_{\mathrm{SNA}} \ll \tau_{\mathrm{VSNA}}$. With this assumption, we derived relations for the characteristic timescales and probabilities on the dwell-time level in terms of single nucleotide timescales and probabilities as done in (Bera, Seifert et al. 2021) (**Supplementary Information Section S1**).

For the power law distribution representing a long-lived pause, we have introduced a regularization at 1 s, but the precise timescale does not matter here, as long as it is set within the region dominated by either of the FNA, SNA or VSNA pathways. Due to this approximation, the long-lived pause probability $f_{\mathrm{LLP}}$ should be interpreted as the relative probability to enter the long-lived pause. For a more elaborate discussion, see **Supplementary Information Section S1**.

**Dwell-time fit-function for RNA synthesis on dsRNA by the SARS-CoV-2 RTC in complex with active nsp13-helicase.**

The dwell-time distributions for elongation dynamics by the SARS-CoV-2 RTC with active nsp13-helicase were fitted with a dwell-time fit-function consisting of two gamma distributions with characteristic timescales $T_{\mathrm{VFNA}}$ and $T_{\mathrm{FNA}}$ fitting the two peaks at short timescale, two exponential distributions with timescales $T_{\mathrm{SNA}}$ and $T_{\mathrm{VSNA}}$ fitting two bumps in the dwell-time distributions and a power law distribution $\sim t^{-3/2}$ fitting the fat-tail in the dwell-time distributions for longer timescales (**Fig. 3B**).

The dwell-time fit-function for the dwell-time distributions of RTC elongation dynamics on a dsRNA template in presence of active nsp13-helicase reads

$$P_{N\mathrm{nt}}(t) \approx \frac{f_{\mathrm{VFNA}}N}{T_{\mathrm{VFNA}}(N-1)!}\left({tN}/{T_{\mathrm{VFNA}}}\right)^{N-1} e^{-tN/T_{\mathrm{VFNA}}} + \frac{f_{\mathrm{FNA}}N}{T_{\mathrm{FNA}}(N-1)!}\left({tN}/{T_{\mathrm{FNA}}}\right)^{N-1} e^{-tN/T_{\mathrm{FNA}}} + Q(t)\left(\frac{f_{\mathrm{SNA}}}{T_{\mathrm{SNA}}}e^{-\frac{t-T_{\mathrm{FNA}}}{T_{\mathrm{SNA}}}} + \frac{f_{\mathrm{VSNA}}}{T_{\mathrm{VSNA}}}e^{-\frac{t-T_{\mathrm{FNA}}}{T_{\mathrm{VSNA}}}} + \frac{f_{\mathrm{LLP}}\sqrt{1+T_{\mathrm{FNA}}}}{2(1+t/1s)^{\frac{3}{2}}}\right) \quad \textbf{(4),}$$

Where the regularization function $Q(t) = \frac{(t/T_{\mathrm{FNA}})^{N-1}}{1+(t/T_{\mathrm{FNA}})^{N-1}}$ was used and the SNA, VSNA and LLP dominated terms were normalized starting from the peak position of the gamma distribution $T_{\mathrm{FNA}}$, like in the dwell-time fit-function for the core RTC. $\sum_j f_j = 1$ for $j \in \{\mathrm{VFNA}, \mathrm{FNA}, \mathrm{SNA}, \mathrm{VSNA}, \mathrm{LLP}\}$ ensured the distribution $P_{N\mathrm{nt}}(t)$ is normalized. Considering two peaks and two bumps could be clearly distinguished in the empirical dwell-time distributions, we assumed clear separation of single nucleotide timescales $\tau_{\mathrm{VFNA}} \ll \tau_{\mathrm{FNA}} \ll \tau_{\mathrm{SNA}} \ll \tau_{\mathrm{VSNA}}$. The power law distribution has a regularization at 1 s, since the underlying kinetics are not expected to change in presence of active nsp13-helicase. Due to this approximation, the long-lived pause probability $f_{\mathrm{LLP}}$ should be interpreted as the relative probability to enter the long-lived pause, as explained for the dwell-time fit-function for the core RTC. For a more elaborate discussion, see **Supplementary Information Section S1**.

No significant trends were observed in the VFNA and VSNA characteristic timescales for the conditions measured, so they were fixed to $T_{\mathrm{VFNA}} = 0.2$ s and $T_{\mathrm{VSNA}} = 4.8$ s respectively (**Supplementary Information Section S1**).

**Assembly model for the RTC-nsp13-helicase complexes on a dsRNA template.**

We constructed an RTC-nsp13 assembly model including four RTC complexes, i.e. the RTC without nsp13-helicase (core RTC), the RTC with nsp13.1 bound (RTC-nsp13.1), the RTC with the two helicases bound and nsp13.2 not engaged (RTC-nsp13.1,2), the RTC with the two nsp13-helicases bound and nsp13.2 engaged with the non-template RNA (RTC-nsp13.1,2*) (**Fig. 3B**).

We observed very fast nucleotide addition bursts spanning tens to hundreds of nucleotides already for low nsp13-helicase concentration on a dsRNA template (**Fig. 1C**), meaning that a single RTC-nsp13.1,2* complex is stable for much longer than the timescale for synthesizing 10 nt RNA (dwell-time window). To discriminate between assembly models, we assumed the binding of two nsp13-helicases to the RTC is in dynamic equilibrium, but they do not exchange within one dwell-time window. Under this assumption, we can write the fractional occupancies $p_{\mathrm{cpx}}$ of the complexes cpx $\in \{\mathrm{c}, 1{,}2, 2^*\} = \{\text{core RTC; RTC} - \text{nsp13.1; RTC} - \text{nsp13.1,2; RTC} - \text{nsp13.1,2}^*\}$ in terms of the free-energy barriers between the complexes $\Delta G_{\mathrm{cpx}\to\mathrm{cpx}'}$

$$
\begin{aligned}
p_{2^*} &= \frac{[\mathrm{nsp13}]^2 e^{-\left(\Delta G_{\mathrm{c}\to 1}+\Delta G_{1\to 2}+\Delta G_{2\to 2^*}\right)}}{Z},\\
p_2 &= \frac{[\mathrm{nsp13}]^2 e^{-\left(\Delta G_{\mathrm{c}\to 1}+\Delta G_{1\to 2}\right)}}{Z},\\
p_1 &= \frac{[\mathrm{nsp13}] e^{-\Delta G_{\mathrm{c}\to 1}}}{Z},\\
p_c &= \frac{1}{Z},
\end{aligned}
\qquad \textbf{(5),}
$$

With $Z = [\mathrm{nsp13}]^2 e^{-\left(\Delta G_{c\to 1}+\Delta G_{1\to 2}+\Delta G_{2\to 2^*}\right)} + [\mathrm{nsp13}]^2 e^{-\left(\Delta G_{c\to 1}+\Delta G_{1\to 2}\right)} + [\mathrm{nsp13}] e^{-\Delta G_{c\to 1}} + 1$. The free-energy barriers between the complexes $\Delta G_{\mathrm{cpx}\to\mathrm{cpx}'}$ are expressed in units of $\mathrm{k_B T}$ with 1 nM as reference concentration. Here we used that the fraction of product of every single equilibrium reaction is given by a Boltzmann factor dependent on the nsp13-helicase concentration and the free-energy barrier (Phillips, Kondev et al. 2009).

Furthermore, the association constants for nsp13.1 and nsp13.2 to the RTC were obtained from the free-energy barriers in the assembly model as

$$K_{\mathrm{a,nsp13.1}} = e^{-\Delta G_{\mathrm{c}\to 1}}; \quad K_{\mathrm{a,nsp13.2}} = e^{-\Delta G_{1\to 2}} \qquad \textbf{(6),}$$

Since we used 1 nM as reference concentration, the association constants are giving in the unit $\mathrm{nM^{-1}}$.

#### Integrating the assembly of the RTC-nsp13-helicase complexes with the elongation dynamics on a dsRNA template.

We then built the RTC Assembly-Elongation dynamics model connecting the RTC assembly model with the elongation dynamics (dwell-time fit-function) for a mix of core RTC's and RTC-nsp13 complexes with a constant tension on the non-template RNA strand.

We assumed that the RTC resides exclusively in one of the RTC-nsp13 states within one dwell-time window. Under this assumption, the first passage time distribution for elongation dynamics is obtained by summing the first passage time distributions $P_{10\mathrm{nt}}(t|\mathrm{cpx})$ for the different RTC complexes cpx $\in \{\mathrm{c}, 1, 2, 2^*\}$ with the fractional occupancies for each RTC complex $p_{\mathrm{cpx}}$ as pre-factor

$$P_{N\text{nt,RTC-nsp13}}(t) = \sum_{\text{cpx}} p_{\text{cpx}} P_{10\text{nt}}(t|\text{cpx}) \quad \textbf{(7),}$$

are the first passage time distributions conditioned that the RTC resides in complex cpx for the 10 nt of interest. We approximated the first passage time distribution for each RTC complex by the dwell-time fit-function (**Eq. 3**). Considering that $P_{10\text{nt}}(t|\text{cpx})$ is also a summation of terms with separated characteristic timescales $T_{\text{NA}}$ for each pathway NA $\in$ {VFNA, FNA, SNA, VSNA}, and that each of these timescales stay within one order of magnitude from the core RTC to saturating nsp13-helicase condition (**Fig. 3C**), we obtained the average characteristic timescale of each NA pathway $T_{\text{NA}}$ and the combined probability $f_{\text{NA}}$ in the RTC-nsp13 mixture, referred to as the effective parameters (**Supplementary Information Section S2.2**)

$$\begin{aligned}
f_{\text{VFNA}} &= p_{2^*}\,, \\
T_{\text{VFNA}} &= N\tau_{\text{VFNA}}\,, \\
f_{\text{FNA}} &= \sum_{\text{cpx}} p_{\text{cpx}} \left(p_{\text{FNA}}^{\text{cpx}}\right)^N, \\
T_{\text{FNA}} &= \frac{\sum_{\text{cpx}} p_{\text{cpx}} \left(p_{\text{FNA}}^{\text{cpx}}\right)^N N\tau_{\text{FNA}}^{\text{cpx}}}{\sum_{\text{cpx}} p_{\text{cpx}} \left(p_{\text{FNA}}^{\text{cpx}}\right)^N}, \\
f_{\text{SNA}} &= \sum_{\text{cpx}} p_{\text{cpx}} \left[\left(p_{\text{FNA}}^{\text{cpx}} + p_{\text{SNA}}^{\text{cpx}}\right)^N - \left(p_{\text{FNA}}^{\text{cpx}}\right)^N\right], \\
T_{\text{SNA}} &= \frac{\sum_{\text{cpx}} p_{\text{cpx}} p_{\text{SNA}}^{\text{cpx}} \left(p_{\text{FNA}}^{\text{cpx}} + p_{\text{SNA}}^{\text{cpx}}\right)^{N-1} N\tau_{\text{SNA}}^{\text{cpx}}}{\sum_{\text{cpx}} p_{\text{cpx}} \left[\left(p_{\text{FNA}}^{\text{cpx}} + p_{\text{SNA}}^{\text{cpx}}\right)^N - \left(p_{\text{FNA}}^{\text{cpx}}\right)^N\right]}, \\
f_{\text{VSNA}} &= \sum_{\text{cpx}} p_{\text{cpx}} \left[\left(p_{\text{FNA}}^{\text{cpx}} + p_{\text{SNA}}^{\text{cpx}} + p_{\text{VSNA}}^{\text{cpx}}\right)^N - \left(p_{\text{FNA}}^{\text{cpx}} + p_{\text{SNA}}^{\text{cpx}}\right)^N\right], \\
T_{\text{VSNA}} &= \frac{\sum_{\text{cpx}} p_{\text{cpx}} p_{\text{VSNA}}^{\text{cpx}} \left(p_{\text{FNA}}^{\text{cpx}} + p_{\text{SNA}}^{\text{cpx}} + p_{\text{VSNA}}^{\text{cpx}}\right)^{N-1} N\tau_{\text{VSNA}}^{\text{cpx}}}{\sum_{\text{cpx}} p_{\text{cpx}} \left[\left(p_{\text{FNA}}^{\text{cpx}} + p_{\text{SNA}}^{\text{cpx}} + p_{\text{VSNA}}^{\text{cpx}}\right)^N - \left(p_{\text{FNA}}^{\text{cpx}} + p_{\text{SNA}}^{\text{cpx}}\right)^N\right]}
\end{aligned} \quad \textbf{(8),}$$

With $\text{cpx} \in \{\text{c}, 1, 2\}$. For a fit to the dwell-time distributions, the expressions for the average characteristic timescales and combined probabilities **Eq. 8** were substituted into the dwell-time fit function **Eq. 4**. Furthermore, we added a relative probability to enter the long-lived pause for each RTC complex $p_{\text{LLP}}^{\text{cpx}}$ such that $p_{\text{FNA}}^{\text{cpx}} + p_{\text{SNA}}^{\text{cpx}} + p_{\text{VSNA}}^{\text{cpx}} + p_{\text{LLP}}^{\text{cpx}} = 1$ and thus $\sum_{\text{j}} f_{\text{j}} = 1$ for $\text{j} \in \{\text{FNA, SNA, VSNA, LLP}\}$. The VFNA and VSNA characteristic timescales were fixed to $T_{\text{VFNA}} = 0.2$ s and $T_{\text{VSNA}} = 4.8$ s respectively, so the corresponding single nucleotide timescales $\tau_{\text{VFNA}}, \tau_{\text{VSNA}}^{\text{c}}, \tau_{\text{VSNA}}^{2}$ were not fitted, but could be directly retrieved from **Eq. 8**.

As explained in the **Results** section, we kept all single nucleotide parameters in the RTC-nsp13.1 complex the same as for the core RTC, while the parameters were free variables for the RTC-nsp13.1,2 complex. The free parameters in the RTC assembly - Elongation dynamics model were the single nucleotide probabilities $p_{\text{FNA}}^{\text{c}}, p_{\text{SNA}}^{\text{c}}, p_{\text{VSNA}}^{\text{c}}, p_{\text{FNA}}^{2}, p_{\text{SNA}}^{2}, p_{\text{VSNA}}^{2}$ and the single nucleotide timescales $\tau_{\mathrm{FNA}}^{\mathrm{c}}$, $\tau_{\mathrm{SNA}}^{\mathrm{c}}$, $\tau_{\mathrm{FNA}}^{2}$, $\tau_{\mathrm{SNA}}^{2}$ for the core RTC (c) and RTC-nsp13.1,2 (2) complex and the free-energy barriers in the RTC assembly model $\Delta G_{\mathrm{c}\to 1}$, $\Delta G_{1\to 2}$, $\Delta G_{2\to 2^*}$.

To perform a global fit on the empirical dwell-time distributions versus nsp13-helicase concentration, we obtained the average characteristic timescales and the combined probability for each pathway at every nsp13-helicase concentration measured from the RTC Assembly - Elongation dynamics model (**Eq. 8** with **Eq. 5** substituted) and substituted these parameters into the dwell-time fit-function **Eq. 4** with VFNA and VSNA characteristic timescales fixed to $T_{\mathrm{VFNA}} = 0.2$ s and $T_{\mathrm{VSNA}} = 4.8$ s respectively. In this way, we obtained a probability density function for each condition measured, which was successfully fitted to the dwell-time distributions (**Fig. 3A**) using the Maximum likelihood fitting routine (see Materials and Methods).

#### Mechanochemical model underlying the RNA tension dependency of the RTC elongation dynamics.

The force dependence of a kinetic rate can be captured by the Arrhenius law

$$\boldsymbol{k_{\mathrm{X}}(F) = k_{\mathrm{X},0} e^{\delta_X F / k_{\mathrm{B}} T}} \qquad \textbf{(9),}$$

Where $\delta_X$ is the distance to the transition state and $k_{\mathrm{X},0}$ is the rate at zero force. In the case of a dsRNA template, the force (tension) is applied on the non-template RNA strand, which acts as an assisting force on elongation by the RTC.

Our mechanochemical model (**Fig. 4G**) is derived based on the trends observed in the characteristic timescales and probabilities versus RNA tension with or without saturating nsp13-helicase concentration (**Results, Fig. 4C-F, Supplementary Information**). We only have a force (RNA tension) dependence on the backward translocation rates in the FNA and SNA pathway, because Bera et al. (2021) found that the distance to the transition state for forward transition by the RTC is negligible. As translocation acts over the distance from the conversion of one base pair dsRNA to ssRNA $a \approx 0.2$ nm (Dulin et al. 2015), the opposing RNA tension ($F$) dependency on the backward translocation rate could be described as the Arrhenius law with distance $-a$

$$k_{\mathrm{pre}}^{\mathrm{NA,cpx}}(F) = k_{\mathrm{pre}}^{\mathrm{NA,cpx}}(0) e^{-aF/k_{\mathrm{B}}T}, \qquad \mathrm{NA} \in \{\mathrm{FNA}, \mathrm{SNA}\}, \qquad \mathrm{cpx} \in \{\mathrm{c}, 1, 2\} \qquad \textbf{(10),}$$

To obtain the expressions for the timescale and probability for a single nucleotide addition in the pathways that showed an RNA tension dependency in the characteristic timescale (FNA and SNA) for each RTC complex $\mathrm{cpx} \in \{\mathrm{c}, 1,2\}$, we constructed the first passage time distribution for completing the irreversible step in the NA pathway. Considering that translocation is not equilibrated, the first passage time (FPT) distributions was obtained by summing over all possible paths with forward and backward translocation (with rates $k_{\mathrm{post}}^{\mathrm{NA,cpx}}$ and $k_{\mathrm{pre}}^{\mathrm{NA,cpx}}(F)$ respectively) followed by NTP binding and incorporation with rate $k_{\mathrm{irr}}^{\mathrm{NA,cpx}}$ (**Fig. 4G, Supplementary Information Section S3**). From this FPT distribution, we obtained the probability and timescale of completing the irreversible step in pathway $\mathrm{NA} \in \{\mathrm{FNA}, \mathrm{SNA}\}$

$$p_{\mathrm{irr}}^{\mathrm{NA,cpx}} = \frac{k_{\mathrm{irr}}^{\mathrm{NA}} k_{\mathrm{post}}^{\mathrm{NA}}}{\left(k_{\mathrm{post}}^{\mathrm{NA}}+k_{\mathrm{out}}^{\mathrm{NA}}\right)\left(k_{\mathrm{pre}}^{\mathrm{NA,cpx}}(F)+k_{\mathrm{irr}}^{\mathrm{NA}}\right)-k_{\mathrm{post}}^{\mathrm{NA}} k_{\mathrm{pre}}^{\mathrm{NA,cpx}}(F)};$$
$$\tau_{\mathrm{irr}}^{\mathrm{NA,cpx}} = \frac{k_{\mathrm{post}}^{\mathrm{NA}}+k_{\mathrm{out}}^{\mathrm{NA}}+k_{\mathrm{pre}}^{\mathrm{NA,cpx}}(F)+k_{\mathrm{irr}}^{\mathrm{NA}}}{\left(k_{\mathrm{post}}^{\mathrm{NA}}+k_{\mathrm{out}}^{\mathrm{NA}}\right)\left(k_{\mathrm{pre}}^{\mathrm{NA,cpx}}(F)+k_{\mathrm{irr}}^{\mathrm{NA}}\right)-k_{\mathrm{post}}^{\mathrm{NA}} k_{\mathrm{pre}}^{\mathrm{NA,cpx}}(F)}; \qquad \textbf{(11),}$$

With $k_{\mathrm{out}}^{\mathrm{FNA}} = k_{\mathrm{in}}^{\mathrm{SNA}} + k_{\mathrm{in}}^{\mathrm{LLP}};\ k_{\mathrm{out}}^{\mathrm{SNA}} = k_{\mathrm{in}}^{\mathrm{VSNA}} + k_{\mathrm{in}}^{\mathrm{LLP}}$.

Considering that the slowest pathway entered is dominating the timescale of the single nucleotide addition, we substituted $\tau_{\mathrm{NA}}^{\mathrm{cpx}} = \tau_{\mathrm{irr}}^{\mathrm{NA,cpx}}$. To obtain the single nucleotide probability for each pathway $\mathrm{NA} \in \{\mathrm{FNA}, \mathrm{SNA}, \mathrm{VSNA}\}$ and the RTC complexes $\mathrm{cpx} \in \{1,2,2^*\}$, we had to consider how the pathways are connected. Each NA cycle starts from the FNA pre-translocated state, from which the SNA pre-translocated state can be entered and the VSNA pre-translocated state from there (**Fig. 4G**). Furthermore, the long-lived pause can be entered from each NA pre-translocated state. With these connections, the single nucleotide probabilities for exiting through the pathways were obtained as

$$p_{\mathrm{FNA}}^{\mathrm{cpx}}(F) = p_{\mathrm{irr}}^{\mathrm{FNA,cpx}}(F);$$
$$p_{\mathrm{SNA}}^{\mathrm{cpx}}(F) = \left(1 - p_{\mathrm{irr}}^{\mathrm{FNA,cpx}}(F)\right)\frac{k_{\mathrm{in}}^{\mathrm{SNA}}}{k_{\mathrm{out}}^{\mathrm{FNA}}} p_{\mathrm{irr}}^{\mathrm{SNA,cpx}}(F);$$
$$p_{\mathrm{VSNA}}^{\mathrm{cpx}}(F) = \left(1 - p_{\mathrm{irr}}^{\mathrm{FNA,cpx}}(F)\right)\frac{k_{\mathrm{in}}^{\mathrm{SNA}}}{k_{\mathrm{out}}^{\mathrm{FNA}}}\left(1 - p_{\mathrm{irr}}^{\mathrm{SNA,cpx}}(F)\right)\frac{k_{\mathrm{in}}^{\mathrm{VSNA}}}{k_{\mathrm{out}}^{\mathrm{SNA}}} p_{\mathrm{irr}}^{\mathrm{VSNA,cpx}}; \qquad \textbf{(12),}$$
$$p_{\mathrm{LLP}}^{\mathrm{cpx}}(F) = 1 - p_{\mathrm{FNA}}^{\mathrm{cpx}}(F) - p_{\mathrm{SNA}}^{\mathrm{cpx}}(F) - p_{\mathrm{VSNA}}^{\mathrm{cpx}}(F)$$

The allosteric effect of nsp13.2 binding to the RTC, resulting in decreased FNA and SNA single nucleotide timescale accompanied by a decreased SNA and VSNA probability for the RTC-nsp13.1,2 complex (**Table S2**), can be modelled as an increase in the free-energy barrier $\Delta G_{\mathrm{NA}}$ for backward translocation rate in each NA pathway

$$k_{\mathrm{pre}}^{\mathrm{NA,2}}(F) = k_{\mathrm{pre}}^{\mathrm{NA,c}}(F) e^{\Delta G_{\mathrm{NA}}}, \qquad \mathrm{NA} \in \{\mathrm{FNA}, \mathrm{SNA}\} \qquad \textbf{(13).}$$

Since the VSNA characteristic timescale was fixed in the fits, the underlying mechanochemistry cannot be extracted from it, so we directly fit $p_{\mathrm{irr}}^{\mathrm{VSNA,cpx}}$ for $\mathrm{cpx} \in \{1, 2\}$.

As a result, the free parameters in the mechanochemical model were $k_{\mathrm{in}}^{\mathrm{SNA}}, k_{\mathrm{in}}^{\mathrm{VSNA}}, k_{\mathrm{in}}^{\mathrm{LLP}}, k_{\mathrm{post}}^{\mathrm{FNA}}, k_{\mathrm{post}}^{\mathrm{SNA}}, k_{\mathrm{pre}}^{\mathrm{FNA.c}}(0), k_{\mathrm{pre}}^{\mathrm{FNA.2}}(0), k_{\mathrm{pre}}^{\mathrm{SNA.c}}(0), k_{\mathrm{pre}}^{\mathrm{SNA.2}}(0), k_{\mathrm{irr}}^{\mathrm{FNA}}, k_{\mathrm{irr}}^{\mathrm{SNA}}, p_{\mathrm{irr}}^{\mathrm{VSNA.c}}, p_{\mathrm{irr}}^{\mathrm{VSNA.2}}$ giving the single nucleotide probabilities and timescales. The substitution into the RTC Assembly - Elongation dynamics model added the free-energy barriers between the RTC states $\Delta G_{c\to 1}$, $\Delta G_{1\to 2}$, $\Delta G_{2\to 2^*}$.

A global fit of the mechanochemical model was successfully performed on the dwell-time distributions of the nsp13-helicase concentration dependency dataset at 20 pN RNA tension and RNA tension dependence datasets without nsp13-helicase and at saturating nsp13-helicase concentration (20 nM) (**Fig. 3ACD & 4C-F**, **Fig. S14**). To fit the dwell-time distributions for each condition measured we had to obtain a fit-function for each condition measured. For this, we substituted the expressions for the single nucleotide probabilities and timescales versus RNA tension (**Eq. 10-13**) in the RTC Assembly - Elongation dynamics model (**Eq. 8** with **Eq. 5** substituted) to obtain the effective probabilities and timescales of each pathway on the dwell-time level and these parameters were substituted into the dwell-time fit-function **Eq. 4**. A more thorough explanation of the argumentation for and derivation of the mechanochemical model can be found in **Supplementary information Section S3**.

#### Performing fits, bootstrapping and fit selection.

For each model fit on the empirical dwell-time distributions, we performed multiple fits and selected the best fit. Then we performed 100 fits for each model on resampled dwell-time distributions, referred to as bootstrapping, from which we obtained the mean value and standard deviation on each parameter in the fitted model. With the goal to minimize the number of free parameters in the model without reducing the goodness-of-fit, we set certain parameters constant during the fitting and compared them by visual inspection and using the Bayesian Information Criterion (BIC) as a guide. For an elaborate discussion on this, refer to **Supplementary Information Section S1 and S2**.

### Supplementary Text

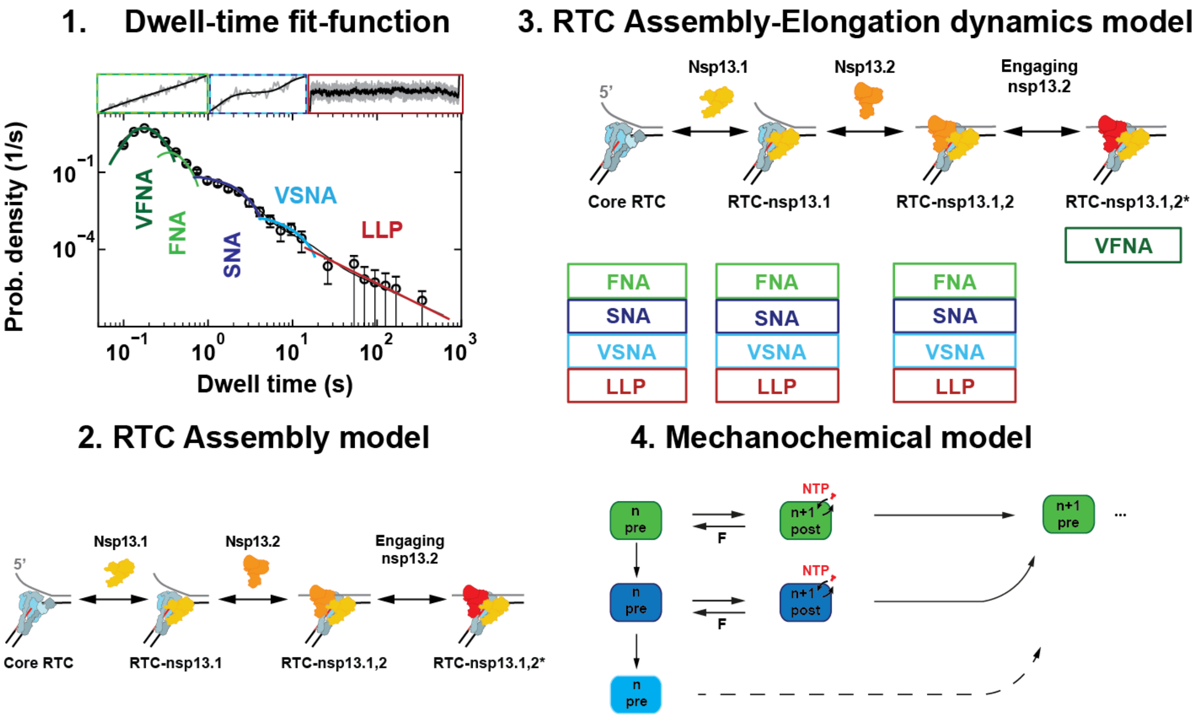

**Fig. S1. Schematic of the build up of the complete model describing elongation dynamics by the core RTC and RTC-nsp13 complexes on dsRNA. (1.)** To extract information from the dwell-time distributions for building our microscopic models, we constructed a dwell-time fit-function consisting of two peaks, representing very fast and fast nucleotide addition (VFNA and FNA), two shoulders, representing slow and very slow nucleotide addition (SNA and VSNA) and a tail of long dwell-times representing long-lived pause recovery (LLP). **(2.)** For RTC-nsp13 complex formation we derived an RTC Assembly model from comparison of elongation dynamics by the SARS-CoV-2 RTC with or without nsp13-helicase on single- or double-stranded RNA and structural studies. **(3.)** By connecting our observations on the elongation dynamics from the empirical dwell-time distributions versus nsp13-helicase concentration to the RTC Assembly model we derived the RTC Assembly-Elongation dynamics model. **(4.)** From the trends observed in the RNA tension dependency of the elongation dynamics for the core RTC and at saturating nsp13-helicase concentration we determine the underlying mechanochemical model.

### Section S1: Dwell-time fit-function

To extract kinetic information from our data, we constructed a fit-function to fit to the empirical dwell-time distributions.

#### Section S1.1: The structure of the empirical dwell-time distributions

The empirical dwell-time distributions for a 10 nt window showed the same signature trends, i.e. peaks and shoulders could be distinguished at different dwell-time durations (**Fig. 1F**), resulting from different processes. In presence of nsp13-helicase there were two peaks at short dwell-times corresponding to two fast multi-step processes, two shoulders at intermediate dwell-times corresponding to two intermediate time processes, and a slow process dominating the longest dwell-times resulting in a power-law tail in the distribution (**Fig. 1F**). To extract the characteristic times and probabilities from our data, we constructed a fit-function that captures the dominant behaviour at all dwell-time durations measured.

In the main manuscript we argued that the four short-to-intermediate characteristic times result from four different nucleotide addition pathways. We named these pathways very fast (VFNA), fast (FNA), slow (SNA) and very slow nucleotide addition (VSNA) depending on their characteristic timescales. In the main text we further argued that the power-law tail originates from long-lived pauses (LLP). Based on the data we argued that the RTC was typically on the RNA for many dwell-time windows before disengaging (**Fig. 1C**), and that a typical dwell-time window was either traversed by the VFNA cycle alone, or by a mix of FNA, SNA and VSNA cycles (**Fig. 4G**). Based on these assumptions, we constructed a dwell-time fit-function including all nucleotide addition (NA) pathways and long-lived pauses.

#### Section S1.2: Dwell-time fit-function for the core RTC

Before building the complete first-passage-time distributions for the nucleotide addition cycle of the SARS-CoV-2 RTC in complex with nsp13-helicase, we considered the distribution for the core RTC alone. In that case, a typical dwell-time window was traversed by a mix of FNA, SNA and VSNA cycles, which we approximated as having well-separated characteristic timescales ($\tau_{\text{FNA}} \ll \tau_{\text{SNA}} \ll \tau_{\text{VSNA}}$) (**Fig. 1E**). This approximation was justified by the timescales visible in **Fig. 3A**.

*Working in Laplace space*

Dwell-times correspond to the first-passage times from entry to exit of the dwell-time window. For dwell-time windows including many NA steps it becomes convenient to work with first-passage time distributions in Laplace space. In Laplace space the convolutions that arise when calculating the probability density of the total time to perform sequential steps become multiplications (Palani 2022). An exponential process with characteristic time $\tau$ and probability $p$ and has the probability-density function (pdf) $P(t) = pe^{-t/\tau}/\tau$ and its Laplace transform is given by

$$\psi(s) = \int_0^\infty \mathrm{d}t\, P(t)e^{-st} = \frac{p}{1 + \tau s}$$

With two sequential steps with respective pdfs $P_1(t_1)$ and $P_2(t_2)$, the Laplace transform of the pdf $P_{1+2}(t)$ of the total time $t = t_1 + t_2$ is simply given by

$$\psi_{1+2}(s) = \frac{p_1}{1+\tau_1 s}\frac{p_2}{1+\tau_2 s}.$$

The above can be done repeatedly for any number $N$ of processes, resulting in

$$\psi_{1+2+\cdots+N}(s) = \frac{p_1}{1+\tau_1 s}\frac{p_2}{1+\tau_2 s}\cdots\frac{p_N}{1+\tau_N s}.$$

The Laplace transform can in principle be inverted to give the exact form of $P_{1+2+\cdots+N}(t)$, but this can be computationally expensive and we instead applied approximations in Laplace space before reverting back to real time analytically (Palani 2022).

*The exact dwell-time distribution for FNA, SNA, and VSNA processes combined*

We started by constructing the first passage time distribution for short and intermediate (s&i) timescales by considering elongating across a $N$ nucleotides (nt) window that randomly alternates through FNA, SNA, VSNA in each step (Bera, Seifert et al. 2021). The first-passage time distribution for the core RTC in Laplace space is

$$\begin{aligned}\psi_{N\mathrm{nt}}^{\mathrm{s\&i}}(s) &= \left(\frac{p_{\mathrm{FNA}}}{1+\tau_{\mathrm{FNA}}s} + \frac{p_{\mathrm{SNA}}}{1+\tau_{\mathrm{SNA}}s} + \frac{p_{\mathrm{VSNA}}}{1+\tau_{\mathrm{VSNA}}s}\right)^N \\ &= \underbrace{p_{\mathrm{FNA}}^N\left(\frac{1}{1+\tau_{\mathrm{FNA}}s}\right)^N}_{\psi_{N\mathrm{nt}}^{\mathrm{FNA}}\text{: only FNA steps}} + \underbrace{\sum_{n_0=0}^{N-1}\binom{N}{n_0, N-n_0, 0} p_{\mathrm{FNA}}^{n_0} p_{\mathrm{SNA}}^{N-n_0}\left(\frac{1}{1+\tau_{\mathrm{FNA}}s}\right)^{n_0}\left(\frac{1}{1+\tau_{\mathrm{SNA}}s}\right)^{N-n_0}}_{\psi_{N\mathrm{nt}}^{\mathrm{SNA}}\text{: at least one SNA but no VSNA steps}} \\ &+ \underbrace{\sum_{n_0=0}^{N-1}\sum_{n_1=0}^{N-n_0-1}\binom{N}{n_0, n_1, N-n_0-n_1} p_{\mathrm{FNA}}^{n_0} p_{\mathrm{SNA}}^{n_1} p_{\mathrm{VSNA}}^{N-n_0-n_1}\left(\frac{1}{1+\tau_{\mathrm{FNA}}s}\right)^{n_0}\left(\frac{1}{1+\tau_{\mathrm{SNA}}s}\right)^{n_1}\left(\frac{1}{1+\tau_{\mathrm{VSNA}}s}\right)^{N-n_0-n_1}}_{\psi_{N\mathrm{nt}}^{\mathrm{VSNA}}\text{: at least one VSNA step}}\end{aligned} \quad \text{(S1)}$$

In order to work out the implication of the separation of timescales, we have separated out the terms that contain only FNA ($\psi_{N\mathrm{nt}}^{\mathrm{FNA}}$), at least one SNA but no VSNA ($\psi_{N\mathrm{nt}}^{\mathrm{SNA}}$), and those that contain at least one VSNA ($\psi_{N\mathrm{nt}}^{\mathrm{VSNA}}$).

The first term in **Eq. S1** ($\psi_{N\mathrm{nt}}^{\mathrm{FNA}}$) can readily be inverted and becomes proportional to the Gamma distribution of order $N$ and with characteristic time $\tau_{\mathrm{FNA}}$

$$P_{N\mathrm{nt}}^{\mathrm{FNA}}(t) = p_{\mathrm{FNA}}^N \Gamma(t, \tau_{\mathrm{FNA}}, N), \qquad \Gamma(t, \tau_{\mathrm{FNA}}, N) = \frac{1}{\tau_{\mathrm{FNA}}}\frac{(t/\tau_{\mathrm{FNA}})^{N-1}e^{-t/\tau_{\mathrm{FNA}}}}{(N-1)!} \quad \text{(S2)}$$

This distribution has the peak positioned at $T_{\mathrm{FNA}} = N\tau_{\mathrm{FNA}}$, which covers the single dominant FNA peak observed for short dwell-times in absence of nsp13-helicase (**Fig. 1E**).

*Approximation capturing the dominant behaviours at intermediate timescales*

With a separation of timescales we did not need to include the effects of short timescales when considering the behaviour around a longer timescale. This corresponds to the statement that a slowly varying distribution is not much effected by being convoluted by a (comparatively) narrow distribution. For the purpose of a fit-function that captures the dominant behaviour at each timescale by setting sub-dominant timescales to zero: $\tau_{\mathrm{FNA}} = 0$ in $\psi_{N\mathrm{nt}}^{\mathrm{FNA}}$ and $\tau_{\mathrm{FNA}} = \tau_{\mathrm{SNA}} = 0$ in $\psi_{N\mathrm{nt}}^{\mathrm{SNA}}$, resulting in

$$\psi_{N\mathrm{nt}}^{\mathrm{SNA}} \approx \sum_{n_0=0}^{N-1} \binom{N}{n_0, N-n_0} p_{\mathrm{FNA}}^{n_0} p_{\mathrm{SNA}}^{N-n_0} \left(\frac{1}{1+\tau_{\mathrm{SNA}} s}\right)^{N-n_0}$$

$$\psi_{N\mathrm{nt}}^{\mathrm{VSNA}} \approx \sum_{n_0=0}^{N-1} \sum_{n_1=0}^{N-n_0-1} \binom{N}{n_0, n_1, N-n_0-n_1} p_{\mathrm{FNA}}^{n_0} p_{\mathrm{SNA}}^{n_1} p_{\mathrm{VSNA}}^{N-n_0-n_1} \left(\frac{1}{1+\tau_{\mathrm{VSNA}} s}\right)^{N-n_0-n_1} . \tag{S3}$$

Considering that the two shoulders observed for intermediate timescales in the empirical dwell-time distributions are well-fitted by exponential distributions (**Fig. 1E**), we further approximated $P_{N\mathrm{nt}}^{\mathrm{SNA}}$ and third $P_{N\mathrm{nt}}^{\mathrm{VSNA}}$ with single exponential distributions

$$P_{N\mathrm{nt}}^{\mathrm{NA}}(t) \approx \frac{q_{\mathrm{NA}}}{T_{\mathrm{NA}}} e^{-\frac{t}{T_{\mathrm{NA}}}}, \qquad \mathrm{NA} \in \{\mathrm{SNA}, \mathrm{VSNA}\} \tag{S4}$$

that capture the total probabilities $q_{\mathrm{NA}}$ and the average time $T_{\mathrm{NA}}$ of each term. From the definition of the Laplace transform it follows that the total probabilities and average timescales then are

$$q_{\mathrm{NA}} = \int_0^\infty \mathrm{d}t\, P_{N\mathrm{nt}}^{\mathrm{NA}}(t) = \psi_{N\mathrm{nt}}^{\mathrm{NA}}(0)\,; \quad T_{\mathrm{NA}} = \frac{\int_0^\infty \mathrm{d}t\, t\, P_{N\mathrm{nt}}^{\mathrm{NA}}(t)}{\int_0^\infty \mathrm{d}t\, P_{N\mathrm{nt}}^{\mathrm{NA}}(t)} = -\partial_s \ln \psi_{N\mathrm{nt}}^{\mathrm{NA}}(0)\,, \qquad \mathrm{NA} \in \{\mathrm{SNA}, \mathrm{VSNA}\}. \tag{S5}$$

The total probabilities and timescales over a dwell-time window were then related to the single-nucleotide step probabilities and timescales through

$$\begin{aligned}
q_{\mathrm{FNA}} &= p_{\mathrm{FNA}}^{N} \\
q_{\mathrm{SNA}} &= (p_{\mathrm{FNA}} + p_{\mathrm{SNA}})^N - p_{\mathrm{FNA}}^N\,, \\
T_{\mathrm{SNA}} &= \frac{N p_{\mathrm{SNA}} (p_{\mathrm{FNA}} + p_{\mathrm{SNA}})^{N-1}}{(p_{\mathrm{FNA}} + p_{\mathrm{SNA}})^N - p_{\mathrm{FNA}}^N} \tau_{\mathrm{SNA}}\,, \\
q_{\mathrm{VSNA}} &= (p_{\mathrm{FNA}} + p_{\mathrm{SNA}} + p_{\mathrm{VSNA}})^N - (p_{\mathrm{FNA}} + p_{\mathrm{SNA}})^N, \\
T_{\mathrm{VSNA}} &= \frac{N p_{\mathrm{VSNA}} (p_{\mathrm{FNA}} + p_{\mathrm{SNA}} + p_{\mathrm{VSNA}})^{N-1}}{(p_{\mathrm{FNA}} + p_{\mathrm{SNA}} + p_{\mathrm{VSNA}})^N - (p_{\mathrm{FNA}} + p_{\mathrm{SNA}})^N} \tau_{\mathrm{VSNA}}
\end{aligned} \tag{S6}$$

*Short time cut-off due to dwell-time window size*

The approximation of the dwell-time distribution, derived in **Eq. S1-S6**, is constructed to capture the dominant behaviour around $T_{\mathrm{SNA}}$ and $T_{\mathrm{VSNA}}$, and breaks down for $t < T_{\mathrm{FNA}}$ and $t < T_{\mathrm{SNA}}$ respectively. The short time behaviour still grows no faster than $\sim t^{N-1}$ since $N$ sequential steps always need to be taken to get through the dwell-time window. This behaviour will always be dominated by the FNA peak, and we ensured this also in our approximation by multiplying the exponential functions with

$$Q(t/T_{\mathrm{FNA}}) = \frac{(t/T_{\mathrm{FNA}})^{N-1}}{1 + (t/T_{\mathrm{FNA}})^{N-1}} \tag{S7}$$

to ensure the appropriate short time behaviour. To keep the proper normalization from $T_{\mathrm{FNA}}$ we took

$$P_{N\mathrm{nt}}^{\mathrm{SNA}}(t) \approx Q(t/T_{\mathrm{FNA}}) \frac{q_{\mathrm{SNA}}}{T_{\mathrm{SNA}}} e^{-\frac{t-T_{\mathrm{FNA}}}{T_{\mathrm{SNA}}}}, \qquad P_{N\mathrm{nt}}^{\mathrm{VSNA}}(t) \approx Q(t/T_{\mathrm{FNA}}) \frac{q_{\mathrm{VSNA}}}{T_{\mathrm{VSNA}}} e^{-\frac{t-T_{\mathrm{FNA}}}{T_{\mathrm{VSNA}}}}. \tag{S8}$$

*Fit function covering short and intermediate timescales*

We arrived at the fit-function that appropriately captures the dominant behaviour for short and intermediate timescales

$$P_{N\mathrm{nt}}^{\mathrm{s\&i}}(t) \approx \frac{q_{\mathrm{FNA}}N}{T_{\mathrm{FNA}}(N-1)!}\left({}^{tN}/_{T_{\mathrm{FNA}}}\right)^{N-1} e^{-tN/T_{\mathrm{FNA}}} + Q(t/T_{\mathrm{FNA}})\left(\frac{q_{\mathrm{SNA}}}{T_{\mathrm{SNA}}}e^{-\frac{t-T_{\mathrm{FNA}}}{T_{\mathrm{SNA}}}} + \frac{q_{\mathrm{VSNA}}}{T_{\mathrm{VSNA}}}e^{-\frac{t-T_{\mathrm{FNA}}}{T_{\mathrm{VSNA}}}}\right). \quad \text{(S9)}$$

Under the assumption that $T_{\mathrm{FNA}} \ll T_{\mathrm{SNA}} \ll T_{\mathrm{VSNA}}$ the corrections ensuring normalization introduced above were very small, but as maximum-likelihood estimation maximizes the total probability of dwell-times it is sensitive to normalization and we choose to strictly enforce it.

*Accounting for long-time pauses*

In addition to a peak and two shoulders, a tail of long dwell-times is observed in the empirical dwell-time distributions for elongation dynamics by the core RTC with a decay of order $\sim t^{-3/2}$ over orders of magnitude (10-1000 s) (**Fig. 3B**). This trend suggests we observed long-lived pauses resulting from random walks that are recovered, such as backtracking (Depken et al. 2009). The first-passage time of backtrack recovery represents the time it takes to re-align with the 3' end of the template after a random walk on the RNA and re-initiate RNA synthesis. Defining the stepping rate towards realignment as $k_{\mathrm{f}}$ (forward) and away from realignment as $k_{\mathrm{b}}$ (backward), the first passage time distribution for such a random walk can be split into three regimes (Depken et al. 2009): For the low-time regime $t \ll t_1 = 1/\sqrt{k_{\mathrm{f}}k_{\mathrm{b}}}$, the distribution is flat; in the intermediate-time regime $t_1 \ll t \ll t_2 = 1/\left(\sqrt{k_{\mathrm{f}}} - \sqrt{k_{\mathrm{b}}}\right)^2$ the distribution decays as a power law $\sim t^{-3/2}$; and for $t \gg t_2$ the distribution is exponentially cut off with characteristic timescale $t_2$.(Depken, Galburt et al. 2009)

We never observed the transition into the short time regime, suggesting that $t_1$ is in a regime where the behaviour is dominated by either of the FNA, SNA or VSNA pathways ($t_1 \ll T_{\mathrm{VSNA}} \approx 5\ s$). We also never observed the exponential cut-off for large timescales, suggesting that $t_2$ is larger than the total measurement time ($t_2 \gg 1000$ s). The fact that the transitions to the short and to the long time regime were not observed means that $t_2/t_1 \gg 200$. If $k_{\mathrm{f}} < k_{\mathrm{b}}$ the average long-lived pause duration and extension would diverge. As we did not see evidence of this, we concluded that $k_{\mathrm{f}} > k_{\mathrm{b}}$. The simplest model for the forward-biased random walk stepping rates dependence on RNA tension ($F$) is that they follow an Arrhenius law

$$k_{\mathrm{f}} = k_0 e^{F\delta/k_B T}, \quad k_{\mathrm{b}} = k_0 e^{-F(d-\delta)/k_B T} \Rightarrow k_{\mathrm{f}}/k_{\mathrm{b}} = e^{Fd/k_B T}, \quad \sqrt{k_{\mathrm{f}}k_{\mathrm{b}}} = k_0 e^{F(\delta - d/2)/k_B T}. \quad \text{(S10)}$$

Using that $t_1 = 1/\sqrt{k_{\mathrm{f}}k_{\mathrm{b}}}$ and $t_2 = 1/\left(\sqrt{k_{\mathrm{f}}} - \sqrt{k_{\mathrm{b}}}\right)^2$, we see that $\mathrm{t}_2/\mathrm{t}_1 \gg 200$ means $1 < k_{\mathrm{f}}/k_{\mathrm{b}} < 1.2$. The fact that the ratio between stepping rates never exceeded 1.2 even for the highest tension (40 pN) implies that $d < 0.02$ nm. If we assumed the distance to the transition state $\delta$ lies withing the step ($0 < \delta < d$), this would in turn imply that $t_1$ only appreciably shifts for forces above $\frac{k_{\mathrm{B}}T}{0.02\ \mathrm{nm}/2} \approx 400$ pN. For our empirical conditions we should therefore be safe assuming that $t_1$ does not depend appreciably on force, and that ratios of amplitudes faithfully report on ratios of probabilities.

The upper limit for $d$ is significantly smaller than the length change from double-stranded to single-stranded RNA per nucleotide ($a \approx 0.2$ nm), indicating that the energy landscape for the long-lived pause is rugged and a full nucleotide step (0.2 nm) could consist of many intermediate shorter steps before the translocation is complete

(Grossman-Haham, Rosenblum et al. 2018). As a result, we could not say how far the backtrack of the RTC goes during the long-lived pause and it could even stay on the same position while going through many conformations with average distance $d < 0.02$ nm.

For the purpose of capturing the power-law decay in the long time regime and the sub-dominance at shorter timescales, the first-passage time distribution of the long-lived pause (LLP) recovery is approximated by a power law decay with amplitude $q_{\text{LLP}}$, cut-off at short timescales (Depken, Galburt et al. 2009).

$$P_{N\text{nt}}^{\text{LLP}}(t) = \frac{q_{\text{LLP}}}{(1 + t/t_1)^{3/2}} \tag{S11}$$

The amplitude $q_{\text{LLP}}$ of the power law decay is proportional to the total probability for entering the long-lived pause in a dwell-time window, but the constant of proportionality is sensitive to the undetermined cut-off time $t_1$. As long as $t_1$ does not change, ratios of amplitudes correspond to ratios of probabilities, so we treated $q_{\text{LLP}}$ as a relative probability.

As for the other intermediate time processes we accounted for the fact that a dwell-time window consists of 10 nt and the contribution of long-lived pauses at short times must be cut-off with a regularization function (**Eq. S7**). Properly normalizing the contribution to exclude times shorter than $T_{\text{FNA}}$ we arrived at the dwell-time fit-function for the core RTC dynamics alternating (alt) between NA pathways and long-lived pauses (LLPs)

$$P_{N\text{nt}}^{\text{alt}}(t, \boldsymbol{p}) \approx P_{N\text{nt}}^{\text{s\&i}}(t) + Q\left(\frac{t}{T_{\text{FNA}}}\right)\frac{1}{2}\sqrt{1 + \frac{T_{\text{FNA}}}{t_1}}P_{\text{LLP}}(t). \tag{S12}$$

Here we have introduced the parameter vector $\boldsymbol{p} = \{q_{\text{FNA}}, q_{\text{SNA}}, q_{\text{VSNA}}, q_{\text{LLP}}, T_{\text{FNA}}, T_{\text{SNA}}, T_{\text{VSNA}}\}$ for later convenience. The weights satisfy $q_{\text{FNA}} + q_{\text{SNA}} + q_{\text{VSNA}} + q_{\text{LLP}} = 1$.

As we could not obtain the actual value of $t_1$ from our data, but the value also does not affect the shape of the long-lived pause recovery distribution where it is dominant, we simply set $t_1 = 1$ s, but recognized that $q_{\text{LLP}}$ cannot be interpreted as a probability other than that ratios of amplitudes correspond to ratios of probabilities as described above.

The expression for $P_{N\text{nt}}^{\text{core}}(t, \boldsymbol{p})$ is strictly only valid for observations that are not limited by the measurement time or resolution. Since the resolution $t_{\text{cut}} = 0.08$ s and measurement time $t_{\max} = 1000$ s were finite, the dwell-time fit-function was normalized to this interval for the purpose of the fits (Bera, Seifert et al. 2021)

$$P_{N\text{nt}}^{\text{core fit}}(t, \boldsymbol{p}) \approx \frac{P_{N\text{nt}}^{\text{alt}}(t, \boldsymbol{p})}{\int_{t_{\text{cut}}}^{t_{\max}} \mathrm{d}\tau P_{N\text{nt}}^{\text{alt}}(\tau, \boldsymbol{p})}\,. \tag{S13}$$

#### Section S1.3: Dwell-time fit-function for saturating nsp13-helicase concentrations

The dwell-time fit-function **Eq. S13** including the FNA, SNA, VSNA pathway and long-lived pauses fitted well to the empirical dwell-time distribution for core RTC elongation dynamics (**Fig. 1E**), so we set out to extend it in presence of nsp13-helicase.

For saturating concentrations of nsp13-helicase, we observed bursts of very fast nucleotide addition (VFNA), in addition to the nucleotide addition signatures observed for the core RTC (**Fig. 1D**, **Fig. 3A**). These bursts of VFNA resulted in a peak for very short dwell-times (**Fig. 1DE**). Since the bursts last longer than the dwell-time window

(**Fig. 1C**), the VFNA pathway was added to the dwell-time fit-function as a separate gamma pdf with $N$ steps and timescale $\tau_{\mathrm{VFNA}}$. After addition of the VFNA peak, the unnormalized fit-function reads

$$P_{N\mathrm{nt}}^{\mathrm{sat}}(t, \boldsymbol{p}^{\mathbf{sat}}, f_{\mathrm{VFNA}}^{\mathrm{sat}}, T_{\mathrm{VFNA}}^{\mathrm{sat}}) \approx f_{\mathrm{VFNA}}^{\mathrm{sat}} \Gamma(t, T_{\mathrm{VFNA}}/N, N) + (1 - f_{\mathrm{VFNA}}^{\mathrm{sat}}) P_{N\mathrm{nt}}^{\mathrm{alt}}(t, \boldsymbol{p}^{\mathbf{sat}}) \tag{S14}$$

The first part of this distribution is a gamma distribution with peak at $T_{\mathrm{VFNA}} = N\tau_{\mathrm{VFNA}}$, where $\tau_{\mathrm{VFNA}}$ is the single nucleotide addition rate. Normalization to the observational interval gave us the final dwell-time fit-function at saturating concentrations of nsp13-helicase

$$P_{N\mathrm{nt}}^{\mathrm{fit}}(t, \boldsymbol{p}^{\mathbf{sat}}, f_{\mathrm{VFNA}}^{\mathrm{sat}}, T_{\mathrm{VFNA}}^{\mathrm{sat}}) \approx \frac{P_{N\mathrm{nt}}^{\mathrm{sat}}(t, \boldsymbol{p}^{\mathbf{sat}}, f_{\mathrm{VFNA}}^{\mathrm{sat}}, T_{\mathrm{VFNA}}^{\mathrm{sat}})}{\int_{t_{\mathrm{cut}}}^{t_{\mathrm{max}}} \mathrm{d}\tau P_{N\mathrm{nt}}^{\mathrm{sat}}(\tau, \boldsymbol{p}^{\mathbf{sat}}, f_{\mathrm{VFNA}}^{\mathrm{sat}}, T_{\mathrm{VFNA}}^{\mathrm{sat}})}. \tag{S15}$$

#### Section S1.4: Dwell-time fit-function for non-saturating nsp13-helicase concentrations

While our dwell-time fit-function was constructed for RTC-nsp13 complexes at saturating nsp13-helicase concentrations, titration of the nsp13-helicase concentration leaded to a mixing of timescales pertaining to the different RTC-nsp13 complexes. Still, the dwell-time distributions at non-saturating nsp13-helicase concentrations were well-captured by the fit-function constructed for saturating nsp13-helicase conditions (**Fig. 3A**), indicating that the variation in timescales for a particular process remained small compared to differences in timescales between processes. We therefore used the dwell-time fit-function for saturating nsp13-helicase concentrations also for non-saturating nsp13-helicase concentrations, but with parameters depending on the concentration

$$P_{N\mathrm{nt}}^{\mathrm{fit}}\left(t, \boldsymbol{p}^{[\mathbf{nsp13}]}, f_{\mathrm{VFNA}}^{[\mathrm{nsp13}]}, T_{\mathrm{VFNA}}^{[\mathrm{nsp13}]}\right). \tag{S16}$$

The timescales and probabilities of the slow to intermediate pathways $\boldsymbol{p}^{[\mathbf{nsp13}]}$ were extracted as averages over the complexes involved. How these averages should be performed is explained in **Section S2.2**.

#### Section S1.5: Settings parameters constant

We started by performing fits on various nsp13-helicase concentrations with all parameters free. This resulted in good fits (**Fig. S8**), but no significant trend was observed in the VFNA and VSNA characteristic timescale (**Fig. S2AEI**). A constant $T_{\mathrm{VFNA}}$ is consistent with each burst arising from a single RTC complex. The lack of any significant trend in the VSNA characteristic timescale could simply be due to that the region where VSNA dominates was generally very small, and the characteristic timescale was hard to determine (**Fig. S9**). To reduce overfitting we fixed the VFNA characteristic timescale (0.2 s) and the VSNA characteristic timescale (4.8 s) without apparent reduction in goodness-of-fit (**Fig. S9**) or shifts in the other parameters (**Fig. S2**). For consistency, we also put the VSNA characteristic timescale fixed in the fits for RTC elongation dynamics on ssRNA (**Fig. S5F**).

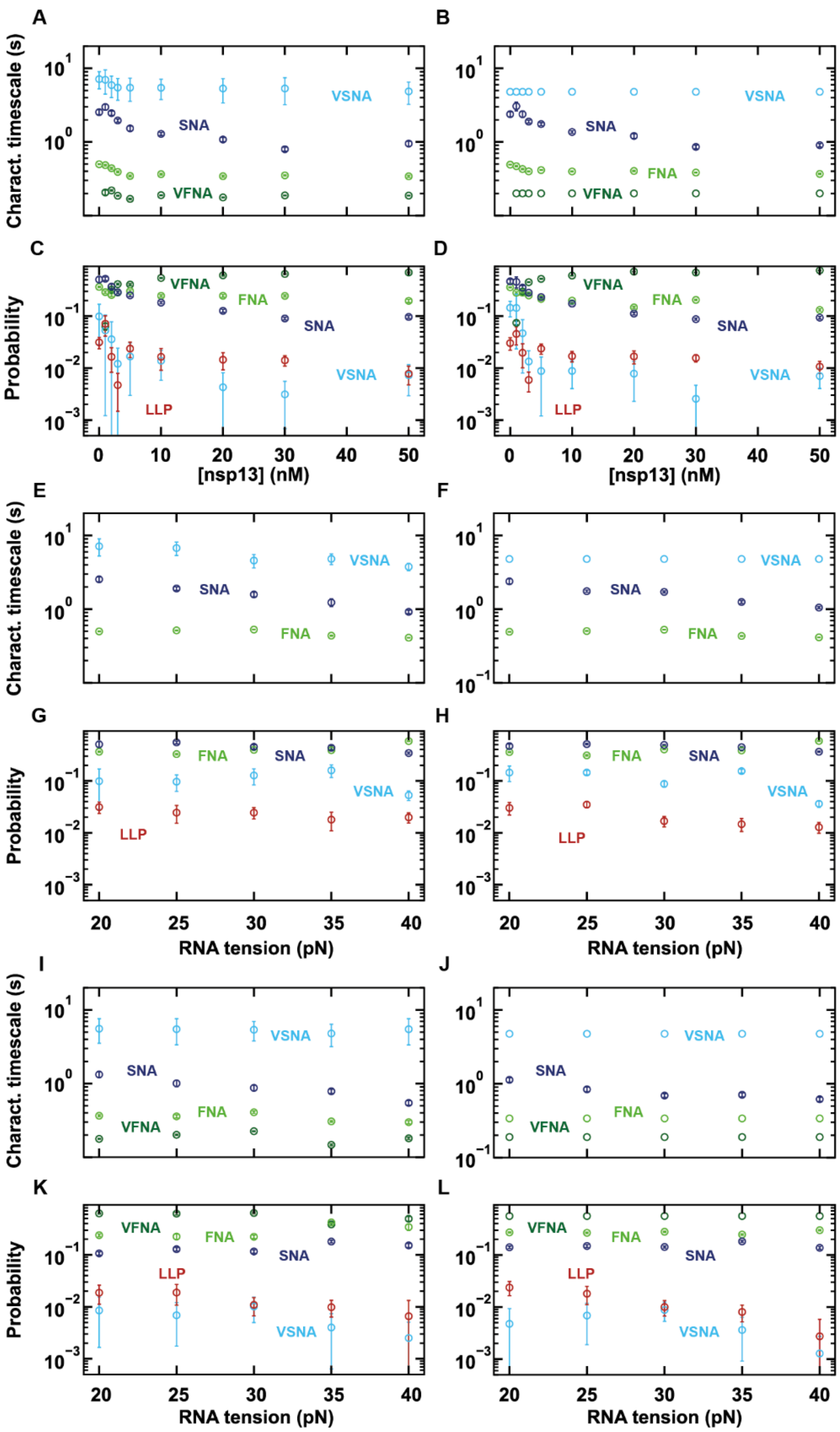

**Fig. S2. Parameters from the dwell-time fits with all parameters free or the VFNA and VSNA timescales fixed for RTC elongation dynamics on a dsRNA template. (A, B)** Characteristic timescales (s) and **(C, D)** Probabilities for RTC elongation dynamics versus nsp13-helicase concentration (nM) from the dwell-time fits with all parameters free (A, C) or with the VFNA and VSNA characteristic timescale fixed to 0.2 s and 4.8 s respectively (B, D). **(E, F)** Characteristic timescales (s) and **(G, H)** Probabilities for core RTC elongation dynamics versus RNA tension (pN) from the dwell-time fits with all parameters free (E, G) or with the VSNA characteristic timescale fixed to 0.2 s and 4.8 s respectively (F, H). **(I, J)** Characteristic timescales (s) and **(K, L)** Probabilities for RTC elongation dynamics versus RNA tension (pN) at saturating nsp13-helicase concentration from the dwell-time fits with all parameters free (I, K) or with the VFNA and VSNA characteristic timescale fixed to 0.2 s and 4.8 s respectively (J, L). The error bars shown for the fitted parameters represent one standard deviation obtained from 100 bootstrap fits.

### Section S2: Connecting the assembly of the RTC-nsp13 complex with the elongation dynamics

#### Section S2.1: Assembly of the RTC-nsp13 complex

We observed no bursts for opening of the dsRNA for Poliovirus RdRP with nsp13-helicase in the reaction mixture at high concentration ([nsp13]=50 nM) (**Fig. 2A, Fig. S2ACD**). This suggested that the bursts observed for CoV RTC in the presence of nsp13-helicase are the result of complex formation and not due to nsp13-helicase interacting directly with the RNA to unwind the RNA duplex. There was also a slowdown of RTC elongation dynamics upon introduction of an ATPase-dead mutant of nsp13-helicase (**Fig. 2B, Fig. S2BCD**), confirming that nsp13-helicase forms a complex with the core RTC. For RTC elongation on a ssRNA template in presence of active nsp13-helicase, we observed significantly faster overall dynamics, but no bursts of nucleotide addition (**Fig. 2C**). This absence of bursts supports the conclusion that the VFNA pathway is the result of an interaction of nsp13-helicase with the non-template RNA strand (**Fig. 2D**). Taken together, these results showed that nsp13-helicase associates with the RTC and engages with the non-template strand to produce elongation bursts (**Fig. 3B**).

The fact that in addition to the VFNA peak the signatures of all other processes remain apparent also in the dwell-time distributions at saturating nsp13-helicase concentrations (**Fig. 3A**) suggested that the nsp13-RTC complex can exist in two forms: an engaged nsp13-RTC complex, which results in the VFNA pathway, and a non-engaged nsp13-bound RTC which have the same nucleotide addition pathways as the core RTC. Cryo-EM structural studies (Chen et al. 2020, Chen et al. 2022 and Yan et al. 2022) have shown that nsp13.1 first binds to the core RTC and a conformational change has to occur before nsp13.2 can bind. Moreover, the structural studies found that nsp13.1 is interacting with the template RNA strand, which implies that nsp13.2 is interacting with the non-template RNA strand (**Fig. 3B**).

To establish if the RTC needs one or two nsp13-helicases to engage with the non-template strand in the VFNA pathway, we tested two RTC-nsp13 assembly models (**Fig. S4A**) on our data.

**Assembly model 1:** nsp13.1 binds to the core RTC (RTC-nsp13.1) and engages with the non-template RNA strand (RTC-nsp13.1*) enabling elongation bursts.

**Assembly model 2:** nsp13.1 binds to the core RTC (RTC-nsp13.1), nsp13.2 binds to RTC-nsp13.1 (RTC-nsp13.1,2) and nsp13.2 engages with the non-template RNA strand (RTC-nsp13.1,2*) enabling elongation bursts.

To discriminate between assembly models, we assumed the binding of two nsp13-helicases is in dynamic equilibrium, but they do not exchange within one dwell-time window. We use Boltzmann statistics to calculate the fractional occupancies of the RTC complexes for our two models:

For **Assembly model 1** the fractional occupancies of each RTC complex were calculated as

$$p_{\mathrm{c}}([\mathrm{nsp13}]) = \frac{1}{Z([\mathrm{nsp13}])}, \qquad p_1([\mathrm{nsp13}]) = \frac{[\mathrm{nsp13}]e^{-\Delta G_{\mathrm{c}\to 1}}}{Z([\mathrm{nsp13}])}, \qquad p_{1^*}([\mathrm{nsp13}]) = \frac{[\mathrm{nsp13}]e^{-(\Delta G_{\mathrm{c}\to 1} + \Delta G_{1\to 1^*})}}{Z([\mathrm{nsp13}])}, \tag{S17}$$

with c representing the core RTC, 1 the complex with one nsp13, but no engagement, and 1* the complex with engagement of the first nsp13. We have also introduced the normalization constant $Z([\mathrm{nsp13}]) = [\mathrm{nsp13}]e^{-(\Delta G_{\mathrm{c}\to 1} + \Delta G_{1\to 1^*})} + [\mathrm{nsp13}]e^{-\Delta G_{\mathrm{c}\to 1}} + 1$, with $\Delta G_{\mathrm{cpx}\to\mathrm{cpx}'}$ representing the free-energies differences between the sequential RTC complexes $\mathrm{cpx} \in \{\mathrm{c}, 1\}$ and $\mathrm{cpx}' \in \{1, 1^*\}$ with 1 nM as reference concentration, expressed in units of $\mathrm{k_B T}$.

For **Assembly model 2** the fractional occupancies of each RTC complex were calculated as

$$p_{\mathrm{c}}([\mathrm{nsp13}]) = \frac{1}{Z([\mathrm{nsp13}])}, \quad p_1([\mathrm{nsp13}]) = \frac{[\mathrm{nsp13}]e^{-\Delta G_{\mathrm{c}\to 1}}}{Z([\mathrm{nsp13}])},$$
$$p_2([\mathrm{nsp13}]) = \frac{[\mathrm{nsp13}]^2 e^{-(\Delta G_{\mathrm{c}\to 1} + \Delta G_{1\to 2})}}{Z([\mathrm{nsp13}])}, \quad p_{2^*}([\mathrm{nsp13}]) = \frac{[\mathrm{nsp13}]^2 e^{-(\Delta G_{\mathrm{c}\to 1} + \Delta G_{1\to 2} + \Delta G_{2\to 2^*})}}{Z([\mathrm{nsp13}])}, \tag{S18}$$

with c representing the core RTC, 1 the complex with one nsp13, 2 the complex with two nsp13-helicases but no engagement, and 2* the complex with two nsp13-helicases and engagement of the second nsp13. We have also introduced the normalization constant $Z([\mathrm{nsp13}]) = [\mathrm{nsp13}]^2 e^{-(\Delta G_{\mathrm{c}\to 1} + \Delta G_{1\to 2} + \Delta G_{2\to 2^*})} + [\mathrm{nsp13}]^2 e^{-(\Delta G_{\mathrm{c}\to 1} + \Delta G_{1\to 2})} + [\mathrm{nsp13}]e^{-\Delta G_{\mathrm{c}\to 1}} + 1$, with $\Delta G_{\mathrm{cpx}\to\mathrm{cpx}'}$ representing the free-energies differences between the sequential RTC complexes $\mathrm{cpx} \in \{\mathrm{c}, 1, 2\}$ and $\mathrm{cpx}' \in \{1, 2, 2^*\}$ with 1 nM as reference concentration, expressed in units of $\mathrm{k_B T}$.

We performed fits of the probability to be in an nsp13-engaged RTC state for model 1 and 2 ($p_{1^*}$ and $p_{2^*}$ respectively) to the fractional occupancy $f_{\mathrm{FNA}}^{[\mathrm{nsp13}]}$ versus nsp13-helicase concentration by minimizing the error-weighted square deviation (**Fig. S4B**). We concluded that Assembly model 2 fits better, which is largely due to its quadratic dependence of $p_{2^*}$ on nsp13-helicase concentration at low concentrations (**Eq. S18**).

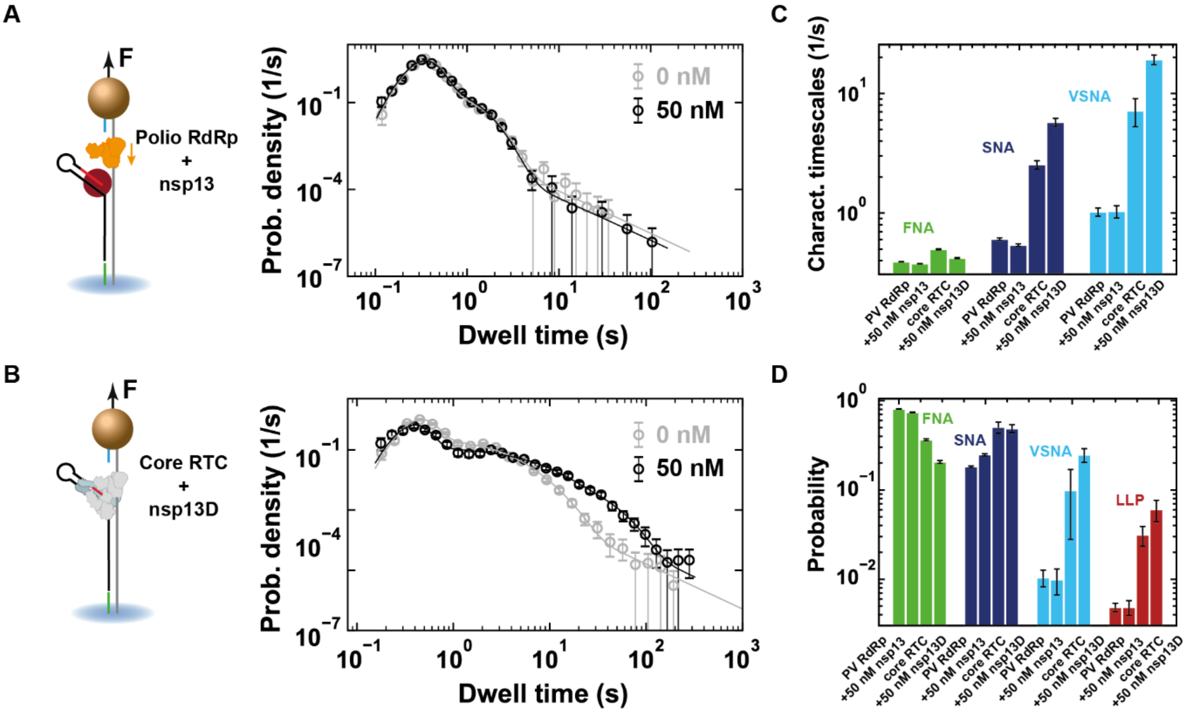

**Fig. S3. Nsp13-helicase specifically associates with the SARS-CoV-2 core RTC and nsp13.2 assists elongation dynamics by engaging with the non-template RNA strand. (A, B)** On the left, schematics describing the measurements performed with nsp13-helicase using either (A) Poliovirus RdRp or (B) SARS-CoV-2 core RTC elongating on a dsRNA template with wild-type nsp13-helicase (nsp13) (A) or ATPase dead nsp13-helicase mutant (nsp13D) (B) respectively. On the right the corresponding dwell-time distributions for elongation dynamics are shown without (grey) or with (black) saturating concentrations of nsp13-helicase (50 nM). **(C, D)** Bar plots of the characteristic timescales (C) and probabilities (D) obtained from the dwell-time fits.

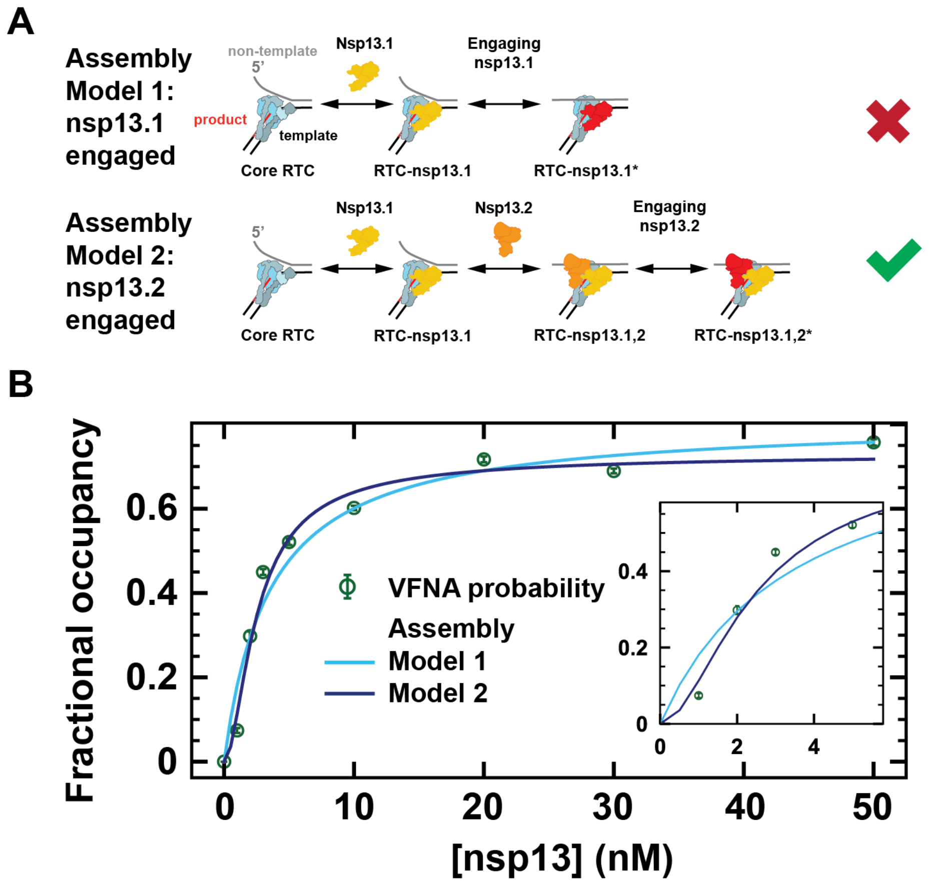

**Fig. S4. Comparison of RTC-nsp13 assembly models shows that nsp13.2 engages with the non-template RNA strand. (A)** Schematic representation of the assembly models tested. In Assembly model 1, nsp13.1 binds first to the RTC (RTC-nsp13.1) and engages with the non-template RNA (RTC-nsp13.1*) making the very fast nucleotide addition pathway. Note that the binding of nsp13.2 is irrelevant in this model, since it is not involved in the engagement of nsp13.1. In Assembly model 2, nsp13.1 binds first, followed by nsp13.2 and nsp13.2 then engages with the non-template RNA (RTC-nsp13.1,2*). From the fits to the VFNA probability (B) we conclude that Assembly model 2 is the correct model. **(B)** Fits with the fractional occupancy of the nsp13-engaged state to the VFNA probability versus nsp13-helicase concentration ([nsp13] (nM)) for the two RTC Assembly models compared. (sky blue line) Fractional occupancy of the RTC-nsp13.1* complex versus nsp13-helicase concentration for Assembly model 1. (dark blue line) Fractional occupancy of the RTC-nsp13.1,2* complex versus nsp13-helicase concentration for Assembly model 2. (green circles with error bars) The mean and standard deviation of the VFNA probability obtained from 100 dwell-time bootstrap fits versus nsp13-helicase concentration with the VFNA and VSNA characteristic timescale fixed.

#### Section S2.2: Effective parameters for elongation dynamics as a function of nsp13-helicase concentration

At intermediate nsp13-helicase concentrations, dwell-time distributions shift based on the stoichiometry of the various RTC complexes, and not per se due to a shift in the elongation dynamics by individual complexes. Since the complexes have fractional occupancies as in **Eq. S18**, the shift in the dwell-time distribution is captured by

$$P^{\text{mix}}_{N\text{nt}}(t) = p_{2^*}([\text{nsp13}])\Gamma(t, T_{\text{VFNA}}/N, N) + \sum_{\text{cpx}\in C} p_{\text{cpx}}([\text{nsp13}])P^{\text{alt}}_{N\text{nt}}(t, \boldsymbol{p}^{\text{cpx}}), \qquad \text{cpx} \in \{\text{c}, 1, 2\} \tag{S19}$$

For a mix of complexes we observe that **Eq. S16** still captures the data (**Fig. 3A**). From this we took that the variations in characteristic timescales between models are moderate enough that they can be captured by effective timescales and probabilities. Therefore we replaced the non-bursting part of the above equation with an effective distribution

$$\sum_{\text{cpx}\in C} p_{\text{cpx}}([\text{nsp13}])P^{\text{alt}}_{N\text{nt}}(t, \boldsymbol{p}^{\text{cpx}}) \approx P^{\text{alt}}_{N\text{nt}}(t, \boldsymbol{p}^{\text{eff}}([\text{nsp13}])), \qquad \text{cpx} \in \{\text{c}, 1, 2\} \tag{S20}$$

where the effective probabilities and characteristic times $\boldsymbol{p}^{\text{eff}}([\text{nsp13}])$ are chosen to match the total probability and average characteristic timescale among complexes for each pathway and nsp13-helicase concentration. This resulted in

$$P^{\text{mix,eff}}_{N\text{nt,cpx}}(t) = p_{2^*}([\text{nsp13}])\Gamma(t; T_{\text{VFNA}}/N, N) + P^{\text{alt}}_{N\text{nt}}(t, \boldsymbol{p}^{\text{eff}}([\text{nsp13}])), \qquad \text{cpx} \in \{\text{c}, 1, 2\} \tag{S21}$$

With

$$\begin{aligned} q^{\text{eff}}_{\text{NA}}([\text{nsp13}]) &= \sum_{\text{cpx}\in C} p_{\text{cpx}}([\text{nsp13}])q^{\text{cpx}}_{\text{NA}}, \qquad \text{NA} \in \{\text{FNA}, \text{SNA}, \text{VSNA}\} \\ T^{\text{eff}}_{\text{NA}} &= \frac{\sum_{\text{cpx}\in C} p_{\text{cpx}}([\text{nsp13}])q^{\text{cpx}}_{\text{NA}}\, T^{\text{cpx}}_{\text{NA}}}{\sum_{\text{cpx}\in C} p_{\text{cpx}}([\text{nsp13}])q^{\text{cpx}}_{\text{NA}}}, \qquad \text{NA} \in \{\text{FNA}, \text{SNA}, \text{VSNA}\} \end{aligned} \tag{S22}$$

The probabilities and characteristic timescales for each pathway $\text{NA} \in \{\text{FNA}, \text{SNA}, \text{VSNA}\}$ entered by complex $\text{cpx} \in \{\text{c}, 1, 2\}$ ($q^{\text{cpx}}_{\text{NA}}$ and $T^{\text{cpx}}_{\text{NA}}$) are expressed in terms of single nucleotide probabilities and timescales as shown for the core RTC in **Section S1.2** (**Eq. S6**)

$$\begin{aligned} q^{\text{cpx}}_{\text{FNA}} &= \left(p^{\text{cpx}}_{\text{FNA}}\right)^N \\ q^{\text{cpx}}_{\text{SNA}} &= \left(p^{\text{cpx}}_{\text{FNA}} + p^{\text{cpx}}_{\text{SNA}}\right)^N - \left(p^{\text{cpx}}_{\text{FNA}}\right)^N, \\ T^{\text{cpx}}_{\text{SNA}} &= \frac{N p^{\text{cpx}}_{\text{SNA}}\left(p^{\text{cpx}}_{\text{FNA}} + p^{\text{cpx}}_{\text{SNA}}\right)^{N-1}}{\left(p^{\text{cpx}}_{\text{FNA}} + p^{\text{cpx}}_{\text{SNA}}\right)^N - \left(p^{\text{cpx}}_{\text{FNA}}\right)^N} \tau^{\text{cpx}}_{\text{SNA}}, \\ q^{\text{cpx}}_{\text{VSNA}} &= \left(p^{\text{cpx}}_{\text{FNA}} + p^{\text{cpx}}_{\text{SNA}} + p^{\text{cpx}}_{\text{VSNA}}\right)^N - \left(p^{\text{cpx}}_{\text{FNA}} + p^{\text{cpx}}_{\text{SNA}}\right)^N, \\ T^{\text{cpx}}_{\text{VSNA}} &= \frac{N p^{\text{cpx}}_{\text{VSNA}}\left(p^{\text{cpx}}_{\text{FNA}} + p^{\text{cpx}}_{\text{SNA}} + p^{\text{cpx}}_{\text{VSNA}}\right)^{N-1}}{\left(p^{\text{cpx}}_{\text{FNA}} + p^{\text{cpx}}_{\text{SNA}} + p^{\text{cpx}}_{\text{VSNA}}\right)^N - \left(p^{\text{cpx}}_{\text{FNA}} + p^{\text{cpx}}_{\text{SNA}}\right)^N} \tau^{\text{cpx}}_{\text{VSNA}} \end{aligned} \tag{S23}$$

Since we have $q^{\text{cpx}}_{\text{FNA}} + q^{\text{cpx}}_{\text{SNA}} + q^{\text{cpx}}_{\text{VSNA}} + q^{\text{cpx}}_{\text{LLP}} = 1$, we obtain from eq. S22 that $q^{\text{eff}}_{\text{FNA}} + q^{\text{eff}}_{\text{SNA}} + q^{\text{eff}}_{\text{VSNA}} + q^{\text{eff}}_{\text{LLP}} = 1 - p_{2^*}([\text{nsp13}])$. To perform the fits we normalized the mixed fit function over the empirical time interval

$$P_{N\mathrm{nt}}^{\mathrm{mix,fit}}(t) \approx \frac{P_{N\mathrm{nt}}^{\mathrm{mix,eff}}(t)}{\int_{t_{\mathrm{cut}}}^{t_{\max}} \mathrm{d}\tau P_{N\mathrm{nt}}^{\mathrm{mix,eff}}(\tau)} \tag{S24}$$

With this fit-function, referred to as the RTC Assembly – Elongation dynamics model, we captured the fact that any dependence on nsp13-helicase concentration arises from a shift in stoichiometry between RTC-nsp13 complexes, and not a shift in dynamics for individual complexes.

#### Section S2.3: Non-bursting elongation dynamics at non-saturating nsp13-helicase concentrations

On top of the elongation bursts (VFNA), we observed a significant decrease in the FNA and SNA characteristic timescale for RTC elongation on the dsRNA template, which saturated above 10 nM nsp13-helicase (**Fig. 3C**). This is supported by a decrease observed in the FNA characteristic timescale for RTC elongation on ssRNA saturating only for high nsp13-helicase concentration (20 nM) (**Fig. S5F**), indicating that the effect resulted from nsp13.2 association and impacts RTC-nsp13.1,2 elongation dynamics. Furthermore, in presence of saturating concentration of ATPase-dead nsp13-helicase (20 nM) we observed a significant decrease in the FNA characteristic timescale, next to an increase in the slower pathway timescales and probabilities (SNA, VSNA and LLP) (**Fig. S5FG**), indicating that this effect was allosteric.

Based on these observations, we performed a global fit to the dwell-time distributions for all nsp13-helicase concentrations measured with different single nucleotide probabilities and timescales for the RTC-nsp13.1,2 complex than the core RTC and RTC-nsp13.1 complex ($p_{\mathrm{NA}}^{1} = p_{\mathrm{NA}}^{\mathrm{c}}, \tau_{\mathrm{NA}}^{1} = \tau_{\mathrm{NA}}^{\mathrm{c}}$ with $\mathrm{NA} \in \{\mathrm{FNA, SNA, VSNA}\}$), referred to as the RTC Assembly – Elongation dynamics model (**Fig S6**). This resulted in a good fit (**Fig. S10**) that captured the trends in the dwell-time fit parameters for all pathways (**Fig. 3CD**).

To make sure that the changes in single nucleotide probabilities and timescales for the RTC-nsp13.1,2 complex are significant, we tested if the dwell-time distributions could be captured well if we keep the single nucleotide probabilities and characteristic timescales fixed to the values obtained from the dwell-time fits to the core RTC data ($p_{\mathrm{NA}}^{\mathrm{cpx}} = p_{\mathrm{NA}}^{\mathrm{c}}, \tau_{\mathrm{NA}}^{\mathrm{cpx}} = \tau_{\mathrm{NA}}^{\mathrm{c}}$ with $\mathrm{cpx} \in \{1,2\}$ and $\mathrm{NA} \in \{\mathrm{FNA, SNA, VSNA}\}$) or if we let only the single nucleotide probabilities for the RTC-nsp13.1,2 complex free in the fit ($p_{\mathrm{NA}}^{1} = p_{\mathrm{NA}}^{\mathrm{c}}, \tau_{\mathrm{NA}}^{\mathrm{cpx}} = \tau_{\mathrm{NA}}^{\mathrm{c}}$ with $\mathrm{cpx} \in \{1,2\}$ and $\mathrm{NA} \in \{\mathrm{FNA, SNA, VSNA}\}$). Both simplifications did not result in a good fit (**Fig. S11 & S12**), showing that both probabilities and timescales of the NA pathways change significantly for the RTC-nsp13.1,2 complex w.r.t. the core RTC, which was confirmed by 100 bootstrap fits (**Table S2**).

Furthermore, we needed to check whether taking the effective characteristic timescale and probabilities for each pathway does not significantly change the probability density function (pdf) and thus the fitted parameters (**Section S2.2**). In order to do this, we substituted the fitted single nucleotide parameters in the total first passage time distributions for the mix of RTC complexes (**Eq. S19**) and compared the resulting pdf curves to the empirical dwell-time distributions. The resulting pdf are within one standard deviation of the empirical dwell-time distributions for most conditions (**Fig. S13**), so we concluded that taking the effective parameters for elongation dynamics does not affect the pdf curves significantly for the conditions measured.

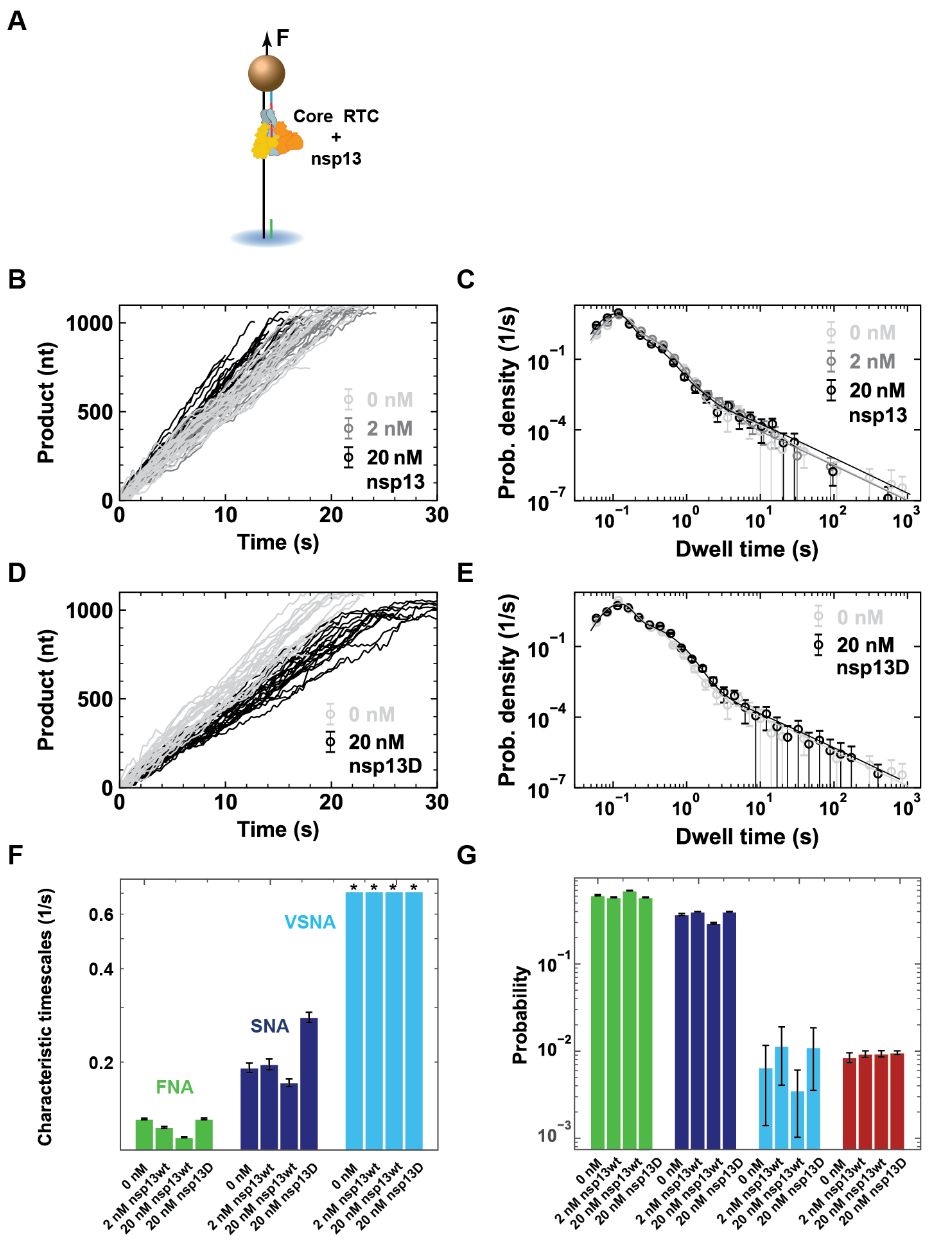

**Fig. S5. Comparison of RTC elongation dynamics on an ssRNA template. (A)** Schematic of core RTC with associated nsp13-helicases elongating on an ssRNA template. **(B)** Elongation traces obtained on an ssRNA template for increasing nsp13-helicase concentrations (0 nM light grey, 2 nM grey, 20 nM black) at constant RNA tension (25 pN). **(C)** Dwell-time distributions of RTC elongation dynamics on an ssRNA template for increasing nsp13-helicase concentrations (0 nM light grey, 2 nM grey, 20 nM black). The mean (circles) and error-bars are shown for 100 bootstraps fits with their respective dwell-time fits (lines). **(D)** Elongation traces obtained on an ssRNA template with or without ATPase-dead (D) nsp13 (0 nM light grey, 20 nM black) at constant RNA tension (25 pN). **(E)** Dwell-time distributions of RTC elongation dynamics on an ssRNA template with or without nsp13D (0 nM light grey, 20 nM black). The mean (circles) and error-bars are shown for 100 bootstraps fits with their respective dwell-time fits (lines). **(F)** Characteristic timescales (s) and **(G)** Probabilities from the dwell-time fits for increasing nsp13-helicase concentrations (0 nM, 2 nM, 20 nM) and saturating nsp13D condition (20 nM). (*) VSNA characteristic timescale is fixed in the fits.

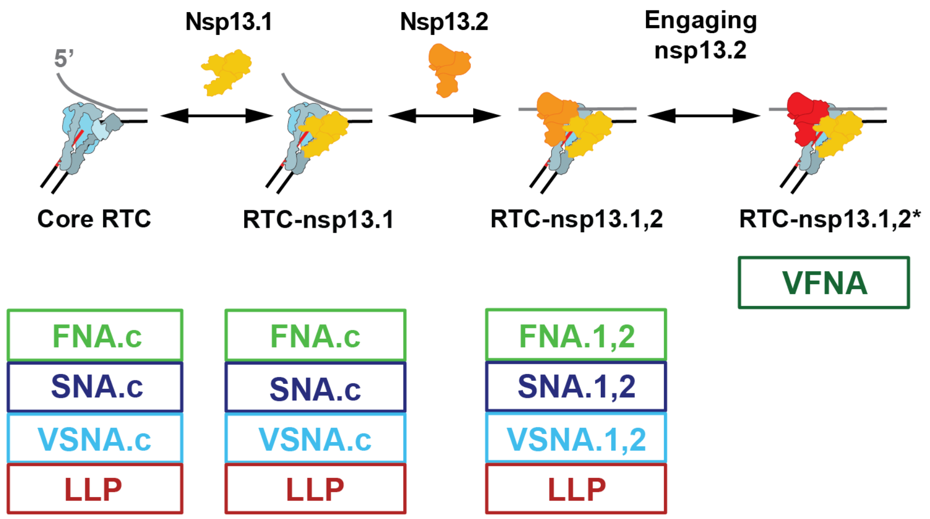

**Fig. S6. Model connecting the RTC assembly with elongation dynamics on dsRNA.** For the core RTC and each nsp13-bound non-engaged state, the fast (FNA), slow (SNA) and very slow nucleotide addition (VSNA) pathways can be entered and the long-lived pause (LLP). For the RTC with engaged nsp13 (RTC-nsp13.1,2*) the very fast nucleotide addition (VFNA) pathway is exclusively entered.

### Section S3: Building a mechanochemical model

Based on the empirical dwell-time distributions (**Fig. 3A**), we distinguished three NA pathways for the SARS-CoV-2 core RTC. Each pathway is characterized by a characteristic timescale and a probability of being entered at the start of each NA cycle (**Section S1**) (Bera et al. 2021). To build a mechanochemical model that captures the interrelated trends of single nucleotide probabilities and timescales, we started with a model where the NA cycle includes translocation, nucleotide binding, catalysis, and pyrophosphate (PPi) release (**Fig. S7A**). Each NA cycle starts with the RTC in the pre-translocated state. From here, the RTC moves reversibly to the post-translocated state where the NTP can bind and unbind. There can be many translocation and NTP binding cycles before the RNA product is extended through irreversible NTP catalysis and PPi release (**Fig. S7A**).

#### Section S3.1: Mechanochemistry of the non-bursting NA pathways

As unwinding the upstream dsRNA costs energy, the presence of the replication fork introduces a bias towards the pre-translocated state. Tension on the RNA destabilizes the replication fork and thus decreases this bias. Bera et al. 2021 found that the transition state for translocation is located close to the pre-translocated state and that the change in bias originates from the tension dependence of the backward translocation rate in the FNA pathway (**Fig. S7A**); we took this to be true for all NA pathways.

We observed that the average elongation velocity by the core RTC on dsRNA ($(3.6 \pm 0.1)$ nt/s) is much lower than on ssRNA ($(51.5 \pm 1.1)$ nt/s) (**Results**). This large velocity difference indicated that moving the replication fork is rate-limiting on dsRNA (Dulin, Vilfan et al. 2015, Dulin, Arnold et al. 2017). We therefore assumed NTP binding is in rapid equilibrium (Bera et al. 2021), used an effective nucleotide incorporation rate ($k_{\mathrm{irr}}$) and backward translocation rate ($k_{\mathrm{pre}}$) and ignored the time for PPi release (**Fig. S7A**). With these assumptions we arrived at a simplified mechanochemical model for the NA cycles in each non-bursting NA pathway.

#### Section S3.2: Connections between non-bursting NA pathways

With the mechanochemistry of a general NA cycle characterized, we turned to how the non-bursting NA pathways are connected. We observed that the FNA probability increased with RNA tension (**Fig. 4E**), while the SNA and VSNA probabilities decreased. Given the increasing bias towards the post-translocated state with tension, these trends suggested that each NA cycle starts in the FNA pre-translocated state and that the SNA and VSNA pathways are entered from there (**Fig. S7B**) (Bera et al. 2021).

If both the SNA and VSNA pathway would be directly entered from the FNA pre-translocated state, we would expect their relative probability to be independent of RNA tension. Instead, we observed that the SNA probability became relatively higher than the VSNA probability with tension (**Fig. 4E**). This suggested that the VSNA pathway is entered from the SNA pre-translocated state (**Fig. S7B**).

Since there were no clear trends in the VSNA characteristic timescale, we could not extract the backward translocation rate. We fixed it for all conditions (**Section S1.5**) and simplified the pathway to a single step with a tension independent probability $p_{\mathrm{irr}}^{\mathrm{VSNA}}$ (**Fig. S7B**).

#### Section S3.3: Mechanochemistry of the bursting NA pathway

The probability of very fast nucleotide addition bursts (VFNA) was independent of RNA tension, indicating that the stability of the RTC-nsp13.1,2* complex is not impacted by increasing RNA tension. Furthermore, the characteristic timescale of the VFNA pathway showed no significant trends versus RNA tension for saturating nsp13-helicase concentration (**Fig. 4DF**) and we concluded that it is limited by an RNA tension independent step.

#### Section S3.4: Mechanochemistry behind the allosteric effect

After deciphering the mechanochemistry and the connections between the NA pathways for the core RTC, we turned to the allosteric effect from the nsp13.2 association with the RTC. Here we assumed that the general reaction scheme is unaffected by binding of nsp13-helicase (**Fig. 4G**), but that the rates can change. Increasing the RNA tension had no significant effect on the FNA characteristic timescale and probability for RTC-nsp13.1,2 (**Fig. 4DF**), indicating that the backward translocation rate was negligible.

The SNA characteristic timescale for RTC-nsp13.1,2 is significantly lower than for the core RTC, indicating that RTC-nsp13.1,2 spends less time in the translocation cycles. Still, the SNA characteristic timescale decreases with increasing RNA tension (**Fig. 4D**), indicating that translocation cycles are not completely abolished, and the backward translocation rate is reduced but not negligible.

Furthermore, the VSNA probability decreased much more than the SNA and LLP probabilities, thus we let the probability for performing the irreversible step change to model the allosteric effect in our simplified description of the VSNA pathway.

#### Section S3.5: Connections for the long-lived pause to NA pathways

The LLP probability in general showed similar tension dependent trends as the VSNA probability (**Fig. 4EF**), suggesting that the LLP state is also entered from the pre-translocated state. For simplicity, we considered the LLP is entered from the pre-translocated state of each NA pathway with a constant relative rate (**Fig. 4G**).

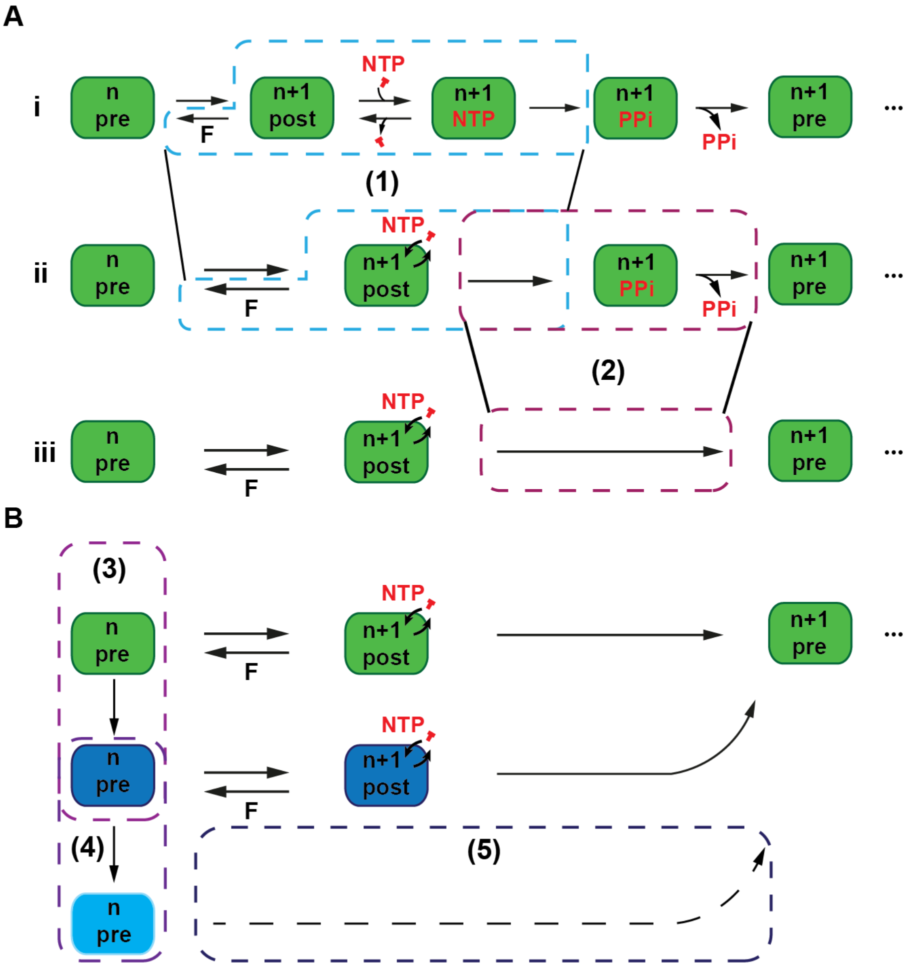

**Fig. S7. Simplification and connections made in the process of building the mechanochemical model for the SARS-CoV-2 RTC elongation dynamics on a dsRNA template. (A)** We started from a general mechanochemical model for the NA cycle (i) including translocation, nucleotide (NTP) binding, catalysis, and pyrophosphate (PPi) release (first line). Based on Bera et al. 2021, the tension (F) dependency is on the backward translocation rate. We determined that the translocation cycle is rate-limiting, therefore (1) (un)binding of NTPs is assumed in rapid equilibrium (ii) and (2) NTP catalysis and PPi release are considered as a single irreversible step (iii), which is fast compared to the characteristic timescale of the NA cycle. **(B)** From comparison of the NA pathway probabilities versus RNA tension for the core RTC we determined that (3) The SNA pathway (dark blue) is entered from the FNA pre-translocated state. (4) The VSNA pathway (sky blue) is entered from the SNA pre-translocated state. (5) Since we could not determine the mechanochemistry behind the VSNA characteristic timescale, we modelled the pathway with a probability to perform the irreversible step.

#### Section S3.6: Extracting parameters from the mechanochemical model

In **Sections S3.1-3.5** we arrived at a mechanochemical model for elongation dynamics by the SARS-CoV-2 RTC with or without nsp13-helicase shown in **Fig. 4G**. In this section we describe how the parameters in this model can be obtained from fits to the empirical dwell-time distributions.

In our mechanochemical model for elongation dynamics by the SARS-CoV-2 RTC (**Fig. 4G**), the single nucleotide timescale of each non-bursting NA pathway corresponds to the time for multiple cycles of forward and backward translocation before completing the irreversible step, starting from the pre-translocated state. We considered that translocation is not equilibrated in the FNA and SNA pathway (**Section S3.1**). Thus to obtain the first passage time distribution for completing the irreversible step in the $\mathrm{NA} \in \{\mathrm{FNA}, \mathrm{SNA}\}$ pathway by RTC complex $\mathrm{cpx} \in \{\mathrm{c}, 1, 2\}$, we summed over all possible paths with an arbitrary number of forward and backward translocations before the irreversible step of NTP binding and catalysis with rate $k_{\mathrm{irr}}^{\mathrm{NA}}$.

$$\Psi_{\mathrm{irr}}^{\mathrm{NA,cpx}}(s) = \sum_{n=0}^{\infty}\left(\frac{k_{\mathrm{post}}^{\mathrm{NA}}}{s + k_{\mathrm{post}}^{\mathrm{NA}} + k_{\mathrm{out}}^{\mathrm{NA}}}\frac{k_{\mathrm{pre}}^{\mathrm{NA,cpx}}(F)}{s + k_{\mathrm{pre}}^{\mathrm{NA,cpx}}(F) + k_{\mathrm{irr}}^{\mathrm{NA}}}\right)^{n}\frac{k_{\mathrm{post}}^{\mathrm{NA}}}{s + k_{\mathrm{post}}^{\mathrm{NA}} + k_{\mathrm{out}}^{\mathrm{NA}}}\frac{k_{\mathrm{irr}}^{\mathrm{NA}}}{s + k_{\mathrm{pre}}^{\mathrm{NA,cpx}}(F) + k_{\mathrm{irr}}^{\mathrm{NA}}} = \frac{k_{\mathrm{irr}}^{\mathrm{NA}} k_{\mathrm{post}}^{\mathrm{NA}}}{\left(s + k_{\mathrm{post}}^{\mathrm{NA}} + k_{\mathrm{out}}^{\mathrm{NA}}\right)\left(s + k_{\mathrm{pre}}^{\mathrm{NA,cpx}}(F) + k_{\mathrm{irr}}^{\mathrm{NA}}\right) - k_{\mathrm{post}}^{\mathrm{NA}} k_{\mathrm{pre}}^{\mathrm{NA,cpx}}(F)} \quad \text{(S25)}$$

With the rate to enter the slower pathways $k_{\mathrm{out}}^{\mathrm{FNA}} = k_{\mathrm{in}}^{\mathrm{SNA}} + k_{\mathrm{in}}^{\mathrm{LLP}}$ or $k_{\mathrm{out}}^{\mathrm{SNA}} = k_{\mathrm{in}}^{\mathrm{VSNA}} + k_{\mathrm{in}}^{\mathrm{LLP}}$ respectively.

From the first passage time distribution in Laplace space $\Psi_{\mathrm{irr}}^{\mathrm{NA,cpx}}(s)$, we obtained the probability and timescales of completing the irreversible step in pathway $\mathrm{NA} \in \{\mathrm{FNA}, \mathrm{SNA}\}$ as the zeroth and first moment of the distribution at $s$=0 respectively,

$$p_{\mathrm{irr}}^{\mathrm{NA,cpx}} = \Psi_{\mathrm{irr}}^{\mathrm{NA,cpx}}(0) = \frac{k_{\mathrm{irr}}^{\mathrm{NA}} k_{\mathrm{post}}^{\mathrm{NA}}}{\left(k_{\mathrm{post}}^{\mathrm{NA}} + k_{\mathrm{out}}^{\mathrm{NA}}\right)\left(k_{\mathrm{pre}}^{\mathrm{NA,cpx}}(F) + k_{\mathrm{irr}}^{\mathrm{NA}}\right) - k_{\mathrm{post}}^{\mathrm{NA}} k_{\mathrm{pre}}^{\mathrm{NA,cpx}}(F)};$$
$$\tau_{\mathrm{irr}}^{\mathrm{NA,cpx}} = -\partial_s \ln \Psi_{\mathrm{irr}}^{\mathrm{NA,cpx}}(0) = \frac{k_{\mathrm{post}}^{\mathrm{NA}} + k_{\mathrm{out}}^{\mathrm{NA}} + k_{\mathrm{pre}}^{\mathrm{NA,cpx}}(F) + k_{\mathrm{irr}}^{\mathrm{NA}}}{\left(k_{\mathrm{post}}^{\mathrm{NA}} + k_{\mathrm{out}}^{\mathrm{NA}}\right)\left(k_{\mathrm{pre}}^{\mathrm{NA,cpx}}(F) + k_{\mathrm{irr}}^{\mathrm{NA}}\right) - k_{\mathrm{post}}^{\mathrm{NA}} k_{\mathrm{pre}}^{\mathrm{NA,cpx}}(F)} \quad \text{(S26)}$$

With $k_{\mathrm{out}}^{\mathrm{FNA}} = k_{\mathrm{in}}^{\mathrm{SNA}} + k_{\mathrm{in}}^{\mathrm{LLP}}$; $k_{\mathrm{out}}^{\mathrm{SNA}} = k_{\mathrm{in}}^{\mathrm{VSNA}} + k_{\mathrm{in}}^{\mathrm{LLP}}$.

The kinetic rates for the RTC-nsp13.1 complex were assumed the same as for the core RTC, since the single nucleotide probabilities and timescales were also the same for these states (**Section S2.3**).

The single nucleotide probabilities for exiting through the FNA pathway by each RTC complex without engaged nsp13-helicase $\mathrm{cpx} \in \{\mathrm{c}, 1, 2\}$ were obtained as the probability to perform the irreversible step in the FNA pathway $p_{\mathrm{FNA}}^{\mathrm{cpx}} = p_{\mathrm{irr}}^{\mathrm{FNA,cpx}}$. The single nucleotide probabilities for performing the irreversible step of the slower $\mathrm{NA} \in \{\mathrm{SNA}, \mathrm{VSNA}\}$ pathways in each RTC complex $\mathrm{cpx} \in \{\mathrm{c}, 1, 2\}$ were obtained as the recursive probability for not exiting through the irreversible step in any of the faster pathways $p_{\mathrm{irr}}^{\mathrm{NA',cpx}}$ with $\mathrm{NA'} \in \{\mathrm{FNA}, \mathrm{SNA}\}$ times the splitting probability for entering the slower pathways $\frac{k_{\mathrm{in}}^{\mathrm{NA}}}{k_{\mathrm{out}}^{\mathrm{NA'}}}$ times the probability of completing the irreversible step in the pathway $p_{\mathrm{irr}}^{\mathrm{NA,cpx}}$ (**Fig. 4G**).

$$p_{\mathrm{FNA}}^{\mathrm{cpx}}(F) = p_{\mathrm{irr}}^{\mathrm{FNA,cpx}}(F);$$

$$p_{\mathrm{SNA}}^{\mathrm{cpx}}(F) = \left(1 - p_{\mathrm{irr}}^{\mathrm{FNA,cpx}}(F)\right)\frac{k_{\mathrm{in}}^{\mathrm{SNA}}}{k_{\mathrm{out}}^{\mathrm{FNA}}} p_{\mathrm{irr}}^{\mathrm{SNA,cpx}}(F); \tag{S27}$$

$$p_{\mathrm{VSNA}}^{\mathrm{cpx}}(F) = \left(1 - p_{\mathrm{irr}}^{\mathrm{FNA,cpx}}(F)\right)\frac{k_{\mathrm{in}}^{\mathrm{SNA}}}{k_{\mathrm{out}}^{\mathrm{FNA}}}\left(1 - p_{\mathrm{irr}}^{\mathrm{SNA,cpx}}(F)\right)\frac{k_{\mathrm{in}}^{\mathrm{VSNA}}}{k_{\mathrm{out}}^{\mathrm{SNA}}} p_{\mathrm{irr}}^{\mathrm{VSNA,cpx}};$$

$$p_{\mathrm{LLP}}^{\mathrm{cpx}}(F) = 1 - p_{\mathrm{FNA}}^{\mathrm{cpx}}(F) - p_{\mathrm{SNA}}^{\mathrm{cpx}}(F) - p_{\mathrm{VSNA}}^{\mathrm{cpx}}(F)$$

With $k_{\mathrm{out}}^{\mathrm{FNA}} = k_{\mathrm{in}}^{\mathrm{SNA}} + k_{\mathrm{in}}^{\mathrm{LLP}}$; $k_{\mathrm{out}}^{\mathrm{SNA}} = k_{\mathrm{in}}^{\mathrm{VSNA}} + k_{\mathrm{in}}^{\mathrm{LLP}}$. The relation for the single nucleotide probability for entering the long-lived pause $p_{\mathrm{LLP}}^{\mathrm{cpx}}$ ensures that the single nucleotide probabilities of all pathways for each RTC complex add up to 1. Note that with keeping $k_{\mathrm{in}}^{\mathrm{LLP}}$ constant for the different non-engaged RTC-nsp13 complexes, we did not enforce that the long-lived pause probability is constant for all non-engaged RTC-nsp13 complexes, like we did in the fit of the RTC assembly – Elongation dynamics model (**Section S2.3**), but with the global fit of the mechanochemical model we found similar values ($p_{\mathrm{LLP}}^{\mathrm{c}} \approx p_{\mathrm{LLP}}^{2}$) (**Table S3**).

Translocation acts over the distance from the conversion of one base pair dsRNA to ssRNA $a = \delta x_{\mathrm{ss}} - \delta x_{\mathrm{ds}} \approx 0.2$ nm (Dulin et al. 2015) and we determined there is no tension dependence of the forward translocation rate (**Section S3.1**). Therefore, the (negative) RNA tension ($F$) dependence of the backward translocation rate could be described as the Arrhenius law with distance $a$

$$k_{\mathrm{pre}}^{\mathrm{NA,cpx}}(F) = k_{\mathrm{pre}}^{\mathrm{NA,cpx}}(0)e^{-aF/k_{\mathrm{B}}T}, \qquad \mathrm{NA} \in \{\mathrm{FNA, SNA}\}, \qquad \mathrm{cpx} \in \{\mathrm{c}, 1, 2\} \tag{S28}$$

The allosteric effect of nsp13.2 binding to the RTC was modelled as an increase in the free-energy barrier $\Delta G_{\mathrm{NA}}$ in units of $\mathrm{k_BT}$ for backward translocation rate $k_{\mathrm{pre,NA.2}}$ in each NA pathway (**Section S3.4**).

$$k_{\mathrm{pre}}^{\mathrm{NA,2}}(F) = k_{\mathrm{pre}}^{\mathrm{NA,c}}(F)e^{\Delta G_{\mathrm{NA}}}, \qquad \mathrm{NA} \in \{\mathrm{FNA, SNA}\} \tag{S29}$$

The other rates in the mechanochemical model were unchanged when nsp13.2 is in complex with the RTC (**Fig. 4G**).

Since the VSNA characteristic timescale was fixed for all conditions (**Section S1.5**), we modelled the pathway as a single irreversible step with probability $p_{\mathrm{irr}}^{\mathrm{VSNA,cpx}}$ for each RTC complex cpx $\in \{\mathrm{c}, 1, 2\}$ (**Fig. 4G**). For the core RTC (cpx =c) and the RTC-nsp13.1 complex (cpx =1) the probabilities for the irreversible step are the same $p_{\mathrm{irr}}^{\mathrm{VSNA,c}} = p_{\mathrm{irr}}^{\mathrm{VSNA,1}}$, but for the RTC-nsp13.1,2 complex the probability $p_{\mathrm{irr}}^{\mathrm{VSNA,2}}$ is decreased, which reflects the substantial decrease in the VSNA single nucleotide probability (**Table S3**) and the total VSNA probability at saturating nsp13-helicase concentration (**Fig. 4F**).

As the long-lived pause probability was not much decreases at saturating nsp13-helicase concentration, but showed a similar trend with RNA tension as the VSNA probability (**Fig. 4EF**), We simply modelled a direct entry of the long-lived pause from the pre-translocated state of the FNA and SNA pathway with relative rate $k_{\mathrm{in}}^{\mathrm{LLP}}$ and from the VSNA pre-translocated state with probability $(1 - p_{\mathrm{irr}}^{\mathrm{VSNA,cpx}})$. The decreased value of $p_{\mathrm{irr}}^{\mathrm{VSNA,2}}$ compensated for the decrease in entry of the long-lived pause from the FNA and SNA pre-translocated state, resulting in a stable long-lived pause probability ($p_{\mathrm{LLP}}^{\mathrm{c}} \approx p_{\mathrm{LLP}}^{2}$) (**Table S3**).

#### Section S3.7: Global fit of the mechanochemical model

To perform a global fit of the mechanochemical model to the empirical dwell-time distributions, we substituted the RNA tension dependencies (**Eq. S28**) and allosteric effects (**Eq. S29**) into the expressions for the single nucleotide probabilities and timescales versus RNA tension (**Eq. S26** with **Eq. S27**) in the RTC assembly-Elongation dynamics model (**Eq. S21-S23**), to obtain a pdf curve for each dwell-time distributions for each condition measured.

A global fit of the mechanochemical model was performed on the dwell-time distributions of the nsp13-helicase concentration dependency dataset at RNA tension $F$ = 20 pN and RNA tension dependency datasets without nsp13-helicase and at saturating nsp13-helicase concentration (**Fig. S14**). The free parameters in the mechanochemical model were $k_{\mathrm{in}}^{\mathrm{SNA}}, k_{\mathrm{in}}^{\mathrm{VSNA}}, k_{\mathrm{in}}^{\mathrm{LLP}}, k_{\mathrm{post}}^{\mathrm{FNA}}, k_{\mathrm{post}}^{\mathrm{SNA}}, k_{\mathrm{pre}}^{\mathrm{FNA.c}}(0), k_{\mathrm{pre}}^{\mathrm{FNA.2}}(0), k_{\mathrm{pre}}^{\mathrm{SNA.c}}(0), k_{\mathrm{pre}}^{\mathrm{SNA.2}}(0),$ $k_{\mathrm{irr}}^{\mathrm{FNA}}, k_{\mathrm{irr}}^{\mathrm{SNA}}, p_{\mathrm{irr}}^{\mathrm{VSNA.c}}, p_{\mathrm{irr}}^{\mathrm{VSNA.2}}$ with which the single nucleotide probabilities and timescales could be obtained (**Eq. S26-S29**). The substitution into the RTC assembly-Elongation dynamics model added the free-energy differences between the states in the RTC assembly model $\Delta G_{\mathrm{c}\rightarrow 1}$, $\Delta G_{1\rightarrow 2}$, $\Delta G_{2\rightarrow 2^*}$ (**Eq. S21-24**).

The global fit of the mechanochemical model on all datasets successfully captured the trends in the probabilities and characteristic timescales of all the pathways (**Fig. 3CD** and **Fig. 4C-F**). As expected, we obtained that for RTC-nsp13.1,2 the FNA backward translocation rate was negligible compared to the FNA forward translocation rate (**Table S3**). The values of the probability to perform the irreversible step in the VSNA pathway for the RTC-nsp13.1,2 complex $p_{\mathrm{irr}}^{\mathrm{VSNA.2}}$ were, however, much lower than for the core RTC ($p_{\mathrm{irr}}^{\mathrm{VSNA.c}}$) (**Table S3**), while we expected it would have a similar or higher value than $p_{\mathrm{irr}}^{\mathrm{VSNA.c}}$ based on the allosteric effect obtained for the FNA and SNA pathway. We did not seek to find an explanation for this, because the VSNA probability was much lower than the FNA and SNA probability and the position of the corresponding bump was hard to determine for saturating concentration of nsp13-helicase (**Section S1.5**).

#### Section S3.8: Description of the mechanochemical model

Comparing the kinetic rates extracted from the global fit for the core RTC and RTC-nsp13.1,2 for each nucleotide addition pathway (**Table S3**). We identify how the trends observed on the dwell-time level arise from the underlying mechanochemistry. Since the core RTC and RTC-nsp13.1 have indiscernible elongation dynamics (**Fig. S6**), we refer to both as core RTC.

Starting with the FNA pathway for the core RTC, we fit out a forward translocation rate $k_{\mathrm{post}}^{\mathrm{FNA}}$ ~3-fold smaller than the backward translocation rate $k_{\mathrm{pre}}^{\mathrm{FNA}}$ at 20 pN RNA tension, resulting in a more populated pre-translocated state than the post-translocated state. However, since the entrance rates for the SNA pathway $k_{\mathrm{in}}^{\mathrm{SNA}}$ and the LLP state $k_{\mathrm{in}}^{\mathrm{LLP}}$ are ~30- and ~10000-fold smaller than the FNA forward translocation rate $k_{\mathrm{post}}^{\mathrm{FNA}}$, the FNA pathway is the most populated pathway. For RTC-nsp13.1,2 we obtained a ~10-fold decrease of the FNA backward translocation rate $k_{\mathrm{pre}}^{\mathrm{FNA}}$ resulting from nsp13.2 allostery (**Table S3**). As a consequence, the FNA characteristic timescale is shorter and the FNA probability is increased relative to the other non-bursting NA pathways for the RTC-nsp13.1,2 complex (**Fig. 4C-F**).

Concerning the SNA pathway, we observed that the forward translocation rate $k_{\mathrm{post}}^{\mathrm{SNA}}$ is ~100-fold slower than the backward translocation rate $k_{\mathrm{pre}}^{\mathrm{SNA}}$ for the core RTC and 20 pN RNA tension (**Table S3**). For the core RTC, the SNA pre-translocated state is therefore much more populated than the SNA post-translocated state. Additionally, since the entrance rates of the VSNA pathway $k_{\mathrm{in}}^{\mathrm{VSNA}}$ and the LLP state $k_{\mathrm{in}}^{\mathrm{LLP}}$ are ~1000- and ~10000-fold smaller than the SNA forward translocation rate $k_{\mathrm{post}}^{\mathrm{SNA}}$ (**Table S3**), the probability to exit through the SNA pathway is much larger than the probability to enter either the VSNA pathway or the LLP state (**Fig. 4CE**). For RTC-nsp13.1,2, the SNA backward translocation rate $k_{\mathrm{pre}}^{\mathrm{SNA}}$ is reduced with respect to the core RTC, but $k_{\mathrm{pre}}^{\mathrm{SNA}}$ is still ~50-fold larger than the SNA forward translocation rate $k_{\mathrm{post}}^{\mathrm{SNA}}$, which remains unchanged (**Table S3**). As a result, the SNA characteristic timescale still shows a significant tension dependency for RTC-nsp13.1,2 (**Fig. 4D**) and the probabilities to enter the VSNA pathway and the LLP state decrease with increasing RNA tension (**Fig. 4F**).

The probability for the irreversible step of the VSNA pathway starting from the VSNA pre-translocated state $p_{\mathrm{irr}}^{\mathrm{VSNA}}$ is ~4 times decreases from the core RTC to the RTC-nsp13.1,2 complex, while the rate into the LLP state stayed constant $k_{\mathrm{in}}^{\mathrm{LLP}}$ (**Table S3**), resulting in a much lower overall VSNA probability compared to the other pathways (**Fig. 4F**).

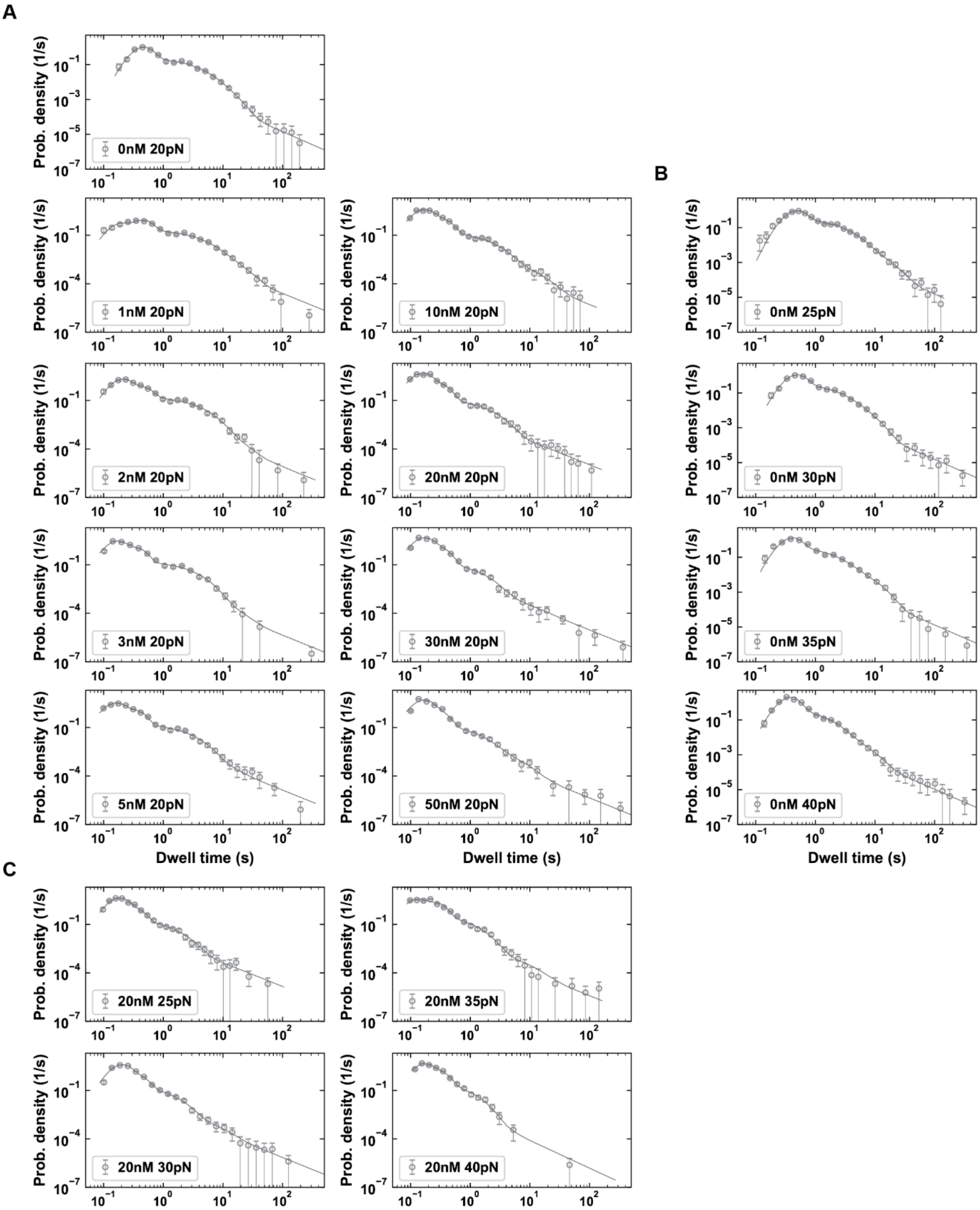

**Fig. S8. Dwell-time fits with all parameters free for RTC elongation dynamics on a dsRNA template with nsp13-helicase**. **(A)** Fit curves on the dwell-time distributions versus nsp13-helicase concentration at 20 pN RNA tension. **(B, C)** Fit curves on the dwell-time distributions versus RNA tension without nsp13-helicase (B) or at 20 nM nsp13-helicase (C). The peaks and shoulders in the distributions are well-fitted for each condition. The dwell-time distributions are divided into 30 bins, zero value bins are removed and the remaining bins are centered again. The circles and error bars represent the mean and standard deviation from the mean for every bin obtained from 100 bootstrap fits.

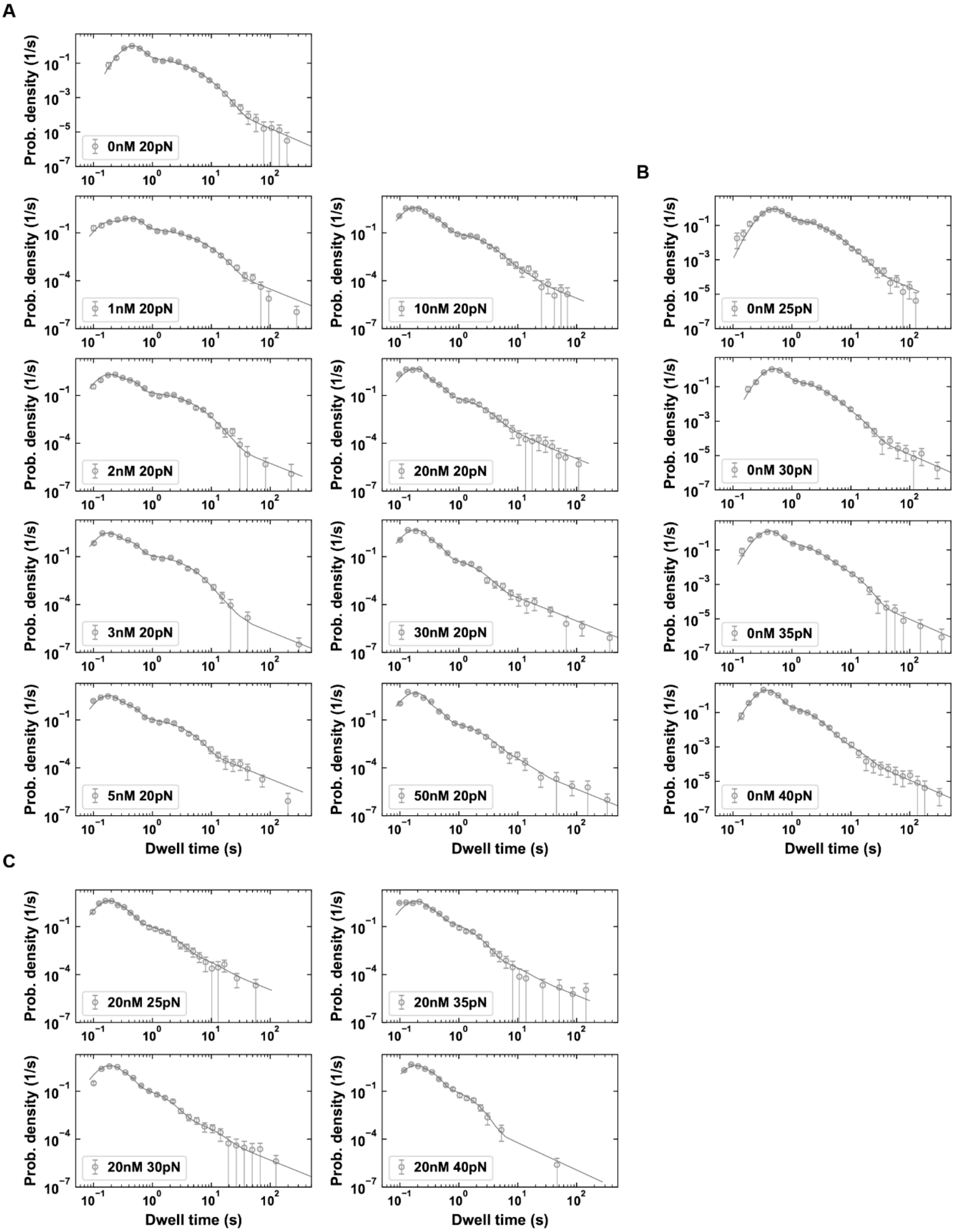

**Fig. S9. Dwell-time fits with the VFNA and VSNA characteristic timescale fixed for RTC elongation dynamics on a dsRNA template with nsp13-helicase**. **(A)** Fit curves on the dwell-time distributions versus nsp13-helicase concentration at 20 pN RNA tension. **(B, C)** Fit curves on the dwell-time distributions versus RNA tension without nsp13-helicase (B) or at 20 nM nsp13-helicase (C). The peaks and shoulders in the distributions are well-fitted for every condition. The VFNA and VSNA characteristic timescale were fixed to 0.2 s and 4.8 s respectively for all conditions**.** Additionally, the FNA characteristic timescale was fixed to ~0.03 s and the VFNA probability to 0.56 for the fits to the dwell-time distributions at 20 nM nsp13-helicase with varying RNA tension. The dwell-time distributions are divided into 30 bins, zero value bins are removed and the remaining bins are centered again. The circles and error bars represent the mean and standard deviation from the mean for every bin obtained from 100 bootstrap fits.

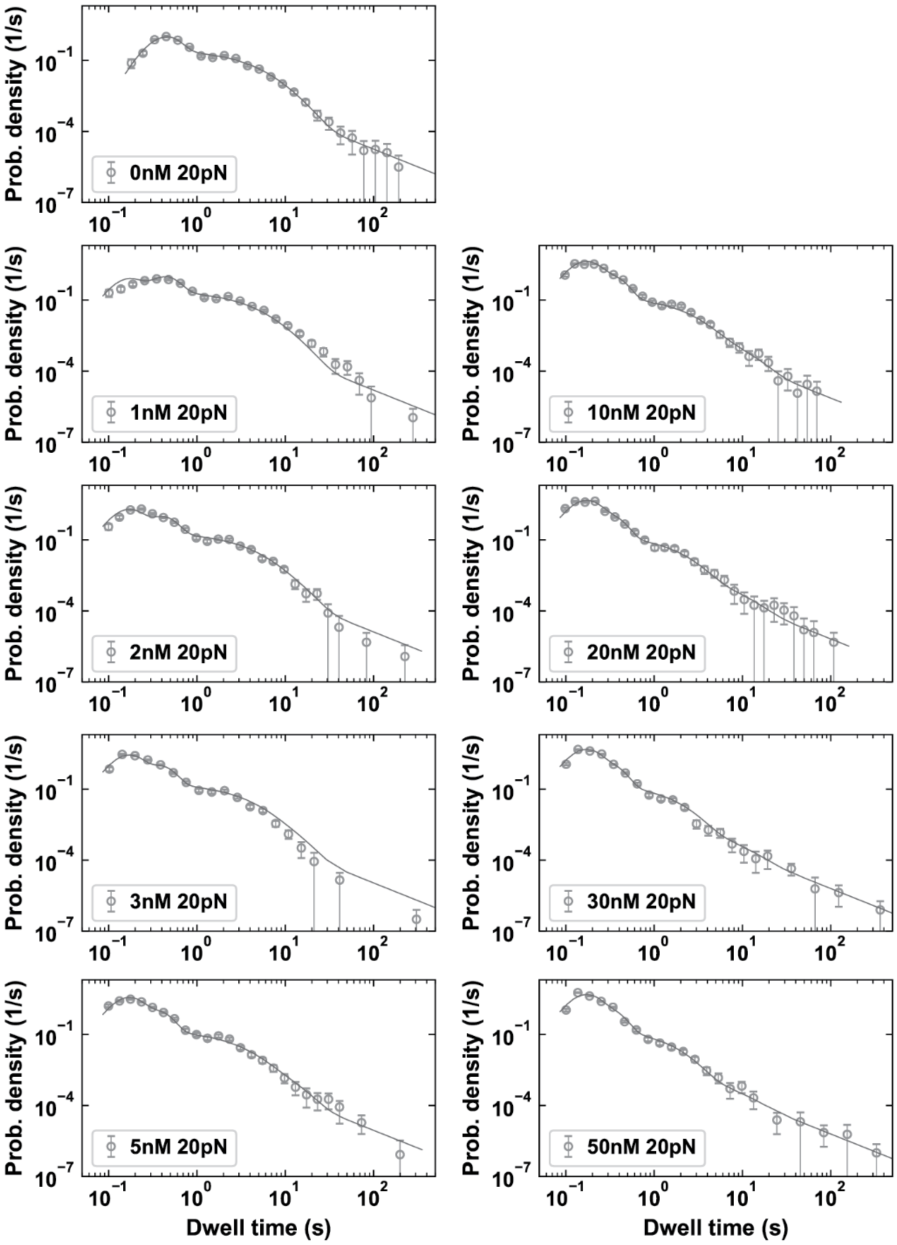

**Fig. S10. Global fit of RTC Assembly-Elongation dynamics model with the single nucleotide parameters free for the RTC-nsp13.1,2 complex.** The peaks and shoulders in the distributions are well-fitted for every condition, including the peak for the condition without nsp13-helicase. The VFNA and VSNA characteristic timescale were fixed to 0.2 s and 4.8 s respectively for all conditions. The dwell-time distributions are divided into 30 bins, zero value bins are removed and the remaining bins are centered again. The circles and error bars represent the mean and standard deviation from the mean for every bin obtained from 100 bootstrap fits.

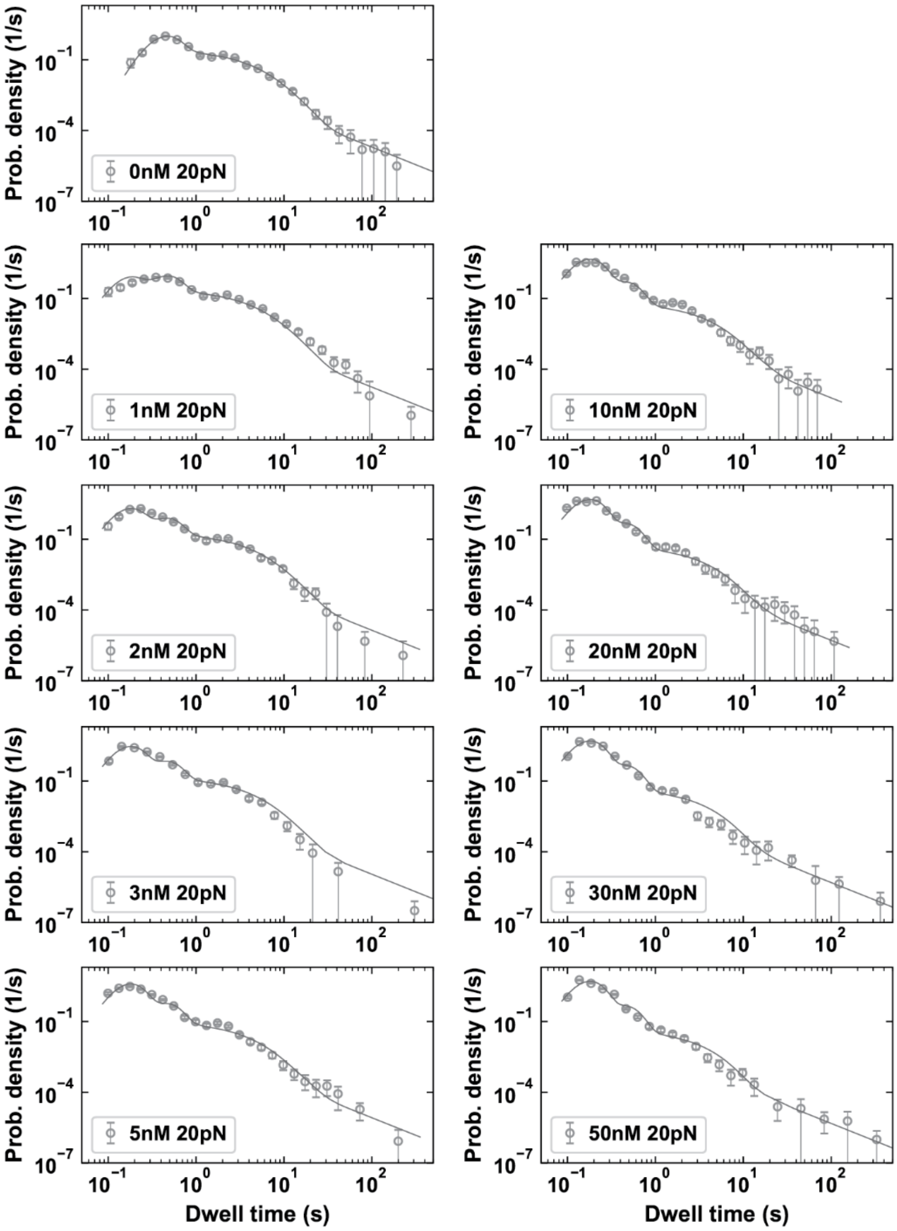

**Fig. S11. Global fit of RTC Assembly-Elongation dynamics model with the single nucleotide probabilities free for the RTC-nsp13.1,2 complex.** The peaks and shoulders in the distributions are well-fitted for low concentrations of nsp13-helicase, but for high concentrations the fits to the second peak and shoulders are significantly off. The VFNA and VSNA characteristic timescale were fixed to 0.2 s and 4.8 s respectively for all conditions. The dwell-time distributions are divided into 30 bins, zero value bins are removed and the remaining bins are centered again. The circles and error bars represent the mean and standard deviation from the mean for every bin obtained from 100 bootstrap fits.

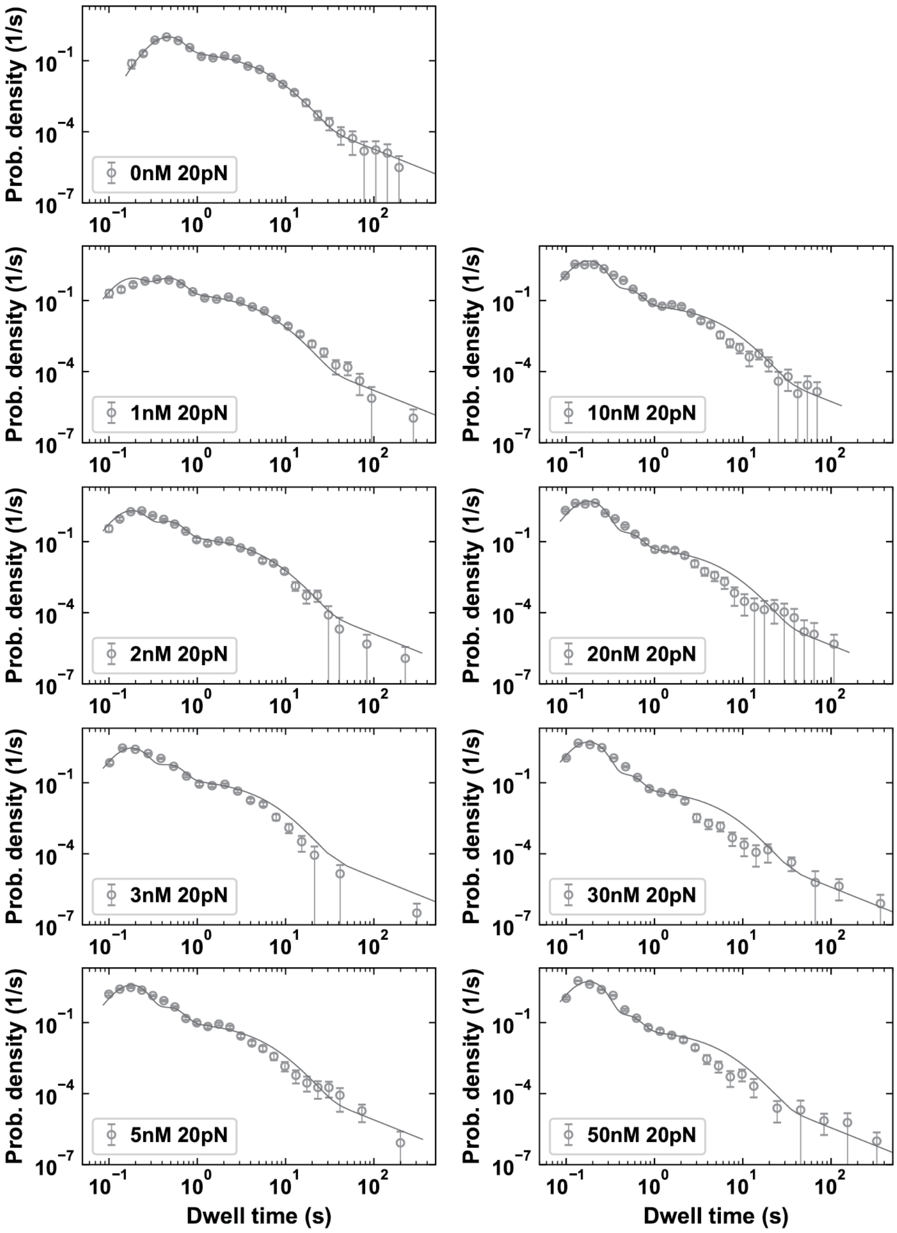

**Fig. S12. Global fit of RTC Assembly-Elongation dynamics model with the single nucleotide probabilities and timescales fixed to the core RTC values.** The single nucleotide parameters in the core RTC and nsp13-bound RTC complexes (RTC-nsp13.1 and RTC-nsp13.1,2) were fixed to the values obtained from the fit to the dwell-times at zero nsp13-helicase concentration and 20 pN RNA tension. The goodness-of-fit of the peaks and shoulders decreases with higher concentrations of nsp13-helicase. This confirms there is a significant change in the single nucleotide parameters for the nsp13.2 non-engaged RTC complex. The dwell-time distributions are divided into 30 bins, zero value bins are removed and the remaining bins are centered again. The circles and error bars represent the mean and standard deviation from the mean for every bin obtained from 100 bootstrap fits.

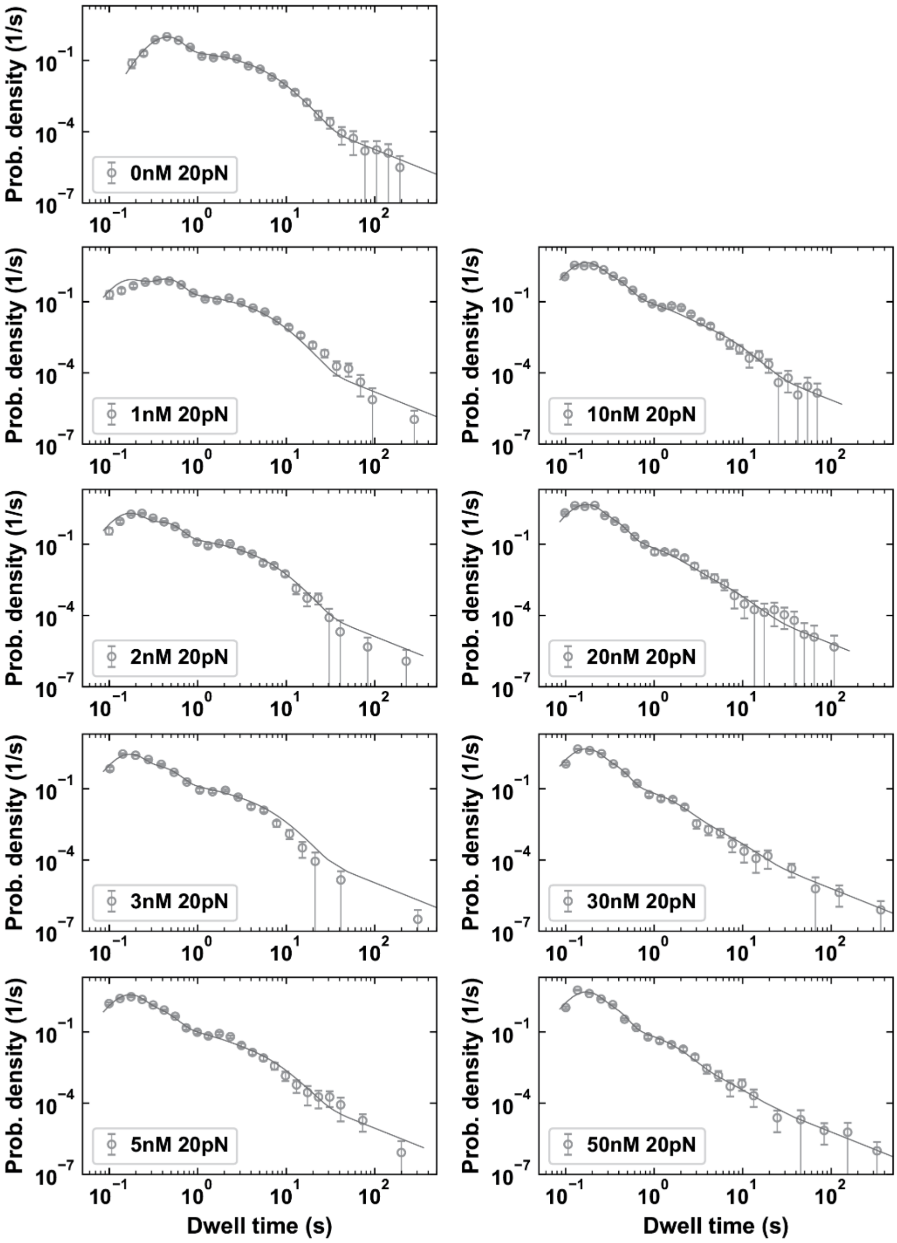

**Fig. S13. Check on timescale averaging with fitted parameters from the RTC Assembly - Elongation dynamics model versus nsp13-helicase concentration.** The elongation dynamics distributions for the different RTC complexes combined were calculated using the parameter values obtained with a global fit of the RTC Assembly - Elongation dynamics model with the effective characteristic timescale and probabilities for each pathway with the single nucleotide probabilities and timescales free for the RTC-nsp13.1,2 complex to the dwell-time distributions versus nsp13-helicase concentration. The pdf curves are mostly within one standard deviation of the dwell-time distributions except for the first bump at intermediate concentrations.

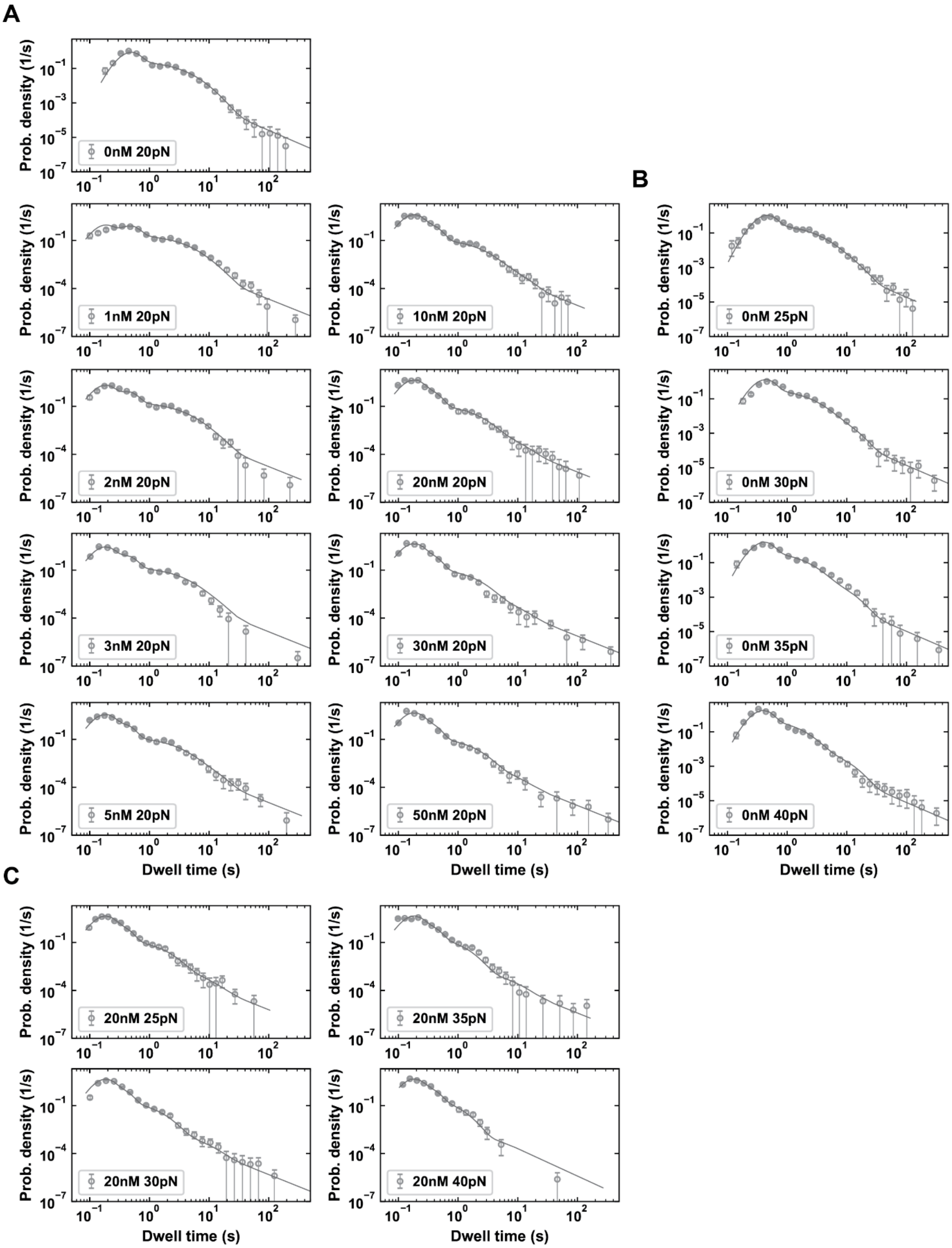

**Fig. S14. Global fit of the combination of the RTC assembly, elongation dynamics and the mechanochemical model for RTC elongation on a dsRNA template in presence and absence of nsp13-helicase. (A)** Fitted pdf curves on the dwell-time distributions versus nsp13-helicase concentration at constant RNA tension (20 pN). **(B, C)** Fitted pdf curves on the dwell-time distributions versus RNA tension at zero nsp13-helicase concentration (B) and at saturating nsp13-helicase concentration (20 nM) (C). The fits capture the trends in the dwell-time distributions for each condition. The dwell-time distributions are divided into 30 bins, zero value bins are removed and the remaining bins are centered again. The circles and error bars represent the mean and standard deviation from the mean for every bin obtained from 100 bootstrap fits.

| Model | Dataset | [nsp13] (nM) | $F$ (pN) | $t_{cut}$ (1/s) | Value type | $\chi^2$ | $\Delta G_1 (k_B T)$ | $\Delta G_{1,2} (k_B T)$ | $\Delta G_{1,2^*} (k_B T)$ |
|---|---|---|---|---|---|---|---|---|---|
| Assembly model 1; RTC-nsp13.1* | [nsp13] dependency | 0-50 | 20 | 0.08 | **Best fit values** | 58932 | 2.9 | - | -1.5 |
| Assembly model 2; RTC-nsp13.1,2* | | | | | **Best fit values** | 21659 | 1.5 | 1.4 | -1.0 |
| | | | | | **Bounds** | | [-3, 3] | [-3, 3] | [-3, 3] |

**Table S1. Parameters for RTC assembly model fits to the VFNA probabilities versus nsp13-helicase concentration.** The fractional occupancy of the nsp13-engaged RTC complex from both models was fitted (RTC-nsp13.1* or RTC-nsp13.1,2*) to the VFNA probability versus nsp13-helicase concentration ([nsp13]). The Chi-squared ($\chi^2$) value for the Assembly model 2 fit was about half the value for the Assembly model 1 fit, showing that Assembly model 2 gives a better fit.

| Model | Conditions | [nsp13](nM) | $F$(pN) | $t_{\text{cut}}(s)$ | Fit conditions | Value type | $LL$ | BIC | $\Delta G_1$ ($k_BT$) | $\Delta G_{1,2}$ ($k_BT$) | $\Delta G_{1,2^*}$ ($k_BT$) |
|---|---|---|---|---|---|---|---|---|---|---|---|
| RTC Assembly - Elongation dynamics | [nsp13] dependency | 0-50 | 20 | 0.08 | RTC-nsp13.1,2 single nt probabilities and timescales free | **Best fit values** | -18372 | 36885 | 1.7 | 0.81 | -0.61 |
| | | | | | | **Mean values** | | | 1.74 | 0.87 | -0.69 |
| | | | | | | **Std values** | | | 0.16 | 0.09 | 0.16 |
| | | | | | RTC-nsp13.1,2 single nt probabilities free | **Best fit values** | -19223 | 38534 | 2.10 | 0.99 | -1.22 |
| | | | | | All single nt parameters fixed | **Best fit values** | -20291 | 40614 | 1.2 | -1.57 | 2.01 |
| | | | | | | **Bounds** | | | [-3, 3] | [-3, 3] | [-3, 3] |

| Value type | $p_{\text{FNA.c}}(F_0)$ | $p_{\text{FNA.1,2}}(F_0)$ | $p_{\text{SNA.c}}(F_0)$ | $p_{\text{SNA.1,2}}(F_0)$ | $p_{\text{VSNA.c}}(F_0)$ | $p_{\text{VSNA.1,2}}(F_0)$ | $p_{\text{LLP}}(F_0)$ | $\tau_{\text{VFNA}}(F_0)(s)$ | $\tau_{\text{FNA.c}}(F_0)(s)$ | $\tau_{\text{FNA.1,2}}(F_0)(s)$ | $\tau_{\text{SNA.c}}(F_0)(s)$ | $\tau_{\text{SNA.1,2}}(F_0)(s)$ | $\tau_{\text{VSNA.c}}(F_0)(s)$ | $\tau_{\text{VSNA.1,2}}(F_0)(s)$ |
|---|---|---|---|---|---|---|---|---|---|---|---|---|---|---|
| **Best fit values** | 0.89 | 0.96 | 0.085 | 0.036 | 0.019 | 0.0006 | 0.0030 | 0.02 | 0.048 | 0.033 | 1.6 | 0.59 | 4.40 | 4.79 |
| **Mean values** | 0.89 | 0.96 | 0.085 | 0.036 | 0.019 | 0.0008 | 0.0033 | 0.02 | 0.048 | 0.034 | 1.56 | 0.56 | 4.41 | 4.78 |
| **Std values** | 0.00 | 0.00 | 0.003 | 0.002 | 0.002 | 0.0007 | 0.0004 | 0 | 0.001 | 0.001 | 0.04 | 0.04 | 0.04 | 0.01 |
| **Best fit values** | 0.90 | 0.96 | 0.081 | 0.033 | 0.017 | 0 | 0.0033 | 0.02* | 0.05* | 0.05* | 1.64* | 1.64* | 4.37* | 4.37* |
| **Best fit values** | 0.9* | 0.9* | 0.077* | 0.077* | 0.019* | 0.019* | 0.0031* | 0.02* | 0.05* | 0.05* | 1.64* | 1.64* | 4.37* | 4.37* |
| **Bounds** | [0, 1] | [0, 1] | [0, 0.5] | [0, 0.1] | [0, 0.1] | [0, 0.1] | [0, 0.1] | [0.008, 0.1] | [0.02, 0.1] | [0.02, 0.1] | [0.05, 5] | [0.05, 5] | | |

**Table S2. Parameter values for the RTC Assembly - Elongation dynamics model.** Parameters for the best fit (row 1) and the mean and standard deviation of 100 bootstrap fits (row 2 and 3) are shown for the fit with the single nucleotide probabilities and timescales free for the RTC-nsp13.1,2 complex, with only the VSNA characteristic timescale fixed. (row 4) Parameters of the best fit with the single nucleotide probabilities for the RTC-nsp13.1,2 complex free and the single nucleotide timescales fixed to the values for the core RTC. (row 5) Parameters of the best fit keeping all the single nucleotide probabilities and timescales for the RTC-nsp13.1,2 complex fixed to the values for the core RTC. (row 6) The boundaries on the fitted parameters. *Parameters that are fixed during the fit.

| Model | Conditions | [nsp13](nM) | $F$(pN) | $t_{cut}(s)$ | Value type | $LL$ | BIC | $\Delta G_1$ ($k_BT$) | $\Delta G_{1,2}$ ($k_BT$) | $\Delta G_{1,2^*}$ ($k_BT$) |
|---|---|---|---|---|---|---|---|---|---|---|
| RTC assembly, elongation dynamics and mechanochemistry | [nsp13] dependency; tension dependency at [nsp13]=0; at [nsp13]=20 nM | 0-50 | 20-40 | 0.08 | **Best fit values** | -47861 | 95907 | 1.38 | 0.90 | -0.72 |
| | | | | | **Mean values** | | | 1.56 | 0.77 | -0.72 |
| | | | | | **Std values** | | | 0.23 | 0.17 | 0.03 |
| | | | | | **Bounds** | - | - | [-3, 3] | [-3, 3] | [-3, 3] |

| Value type | $k_{in,SNA}$ $(1/s)$ | $k_{in,VSNA}$ $(1/s)$ | $k_{in,LLP}$ $(1/s)$ | $k_{post,FNA}$ $(1/s)$ | $k_{post,SNA}$ $(1/s)$ | $k_{pre,FNA.c}$ $(1/s)$ | $k_{pre,FNA.1,2}$ $(1/s)$ | $k_{pre,SNA.c}$ $(1/s)$ | $k_{pre,SNA.1,2}$ $(1/s)$ | $k_{irr,FNA}$ $(1/s)$ | $k_{irr,SNA}$ $(1/s)$ | $p_{irr,VSNA.c}$ | $p_{irr,VSNA.1,2}$ |
|---|---|---|---|---|---|---|---|---|---|---|---|---|---|
| **Best fit values** | 3.9 | 0.13 | 0.021 | 97 | 145 | 228 | 20 | 15310 | 5912 | 40.1 | 20.2 | 1.00 | 0.30 |
| **Mean values** | 3.9 | 0.14 | 0.015 | 104 | 121 | 252 | 32 | 12858 | 4991 | 39.9 | 20.0 | 0.92 | 0.25 |
| **Std values** | 0.1 | 0.01 | 0.006 | 5 | 27 | 22 | 9 | 2740 | 1101 | 0.4 | 0.5 | 0.05 | 0.15 |
| **Bounds** | [0, 10] | [0, 10] | [0, 10] | [1, Inf] | [1, Inf] | [1, Inf] | [1, Inf] | [1, Inf] | [1, Inf] | [0, 100] | [0, 100] | [0, 1] | [0, 1] |

| Value Type | $p_{FNA.c}$ $(F_0)$ | $p_{FNA.1.2}$ $(F_0)$ | $p_{SNA.c}$ $(F_0)$ | $p_{SNA.1.2}$ $(F_0)$ | $p_{VSNA.c}$ $(F_0)$ | $p_{VSNA.1.2}$ $(F_0)$ | $p_{LLP.c}$ $(F_0)$ | $p_{LLP.1.2}(F_0)$ | $\tau_{VFNA}(F_0)(s)$ | $\tau_{FNA.c}(F_0)(s)$ | $\tau_{FNA.1.2}(F_0)(s)$ | $\tau_{SNA.c}(F_0)(s)$ | $\tau_{SNA.1.2}(F_0)(s)$ | $\tau_{VSNA.c}(F_0)(s)$ | $\tau_{VSNA.1.2}(F_0)(s)$ |
|---|---|---|---|---|---|---|---|---|---|---|---|---|---|---|---|
| **Best fit values** | 0.89 | 0.95 | 0.086 | 0.041 | 0.022 | 0.0012 | 0.0043 | 0.0038 | 0.02 | 0.052 | 0.036 | 1.57 | 0.74 | 4.3 | 4.8 |
| **Mean values** | 0.88 | 0.95 | 0.087 | 0.042 | 0.022 | 0.0011 | 0.0049 | 0.0040 | 0.02 | 0.052 | 0.037 | 1.59 | 0.75 | 4.4 | 4.8 |
| **Std values** | 0.00 | 0.00 | 0.002 | 0.001 | 0.001 | 0.0007 | 0.0006 | 0.0007 | 0.00 | 0.000 | 0.000 | 0.04 | 0.04 | 0.0 | 0.0 |

**Table S3. Global fit parameters for the RTC assembly, elongation dynamics and mechanochemical model**. Values for the best fit, the mean and standard deviation from 100 bootstrap fits and the boundaries on the fitted parameters are shown. (Top) Conditions for the fitted data, goodness-of-fit parameters (BIC and *LL*) and parameters for the free-energy landscape. (Middle) Microscopic parameters of the mechanochemical model. (Bottom) The parameters for the RTC Assembly - Elongation dynamics model at RNA tension $F_0$ = 20 pN calculated from the microscopic parameters.
